# High-throughput robotic isolation of human iPS cell clones reveals frequent homozygous induction of identical genetic manipulations by CRISPR-Cas9

**DOI:** 10.1101/2024.09.18.613641

**Authors:** Gou Takahashi, Minato Maeda, Kayoko Shinozaki, Gakuro Harada, Saburo Ito, Yuichiro Miyaoka

**Author notes:** Correspondence (Y.M.), X account: @YuichiroMiyaoka. Current address: Department of Animal Science, Tokyo University of Agriculture, Atsugi, Kanagawa 243-0034, Japan (Gou Takahashi).

## Abstract

Genome editing in human iPS cells is a powerful approach in regenerative medicine. CRISPR-Cas9 is the most common genome editing tool, but it often induces byproduct insertions and deletions in addition to the desired edits. Therefore, genome editing of iPS cells produces diverse genotypes. Existing assays mostly analyze genome editing results in cell populations, but not in single cells. However, systematic profiling of genome editing outcomes in single iPS cells was lacking. In this study, we developed a method for high-throughput iPS cell clone isolation based on the precise robotic picking of cell clumps derived from single cells grown in extracellular matrices. We analyzed over 1,000 genome-edited iPS cell clones and found that homozygous editing was much more frequent than heterozygous editing. We also observed frequent homozygous induction of identical genetic manipulations, including insertions and deletions. Our new cloning method and findings will facilitate the application of genome editing to human iPS cells.

## INTRODUCTION

Genome editing has revolutionized our ability to study and cure genetic disorders (Gaj et al., 2013). In particular, genome editing in induced pluripotent stem (iPS) cells allows the development of human cell-based isogenic disease models and potential cell therapies (Hockemeyer and Jaenisch, 2016). Clustered regularly interspaced short palindromic repeats (CRISPR)-associated protein 9 (Cas9), which relies on its ability to cleave genomic DNA with target sequences, is the most widely used genome editing tool (Jinek et al., 2012). The double-strand breaks at target sites induced by CRISPR-Cas9 mainly evoke two DNA repair pathways. One is non-homologous end-joining (NHEJ), which joins two broken ends of DNA with diverse insertions and deletions (indels) at the joined sites. The other is homology-directed repair (HDR), which repairs broken DNA by recombination between genomic DNA and template DNA with sequence homology (Gaj et al., 2016). Therefore, we can achieve precise genetic manipulation via HDR by providing cells with donor DNA with the intended sequences. In general, NHEJ is much more frequent than HDR, and both HDR and NHEJ can be concurrently induced in the same cell. As a result, even when we attempt to induce specific genetic manipulations via HDR in iPS cells, the resulting clones have diverse genotypes. Therefore, it is important to understand the diverse allelic combinations and frequencies of genome editing in iPSCs.

Despite the importance of monitoring genome editing outcomes in individual cells, most assays analyze genome editing results in cell populations, but not in single cells, for example, the T7E1 assay and pooled amplicon sequencing (Germini et al., 2018). As a new approach, we previously isolated more than 2,600 clones of genome-edited human cultured cells (HEK293T, HeLa, and PC9 cells) using an automated single-cell dispensing system (Takahashi and Miyaoka, 2023). By analyzing the genotypes of these isolated clones, we found a strong binary tendency of genome editing induced by CRISPR-Cas9; that is, individual cells are often either not edited at all or all target alleles are fully edited (Takahashi et al., 2022).

However, owing to the high mortality, we could not apply the same single-cell dispensing system to human iPS cells as human cultured cell lines. Efficient methods to isolate human iPS cell clones are in high demand, not only for the analysis of genome editing results, but also for the isolation of iPS cell lines with desired genetic manipulations. Recent studies have reported promising additives to avoid cell death in iPS cells, but they are not yet conclusive (Chen et al., 2021). Therefore, in this study, we developed a method that utilizes a cell-handling robot to efficiently isolate a large number of clones from genome-edited iPS cell pools. Using this method, we obtained more than 1,000 genome-edited iPS cell clones and analyzed their genotypes. We found that the same genetic manipulations (HDR and various indels) were homozygously induced in human iPS cells. Our new approach to efficiently isolate human iPS cell clones and profiles of genome editing outcomes in human iPS cells will greatly contribute to regenerative medicine.

## Results

### Development of an efficient robotic isolation method for genome-edited iPS cell clones grown in Matrigel domes

First, we developed a method to isolate clones from genome-edited iPS cells using a cell-handling robot, CELL HANDLER (Yamaha Motor) (Figure 1A). In this method, iPS cells were transfected with a plasmid (px459-HypaCas9) to express HypaCas9 and single-stranded donor DNA to induce a pathogenic point mutation (ATP7B R778L, GRN R493X, or RBM20 R636S) (Kato-Inui et al., 2018). The puromycin-resistant gene was co-expressed with HypaCas9 via the T2A peptide so that the transfected cells were selected using puromycin and then dispersed into single cells. We formed domes composed of Matrigel and allowed these single iPS cells to grow to maximize the number of single cells growing in a well of a 6-well plate (Figure 1B and 1C). Placing cells in a 3-dimensional structure also made the robotic cell-picking process less damaging and more efficient than with 2-dimensional cultures. Culturing these single cells for approximately 1 week resulted in the formation of cell clumps with a diameter of 100–150 μm, which were suitable for being picked by CELL HANDLER (Figure 1D). CELL HANDLER captured multiple images with an automatic focus to scan for cell clumps throughout the Matrigel domes. Specialized software processed these images to recognize and distinguish cell clumps by major diameter, circularity, and neighbor distance (Table S1). We measured these criteria for typical cell clumps derived from single cells (Figure 1D). Cell clumps that met these criteria were selected and transferred into a new 96-well plate for expansion in canonical two-dimensional culture. After 2-4 weeks, we isolated genome-edited iPS cell clones that maintained the expression of pluripotency marker genes (Figure S1A). The overall cloning efficiency was approximately 60.1% (1372 clones out of 2255 wells in 36 96-well plates) using CELL HANDLER (Table 1 and Figure S1B).

**Figure 1.**
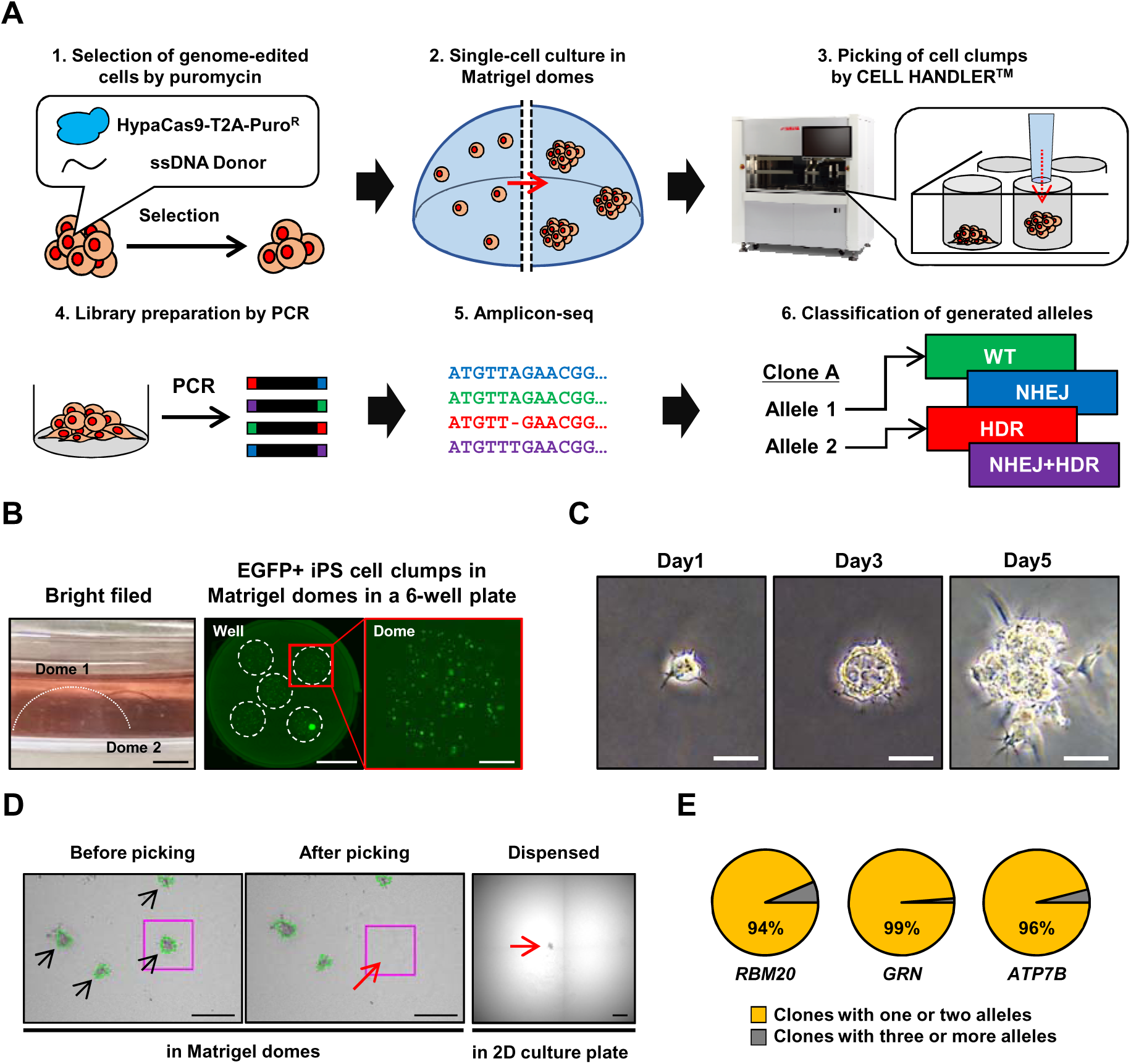
Robotic isolation of genome-edited human iPS cell clones. **(A)** Schematic representation of the robotic isolation of genome-edited human iPS cell clones. After transfection of HypaCas9 and donor DNA, and selection with puromycin, genome-edited single iPS cells were cultured in Matrigel domes to form cell clumps (top left and top center). Cell clumps were isolated by robotic picking by CELL HANDLER (top right). Clones were genotyped by amplicon sequencing (bottom left, center, and right). **(B)** Bright-field and fluorescence images of Matrigel domes in wells and EGFP-positive iPS cells growing in Matrigel domes. Five to six Matrigel domes were created in each well of a 6-well plate. Scale bar: left and right panels = 2 mm, center panel = 1 cm). **(C)** Tracking images of a single iPS cell forming a cell clump in a Matrigel dome from day 1 to 5. Scale bar: 10 μm. **(D)** Picking and seeding of cell clumps by CELL HANDLER. CELL HANDLER captured images before and after picking. In the “Before picking” image, the black arrows indicate target clumps recognized by CELL HANDLER (outlined with green lines). In the “After picking” image, the magenta square indicates the position of a cell clump picked by CELL HANDLER. The “Dispensed” image shows the cell clump immediately after seeding into the 96-well plate (red arrow). Scale bar: 300 μm. **(E)** Proportion of the isolated iPS cell clones with different allelic numbers. We analyzed the number of allelic types in isolated clones. Clones with ≥ 3 allelic types were removed from further analysis, as those clones may not be derived from single genome-edited iPS cells.

**Table 1.**
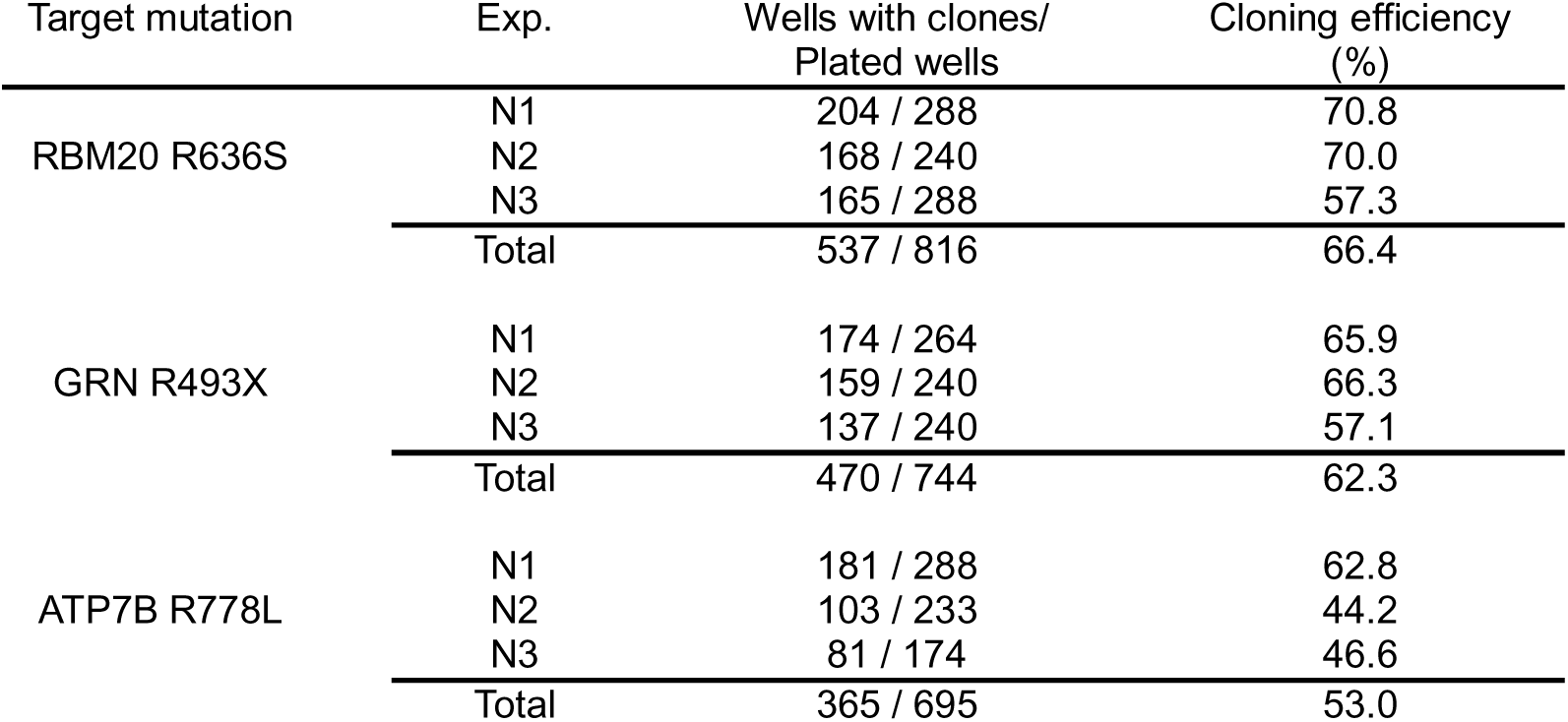
iPS cell cloning efficiencies in 96-well plates by robotic picking.

### Assessment of clonality of iPS cells isolated by robotic picking

To assess the clonality of the isolated iPS cells by robotic picking, we established iPS cell lines expressing EGFP, mCherry, or EBFP. We then mixed these three cell lines and performed robotic picking (Figure S1C). We repeated this cloning process twice and isolated 28 and 43 clones. We confirmed that none of the clones contained 2 or more colors (Table S2). We then analyzed the number of alleles in isolated iPS cells after editing RBM20, GRN, and ATP7B. We investigated the genome editing outcomes by amplicon sequencing as previously reported (Takahashi et al., 2022), and classified the resulting alleles into 4 types: wild-type (WT), NHEJ, HDR, and NHEJ+HDR, using CRISPResso2 (Clement et al., 2019). This allelic classification revealed that 96.3% of the isolated clones had 1 or 2 allelic types on average at the 3 target sites (Figure 1E). These results suggest that the majority of the isolated clones were derived from single cells. In contrast, 3.7% of clones harbored three or more allelic types, and these clones were excluded from subsequent analyses.

### Homozygous NHEJ is the most common genotype in genome-edited iPS cells

Because iPS cells are diploid, there are 10 different genotypes: WT/WT, WT/NHEJ, WT/HDR, WT/NHEJ+HDR, NHEJ/NHEJ, NHEJ/HDR, NHEJ/NHEJ+HDR, HDR/HDR, HDR/NHEJ+HDR, and NHEJ+HDR/NHEJ+HDR. We classified the genotypes of isolated iPS clones and found that NHEJ/NHEJ, WT/WT, and WT/NHEJ were the most common, second, and third most common genotypes, respectively, in all 3 target genes (Figure 2A).

**Figure 2.**
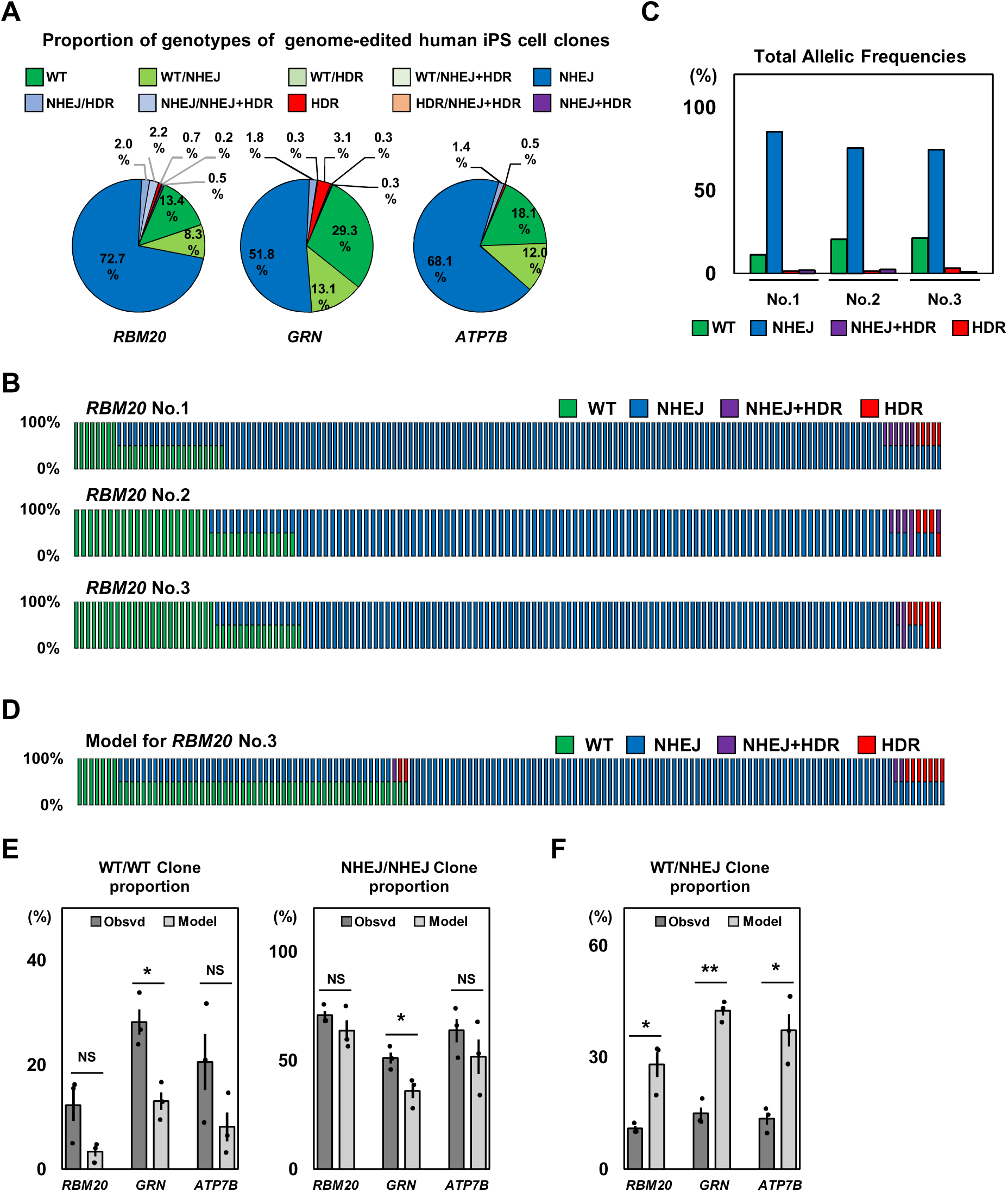
Genotypes of genome-edited iPS cell clones. **(A)** Proportion of genotypes of genome-edited human iPS cell clones. Almost no WT/HDR and WT/NHEJ+HDR clones were isolated. **(B)** Genome editing outcomes in isolated clones derived from single human iPS cells edited by Cas9. RBM20 editing outcomes are shown (No.1 to No.3). Each bar represents 1 clone, and the genotypes of WT (green), NHEJ (blue), HDR (red), and HDR + NHEJ (purple) in 1 clone are also shown in each bar. **(C)** Total allelic frequencies of WT (green), NHEJ (blue), HDR (red), and HDR + NHEJ (purple) in genome-edited iPS cells in the 3 experiments shown in (B). **(D)** Model of the distributions of clones with different genotypes assuming different alleles are randomly distributed at the observed frequencies. The model for RBM20 No.3 is shown. **(E)** Comparison of the proportions of WT/WT and NHEJ/NHEJ clones between the mathematical models and the observed cells in RBM20, GRN, and ATP7B. Student’s t-test was used to evaluate differences. **p <0.01, *p <0.05 and NS: not significantly different (p >0.2). **(F)** Comparison of the proportions of WT/NHEJ clones between the mathematical models and the observed cells in RBM20, GRN, and ATP7B. Student’s t-test was used to evaluate differences. **p <0.01 and *p <0.05

We previously reported that genome editing in cultured cells, such as HEK293T, occurred in a binary manner; that is, all targeted sequences were either edited or not edited (Takahashi et al., 2022). Therefore, we examined whether the genome editing of iPS cells also occurred in a binary manner. For this purpose, we investigated the proportion of WT/WT, WT/NHEJ, and NHEJ/NHEJ clones. We created diagrams to visualize the genotypes of the isolated clones (Figure 2B, S2A, and S2B). We also generated model diagrams to represent genotypes of the same number of clones if all allele types were randomly distributed, as described previously (Takahashi et al., 2022). We obtained the overall frequencies of the WT, NHEJ, HDR, and NHEJ+HDR alleles from the genotypes of the isolated clones (Figure 2C, S2C, and S2D). These overall allelic frequencies were nearly identical to those of the pooled populations of genome-edited iPS cells before clone isolation, suggesting that our robotic clone-picking method does not cause any bias in the genotypes of the isolated cells (Figure 2A and S3). We uniformly and randomly redistributed these alleles in the same number of clones as the isolated ones, which served as models for comparison (Figure 2D, S4A, S4B, and S4C). Relative to these models, we generally observed more isolated WT/WT and NHEJ/NHEJ clones, although the differences were not statistically significant for RBM20 and ATP7B (Figure 2E). The proportion of WT/NHEJ clones was significantly lower in the isolated clones than in the model for all three edited genes, indicating that genome editing in human iPS cells was also binary (Figure 2F).

### Frequent homozygous induction of identical indels by NHEJ in iPS cells

So far in our study, we have classified all different indels into a single category: “NHEJ”. Although NHEJ induced diverse indels, there were several specific indels with notable frequencies. Therefore, we re-genotyped the isolated genome-edited iPS clones by distinguishing the frequent indels in each gene.

We re-analyzed the amplicon sequencing data, identified the top eight most frequent indel alleles, and classified 12 alleles in the isolated iPS clones genome-edited in RBM20, GRN, and ATP7B, which represented approximately 70% of the overall allelic frequency (Figure 3A, 3B, S5A-B, S6B, and S6D). We found that many iPS clones were homozygous for identical indels. In particular, several clones were homozygous for an indel at a low frequency (Figure S6A, S6C, and S6E). For example, we isolated one clone with a homozygous 1-bp deletion in RBM20 No.3 (overall frequency was only 1.3%) and two clones with a homozygous 11-bp deletion from RBM20 No.1 and 3 (1.4%) (Figures 3A and 3B). Therefore, we compared the proportion of clones with homozygous alleles for the 3 genes. We examined the frequencies of iPS cell clones homozygous for the top three frequent indel alleles in all three target genes and found that they were all higher than those of the models, although some of them were not statistically significant (Figures 3C, 3D, S6A, S6C, and S6E). Overall, genome editing in iPS cells tends to result in identical sequence manipulations in both copies of the target sequences in single cells.

**Figure 3.**
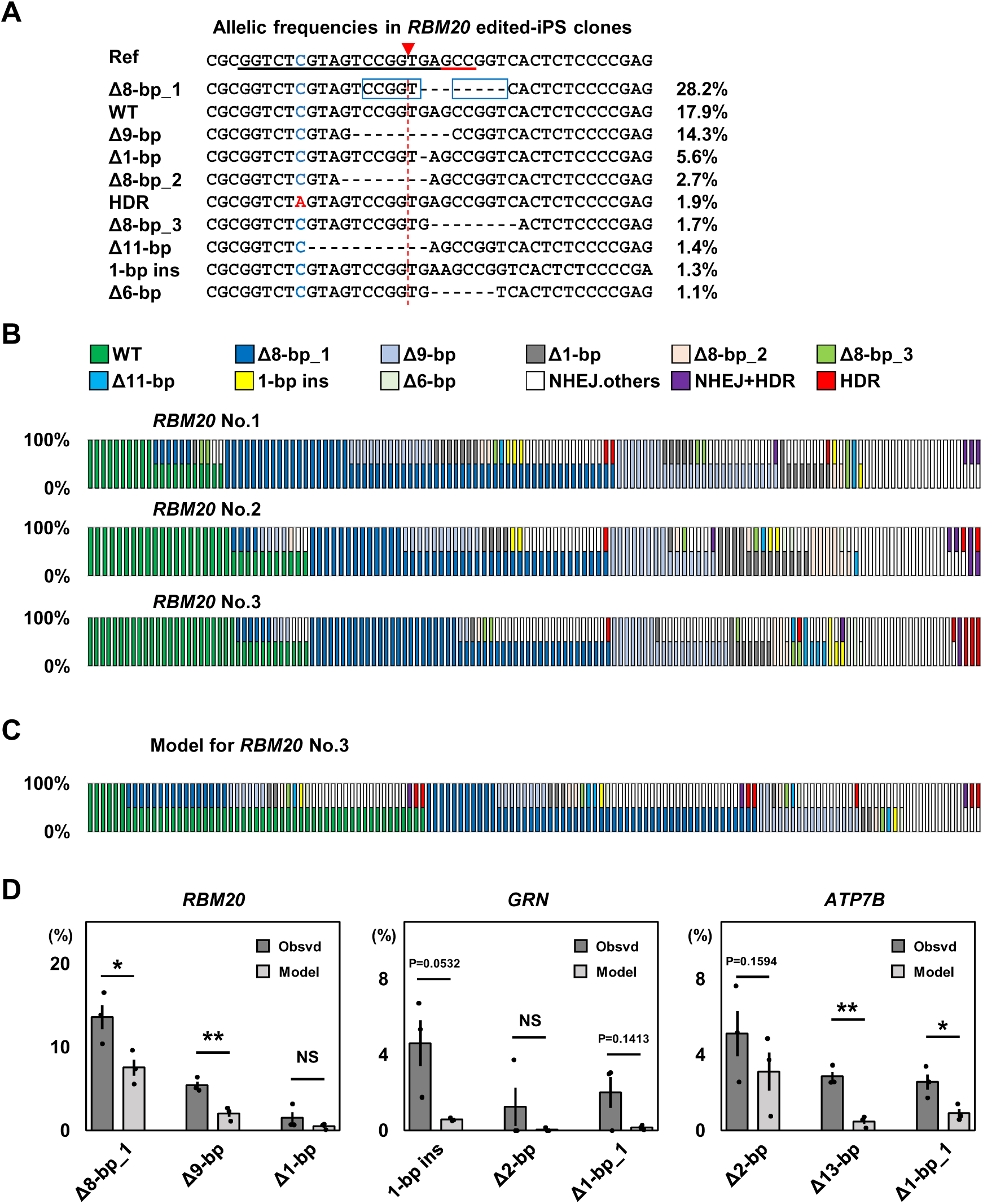
Profiles of various indels induced by NHEJ. **(A)** Sequences and frequencies of different alleles generated by RBM20 R636S editing. The 10 most frequently observed alleles after RBM20 editing are shown. Black and red underline indicates guide RNA and PAM sequences, respectively. Red triangles and red dotted lines indicate the cleavage site by Cas9. Blue and red characters indicate unedited and substituted nucleotides, respectively. Blue boxes highlight the microhomology sequence (CCGGT). **(B)** Genome editing outcomes with distinguished NHEJ sequences in isolated clones derived from single human iPS cells edited by Cas9. RBM20 editing outcomes are shown (No.1 to No.3) **(C)** Model of the distributions of clones with different genotypes assuming different alleles are randomly distributed at the observed frequencies. The model for RBM20 No. 3 is shown. **(D)** Comparison of the proportions of homozygous NHEJ clones with the top 3 most frequent indels between the mathematical models and the actually observed cells. Values ±S.E. are shown (n = 3). Student’s t-test was used to evaluate differences. **p <0.01, *p <0.05 and NS: not significantly different (p >0.2).

### Robotic isolation of genome-edited iPS cell lines with rare genotypes

Because our new method based on robotic handling of cells allowed high-throughput isolation of genome-edited iPS cell clones, we tested whether this method allowed us to establish iPS clones with rare genotypes. As shown in Figure 3B, genome editing in iPS cells produced mainly non-edited cells or cells homozygous for NHEJ, and cells with heterozygous genotypes were relatively rare. Therefore, we characterized 2 clones with heterozygous NHEJ in GRN, whose mutations are associated with frontotemporal lobar degeneration (Chen-Plotkin et al., 2011). Based on our amplicon sequencing analysis, 1 clone (GRN-Het 1-bp ins) had a heterozygous 1-bp insertion, and the other clone (GRN-Het Δ19-bp) had a heterozygous 19-bp deletion (Figure 4A). Both mutations were expected to cause nonsense-mediated decay owing to frameshift mutations (Figure 4A). Because we extracted genomic DNA from isolated clones while freezing a portion of these clones, we thawed and recovered these two clones. Sanger sequencing and digital PCR confirmed that the recovered clones maintained the identified heterozygous mutations (Figure 4B, 4C, and S7). In addition, these clones maintained the expression of the pluripotency markers SOX2 and OCT4 (Figure 4D). These results indicate that our iPS cell cloning method allows the establishment of clones with rare genotypes, which is highly useful for genome editing in iPS cells to study and cure diseases.

**Figure 4.**
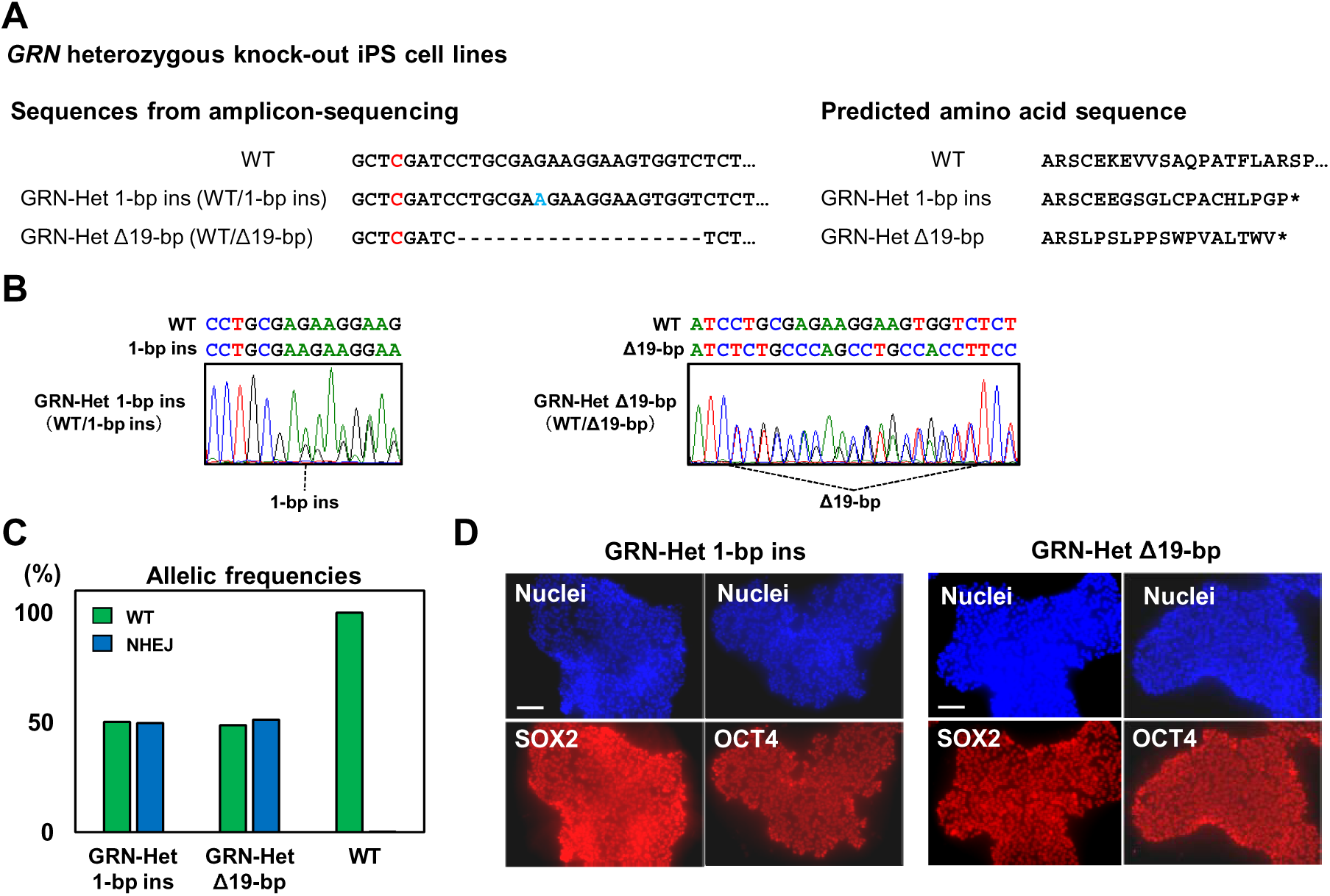
Maintenance of pluripotency and genotypes after robotic isolation of iPS clones. **(A)** Two iPS cell lines with heterozygous NHEJ in GRN identified by amplicon sequencing. DNA sequences and predicted amino acid sequences are shown. These 2 indels were expected to be nonsense mutations. **(B)** Sanger sequencing of the 2 heterozygous NHEJ iPS cell lines. **(C)** Digital PCR analysis of the NHEJ frequency in the 2 heterozygous NHEJ iPS cell lines. The 2 lines had approximately 50% wild-type and 50% NHEJ alleles. WT genomic DNA is the control for digital PCR. **(D)** Immunocytochemistry of SOX2 and OCT4 in the 2 GRN heterozygous KO iPS cell lines. Scale bar: 100 μm.

## Discussion

Since complete control of genome editing has not yet been achieved, we must rely on single-cell cloning to establish genome-edited iPS cell lines with desired genotypes. In this study, we achieved high cloning efficiency using iPS cell culture in a 3-dimensional Matrigel dome and robotic isolation of cell clumps using CELL HANDLER. Small molecule additives, such as the CEPT cocktail, have been developed to avoid cell death induced by the singularization of human iPS cells to enhance single-cell cloning (Tristan et al., 2023). Therefore, a combination of our robotic handling method and these small-molecule additives may further enhance the single-cell cloning of iPS cells.

We previously reported that genome editing in cultured human cells is induced in a binary manner, where all target alleles are either completely edited or not edited at all. We found this dichotomous effect in human iPS cells as well, as we observed fewer WT/NHEJ heterozygously edited clones relative to the mathematical expectation (Figure 2F). Patients with some genetic disorders are heterozygous for mutations. Our findings indicate that establishing human iPS cells with these heterozygous mutations using CRISPR-Cas9 is challenging. Recently, Kawamata et. al. reported the same binary trend in genome editing and demonstrated that decreasing cleavage activity allows for heterozygous editing (Kawamata et al., 2023). Therefore, to achieve heterozygous editing of human iPS cells, it may be necessary to lower CRISPR-Cas9 activity.

In this study, we investigated and distinguished indels with different sequences. Deletions were the predominant genome editing outcome for all target genes (Figures 3A, S5A, and S5B). We also observed 1-bp insertions in the cleavage regions of the three genes. As reported in previous studies, these 1-bp insertions presumably occur because Cas9 occasionally cleaves DNA sequences with single-base protrusions (Chakrabarti et al., 2019; Lemos et al., 2018; Shen et al., 2018). Moreover, many of the observed deletions were derived from microhomology-mediated end-joining (MMEJ), as previously reported by other groups (Chen et al., 2019; Kim et al., 2018; van Overbeek et al., 2016; Shen et al., 2018). The most frequent 8-bp deletion in RBM20 was a typical example of MMEJ, in which a 5-bp microhomology (CCGGT) mediated the deletion (Figure 3A).

Interestingly, these NHEJ (including MMEJ) alleles were found to be homozygously induced in many clones (Figure 3D). The DNA repair mechanism is cell cycle-dependent and NHEJ is typically active throughout the cell cycle, whereas HDR is only active in the S/G2 phase (Danner et al., 2017). One possibility is that 1 of the 2 alleles was first edited by NHEJ and the other allele was edited by HDR using the NHEJ allele as a homologous template, resulting in 2 identical NHEJ alleles. Further studies are needed to confirm this hypothesis. In summary, our study provides new insights into genome editing in human iPS using CRISPR-Cas9. Single-cell cloning remains the most straightforward strategy for isolating genome-edited iPS cell clones. However, the manual selection of cell colonies requires considerable work and time. Moreover, numerous culture plates are required to grow iPS cell colonies for picking, while keeping these colonies separated from each other in 2-dimensional culture. In our method, CELL HANDLER takes the burden of colony picking, and the 3-dimensional culture allows a large number of clones to grow as cell clumps. These advantages enable the large-scale establishment of genome-edited iPS clones, which will be highly valuable for studying genetic disorders and developing gene therapies using iPS cells.

## Supporting information

Supplemental Infomation

## Experimental Procedures

### Resource availability

#### Corresponding author

Further information and requests for resources and reagents should be directed to and will be fulfilled by the corresponding author.

#### Materials availability

All reagents and materials used in this manuscript are available upon request or prepared for availability from commercial sources. GRN heterozygous knockout iPS cell lines generated in this study will be made available on request, but we may require a payment and/or a completed material transfer agreement if there is potential for commercial application.

#### Data and code availability

The amplicon sequencing data analyzed in this study will be shared upon request. Raw amplicon sequencing data are available at the DDBJ Sequence Read Archive.

### Plasmids and single-stranded DNA (ssDNA) donor

The px459-HypaCas9 plasmid used in this study to express HypaCas9 and the puromycin-resistant gene has been described previously (Kato-Inui et al., 2018) (Addgene Plasmid #108294). The single-stranded DNA donors and gRNAs used in this study have also been reported previously (Takahashi et al., 2022). The sequences are listed in Tables S3.

### Maintenance of iPS cells

WTC11 iPS cells (Coriell Institute for Medical Research, GM25256) were used for all experiments. iPS cells were maintained on thin-coated GFR Matrigel Matrix (Corning, 356231) in mTeSR Plus (STEMCELL Technologies, ST-100-0276) medium supplemented with 1% penicillin-streptomycin (P/S) (Nacalai tesque, 26253-84). Cells were dissociated using the Accutase (Nacalai tesque, 12679-54), to passaged wells, we added Y-27632 (final concentration 10 uM), a Rho-associated kinase inhibitor (Ri) (FUJIFILM Wako Pure Chemical, 034-24024), to promote cell survival.

### Transfection

WTC11 iPS cells were seeded at 4×10^4 cells/well in a Matrigel-coated 24-well plate. The next day, at least one hour before transfection, the medium was replaced with 500 μL of fresh mTeSR Plus with P/S and Ri. Lipofectamine Stem (Thermo Fisher, STEM00003) was used to transfect 400 ng/well of px459-HypaCas9 and 100 ng/well of ssDNA according to the manufacturer’s instructions. After 1 and 2 days, the medium was replaced with 500 μL of fresh mTeSR Plus with P/S. At 4-6 days after transfection, when the cells reached confluence, we dissociated the iPS cells into single cells and embedded them into Matrigel domes.

### Formation of Matrigel domes containing single iPS cells

GFR Matrigel (Corning, 356231) was kept on ice until use. Confluent iPS cells on a 24-well plate were detached from the plate using 100 μL/well of Accutase. Then, 400 μL of PBS was added to suspend the cells, which were centrifuged at 300 × *g* for 3 min. The supernatant was removed and 1 mL of mTeSR Plus with P/S and Ri was added to gently suspend the cells. After counting the number of cells using Countess II (Thermo Fisher Scientific), the cell suspension was diluted to 10-30 cells/μL. Matrigel (400 μL) was dispensed into a new 1.5 mL tube on ice, to which 100 μL of diluted cell suspension was added and mixed gently. Then, 50 μL of the cell and Matrigel mixture was aliquoted into each well of a 6-well plate (5-6 domes/well) (Figure 1B). Immediately after placing aliquots, the bottom of the plate was warmed by hand for 2-3 min to allow the domes to start to gel. The plate was then inverted and kept at 37°C for 2 h to solidify the Matrigel domes. After solidification, 3 mL of mTeSR Plus with P/S and Ri were added and kept in a CO_2_ incubator.

### Recognition, picking, and seeding of iPS cell clumps by CELL HANDLER

Recognition, picking, and seeding of iPS cell clumps by CELL HANDLER (Yamaha Motor) when the cell clumps reached a size of 100-200 μm diameters. CELL HANDLER acquired a total of 28 focus images in the Z-axis for a Matrigel dome to select cell clumps to pick. If cell clumps matched the parameters for picking, CELL HANDLER carried out the picking and seeding (Figure 1D and Table S1). The picked cell clumps were automatically transferred to the wells of a Matrigel-coated 96-well plate containing 100 μL of preheated mTeSR Plus with P/S and Ri medium. CELL HANDLER captured images of the cell clumps before and after picking to verify the successful acquisition of cell clumps from the Matrigel domes. Supplemental movies (Video S1) show picking processes using CELL HANDLER.

### Expansion and freezing of isolated iPS cell clones

iPS clones were cultured for 2-3 weeks after robotic picking to ensure sufficient clone expansion. The medium was removed, and 30 μL/well of Accutase was added to detach and resuspend the cells. Half of the cell suspension was transferred to a new 96-well plate and used for genome extraction as described previously (Miyaoka et al., 2014). The other half of the cell suspension in the original 96-well plate was mixed with 75 μL of 10% dimethyl sulfoxide (DMSO) and 90% fetal bovine serum. The mixtures were layered with 75 μL of mineral oil, and the plate was sealed with Parafilm for cryopreservation at -80°C. The two clone lines in Fig. 4A were selected by amplicon sequencing analysis as described below, in which clones with heterozygous knockout of the GRN gene were selected. To confirm that these clones were pure, genotypes were identified by quantifying allele frequencies using digital PCR, as described previously (Miyaoka et al., 2014).

### Preparation of multiplexed amplicon sequencing libraries

Multiplexed amplicon sequencing libraries were prepared using 2 rounds of PCR, as described previously (Takahashi et al., 2022). After the second PCR, the amplified DNA fragments from the pooled 48 samples were purified by gel extraction using the NucleoSpin Gel and PCR Clean-up Midi kit (TaKaRa, 740986.20). We repeated this DNA purification process every 48 samples. The purified DNAs was mixed in equal molar ratios, and the library was prepared according to Illumina’s instructions. DNA concentrations of the mixed libraries were quantified using the GenNext NGS Library Quantification Kit (Toyobo, NLQ-101). After quantification, PhiX Control v3 (Illumina, FC-110-3001) was added at a final concentration of 20% for amplicon sequencing. Sequencing was performed with MiSeq (Illumina) using the MiSeq v2 reagent kit (Illumina, MS-102-2003) or MiSeq Reagent kit v2 Nano (Illumina, MS-103-1001) according to the manufacturer’s instructions.

### Amplicon sequencing data analysis and allele classification

Fastq files generated by MiSeq were imported into CLC Genomics Workbench (QIAGEN). Adapter sequences were removed and demultiplexed using the DNA Index. The data were then analyzed using CRISPResso2 (https://github.com/pinellolab/CRISPResso2) in CRISPResso Batch mode (Clement et al., 2019). CRISPResso2 was installed as recommended using a Docker containerization system. In this study, all reads identified as ambiguous by a CRISPResso2 analysis were classified as NHEJ.

Mathematical models for the distribution of clone genotypes, assuming that the WT, NHEJ, HDR, and HDR+NHEJ alleles were randomly induced in all target alleles, were built by distributing these edited alleles to the isolated iPS cell clones at their observed overall frequencies, as described previously (Takahashi et al., 2022). We also distinguished the major NHEJ alleles based on amplicon sequencing data. These 8 NHEJ alleles, together with WT, HDR, and HDR+NHEJ alleles, accounted for approximately 60-70% of the total alleles. Other NHEJ alleles were grouped and labeled “NHEJ.other”.

### Immunocytochemistry

iPS cells were gently washed in PBS and fixed in 4% PFA (EMS; 50-980-487) for 15 min. Samples were washed 3 times in PBS and incubated with blocking buffer (3% goat normal serum and 0.2% Triton-X in PBS) for 1 h at room temperature. Samples were then incubated overnight with the primary antibody (Table S4) at the appropriate concentration in primary blocking buffer at 4°C overnight, washed 3 times in DPBS, and incubated with secondary antibodies (Table S4) diluted in blocking buffer at room temperature for 1 h. The samples were incubated with Hoechst 33342 (Nacalai tesque, 04929-82) for 10 min at room temperature and washed twice with PBS. Fluorescent images were captured using a Keyence BZ-X800 all-in-one microscope (Keyence) and analyzed using BZ-X Analyzer software (Keyence).

### Statistics

Transfection was performed in triplicate (three biological replicates). Values are displayed as the mean ± standard error (S.E.). Statistical significance between the 2 groups was assessed by a non-paired 2-tailed Student’s t-test and is displayed in the figures with asterisks as follows: *p < 0.05; **p < 0.01. NS: not significantly different (p > 0.2).

## Supplemental Information

Supplemental information can be found online.

Figures S1-S7

Tables S1-S4

Videos S1

## Author Contributions

G.T. and Y.M. conceived of the study and designed the experiments. G.T. and M.M. transfected iPS cells and encapsulated a single iPS cell in the Matrigel dome. G.H. performed pilot experiments to set up the parameters for CELL HANDLER. S.I. performed cell clamp picking using CELL HANDLER presented in this study. G.T., M.M., and K.S. classified the genotypes obtained using amplicon sequencing. G.T. and K.S. created the NHEJ model. Y.M. supervised the project. G.T. and Y.M. wrote the manuscript with the help of all the authors.

## Acknowledgments

This work was supported by the Japan Society for the Promotion of Science (JSPS) Grant-in-Aid for Challenging Research (Pioneering) 20K21409, Grant-in-Aid for Scientific Research (B) 20H03442 and 24K02028, Interstellar Initiative Beyond from Japan Agency for Medical Research and Development (AMED) (23jm0610092h0001), Takeda Science Foundation Medical Research Grant, Sumitomo Foundation Grant for Basic Science Research Projects, Ichiro Kanehara Foundation Grant to Y.M., and JSPS Grant-in-Aid for Early-Career Scientists (18K15054 and 22K15386). We thank Y. Utsumi and R. Abe (Yamaha Motor Co., Ltd.) and all lab members for their helpful discussions.

## References

Chakrabarti, A.M., Henser-Brownhill, T., Monserrat, J., Poetsch, A.R., Luscombe, N.M., and Scaffidi, P. (2019). Target-Specific Precision of CRISPR-Mediated Genome Editing. Mol. Cell 73, 699–713.e6.

Chen, W., McKenna, A., Schreiber, J., Haeussler, M., Yin, Y., Agarwal, V., Noble, W.S., and Shendure, J. (2019). Massively parallel profiling and predictive modeling of the outcomes of CRISPR/Cas9-mediated double-strand break repair. Nucleic Acids Res. 47, 7989–8003.

Chen, Y., Tristan, C.A., Chen, L., Jovanovic, V.M., Malley, C., Chu, P.-H., Ryu, S., Deng, T., Ormanoglu, P., Tao, D., et al. (2021). A versatile polypharmacology platform promotes cytoprotection and viability of human pluripotent and differentiated cells. Nat. Methods 18, 528–541.

Clement, K., Rees, H., Canver, M.C., Gehrke, J.M., Farouni, R., Hsu, J.Y., Cole, M.A., Liu, D.R., Joung, J.K., Bauer, D.E., et al. (2019). CRISPResso2 provides accurate and rapid genome editing sequence analysis. Nat. Biotechnol. 37, 224–226.

Danner, E., Bashir, S., Yumlu, S., Wurst, W., Wefers, B., and Kühn, R. (2017). Control of gene editing by manipulation of DNA repair mechanisms. Mamm. Genome 28, 262–274.

Gaj, T., Gersbach, C.A., and Barbas, C.F. (2013). ZFN, TALEN, and CRISPR/Cas-based methods for genome engineering. Trends Biotechnol. 31, 397–405.

Gaj, T., Sirk, S.J., Shui, S.-L., and Liu, J. (2016). Genome-Editing Technologies: Principles and Applications. Cold Spring Harb. Perspect. Biol. 8.

Germini, D., Tsfasman, T., Zakharova, V.V., Sjakste, N., Lipinski, M., and Vassetzky, Y. (2018). A comparison of techniques to evaluate the effectiveness of genome editing. Trends Biotechnol. 36, 147–159.

Hockemeyer, D., and Jaenisch, R. (2016). Induced pluripotent stem cells meet genome editing. Cell Stem Cell 18, 573–586.

Jinek, M., Chylinski, K., Fonfara, I., Hauer, M., Doudna, J.A., and Charpentier, E. (2012). A programmable dual-RNA-guided DNA endonuclease in adaptive bacterial immunity. Science 337, 816–821.

Kato-Inui, T., Takahashi, G., Hsu, S., and Miyaoka, Y. (2018). Clustered regularly interspaced short palindromic repeats (CRISPR)/CRISPR-associated protein 9 with improved proof-reading enhances homology-directed repair. Nucleic Acids Res. 46, 4677–4688.

Kawamata, M., Suzuki, H.I., Kimura, R., and Suzuki, A. (2023). Optimization of Cas9 activity through the addition of cytosine extensions to single-guide RNAs. Nat. Biomed. Eng. 7, 672–691.

Kim, S.-I., Matsumoto, T., Kagawa, H., Nakamura, M., Hirohata, R., Ueno, A., Ohishi, M., Sakuma, T., Soga, T., Yamamoto, T., et al. (2018). Microhomology-assisted scarless genome editing in human iPSCs. Nat. Commun. 9, 939.

Lemos, B.R., Kaplan, A.C., Bae, J.E., Ferrazzoli, A.E., Kuo, J., Anand, R.P., Waterman, D.P., and Haber, J.E. (2018). CRISPR/Cas9 cleavages in budding yeast reveal templated insertions and strand-specific insertion/deletion profiles. Proc. Natl. Acad. Sci. USA 115, E2040–E2047.

Miyaoka, Y., Chan, A.H., Judge, L.M., Yoo, J., Huang, M., Nguyen, T.D., Lizarraga, P.P., So, P.-L., and Conklin, B.R. (2014). Isolation of single-base genome-edited human iPS cells without antibiotic selection. Nat. Methods 11, 291–293.

van Overbeek, M., Capurso, D., Carter, M.M., Thompson, M.S., Frias, E., Russ, C., Reece-Hoyes, J.S., Nye, C., Gradia, S., Vidal, B., et al. (2016). DNA Repair Profiling Reveals Nonrandom Outcomes at Cas9-Mediated Breaks. Mol. Cell 63, 633–646.

Shen, M.W., Arbab, M., Hsu, J.Y., Worstell, D., Culbertson, S.J., Krabbe, O., Cassa, C.A., Liu, D.R., Gifford, D.K., and Sherwood, R.I. (2018). Predictable and precise template-free CRISPR editing of pathogenic variants. Nature 563, 646–651.

Takahashi, G., and Miyaoka, Y. (2023). Large-scale single-cell cloning of genome-edited cultured human cells by On-chip SPiS. STAR Protocols 4, 102364.

Takahashi, G., Kondo, D., Maeda, M., Morishita, Y., and Miyaoka, Y. (2022). Genome editing is induced in a binary manner in single human cells. IScience 25, 105619.

Tristan, C.A., Hong, H., Jethmalani, Y., Chen, Y., Weber, C., Chu, P.-H., Ryu, S., Jovanovic, V.M., Hur, I., Voss, T.C., et al. (2023). Efficient and safe single-cell cloning of human pluripotent stem cells using the CEPT cocktail. Nat. Protoc. 18, 58–80.

