## Supplemental Figure for "High-throughput robotic isolation of human iPS cell clones reveals frequent homozygous induction of identical genetic manipulations by CRISPR-Cas9"

### Slide 1
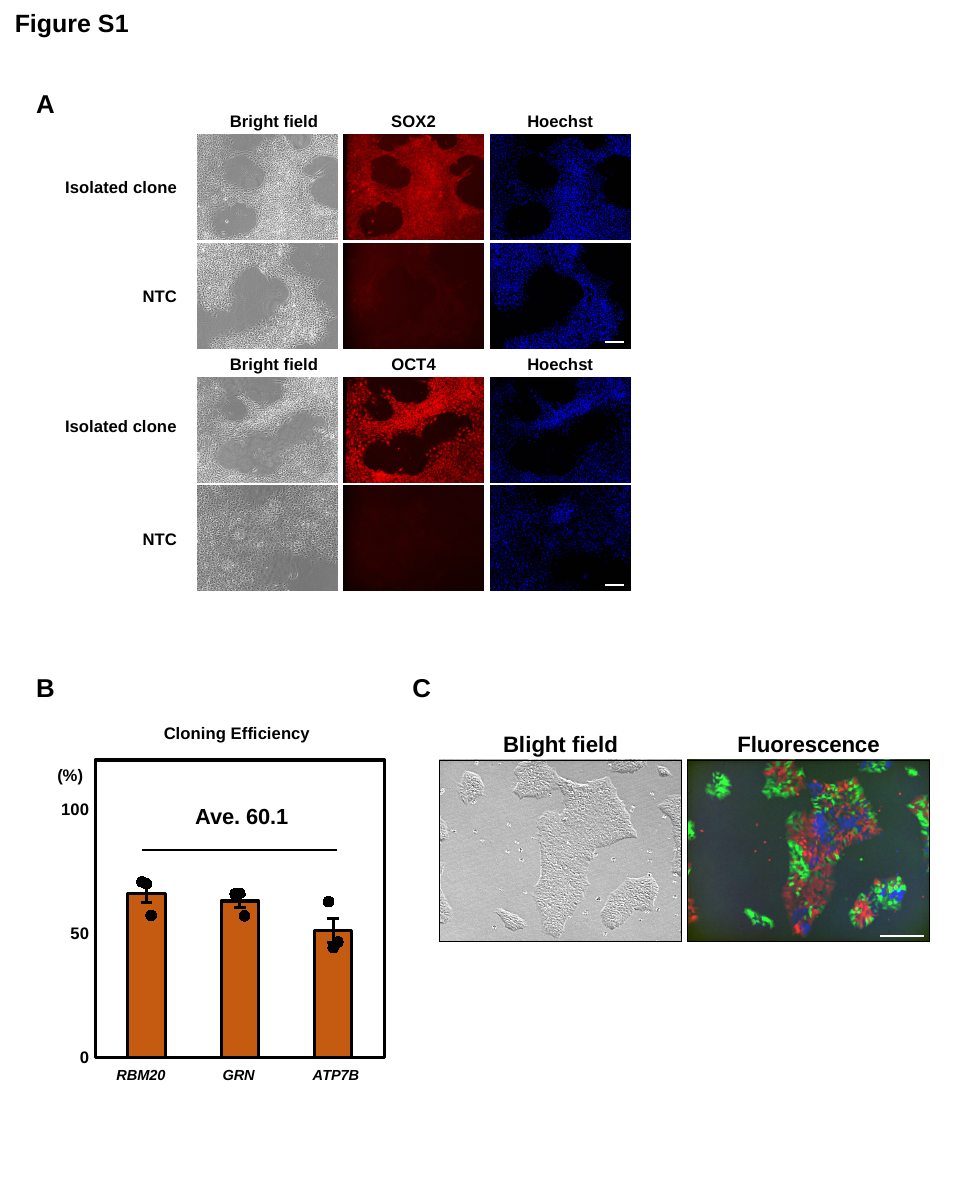

Figure S1
A
Bright field
SOX2
Hoechst
Isolated clone
NTC
Bright field
OCT4
Hoechst
Isolated clone
NTC
B
C
#### Chart
| Category | Single cell cloning 比較 | 系列1 | 系列2 | 系列3 |
|---|---|---|---|---|RBM20
GRN
ATP7B
Cloning Efficiency
(%)
Ave. 60.1
Blight field
Fluorescence

### Slide 2
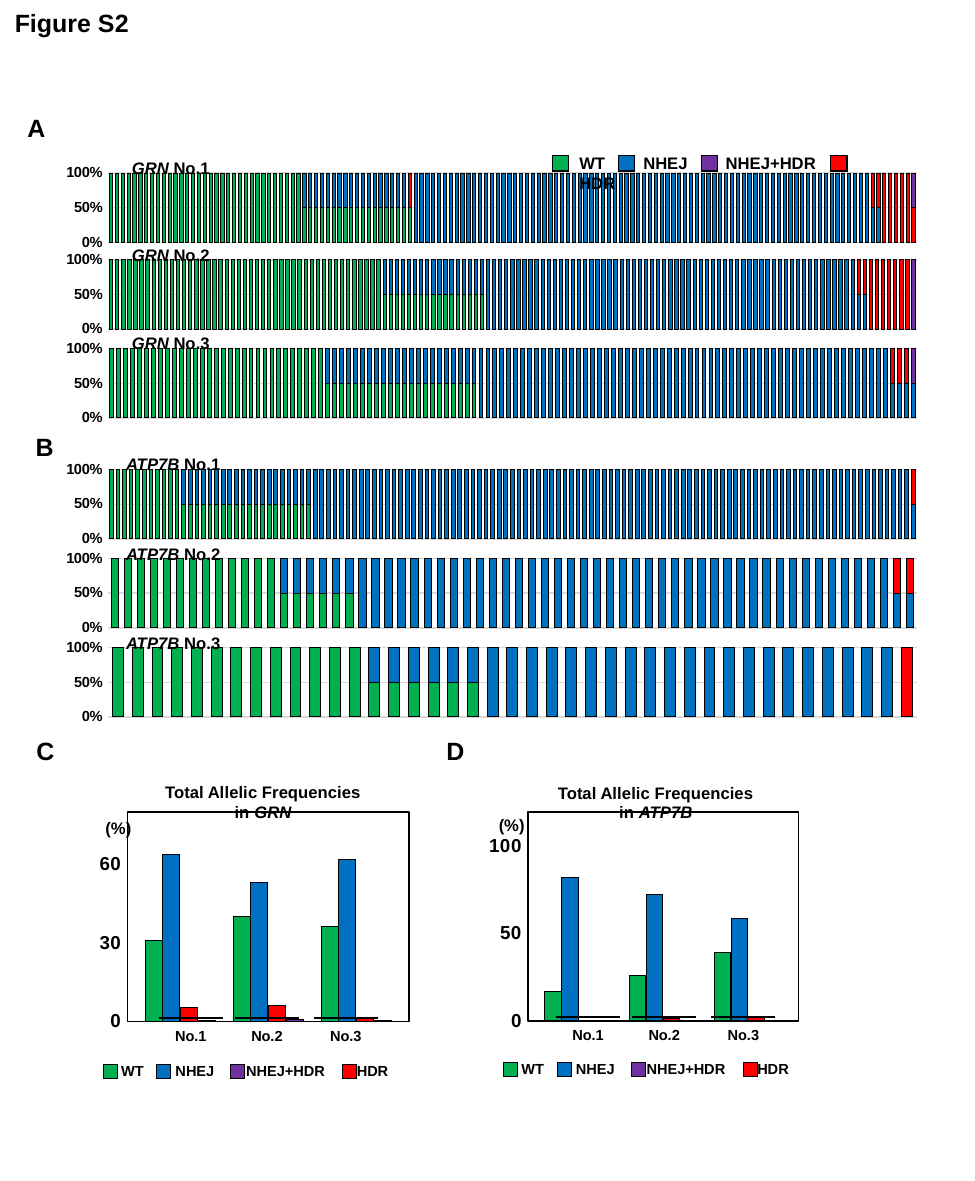

Figure S2
A
WT NHEJ NHEJ+HDR HDR
GRN No.1
#### Chart
| Category | GRN N2 | GRN N2 | GRN N2 | GRN N2 |
|---|---|---|---|---|
| 386 | 1.0 | 0.0 | 0.0 | 0.0 |
| 393 | 1.0 | 0.0 | 0.0 | 0.0 |
| 401 | 1.0 | 0.0 | 0.0 | 0.0 |
| 402 | 1.0 | 0.0 | 0.0 | 0.0 |
| 404 | 1.0 | 0.0 | 0.0 | 0.0 |
| 408 | 1.0 | 0.0 | 0.0 | 0.0 |
| 411 | 1.0 | 0.0 | 0.0 | 0.0 |
| 411 | 1.0 | 0.0 | 0.0 | 0.0 |
| 411 | 1.0 | 0.0 | 0.0 | 0.0 |
| 411 | 1.0 | 0.0 | 0.0 | 0.0 |
| 411 | 1.0 | 0.0 | 0.0 | 0.0 |
| 411 | 1.0 | 0.0 | 0.0 | 0.0 |
| 411 | 1.0 | 0.0 | 0.0 | 0.0 |
| 411 | 1.0 | 0.0 | 0.0 | 0.0 |
| 411 | 1.0 | 0.0 | 0.0 | 0.0 |
| 411 | 1.0 | 0.0 | 0.0 | 0.0 |
| 411 | 1.0 | 0.0 | 0.0 | 0.0 |
| 411 | 1.0 | 0.0 | 0.0 | 0.0 |
| 411 | 1.0 | 0.0 | 0.0 | 0.0 |
| 411 | 1.0 | 0.0 | 0.0 | 0.0 |
| 411 | 1.0 | 0.0 | 0.0 | 0.0 |
| 411 | 1.0 | 0.0 | 0.0 | 0.0 |
| 411 | 1.0 | 0.0 | 0.0 | 0.0 |
| 411 | 1.0 | 0.0 | 0.0 | 0.0 |
| 411 | 1.0 | 0.0 | 0.0 | 0.0 |
| 411 | 1.0 | 0.0 | 0.0 | 0.0 |
| 411 | 1.0 | 0.0 | 0.0 | 0.0 |
| 411 | 1.0 | 0.0 | 0.0 | 0.0 |
| 411 | 1.0 | 0.0 | 0.0 | 0.0 |
| 411 | 1.0 | 0.0 | 0.0 | 0.0 |
| 411 | 1.0 | 0.0 | 0.0 | 0.0 |
| 411 | 1.0 | 0.0 | 0.0 | 0.0 |
| 411 | 1.0 | 0.0 | 0.0 | 0.0 |
| 411 | 0.5 | 0.5 | 0.0 | 0.0 |
| 411 | 0.5 | 0.5 | 0.0 | 0.0 |
| 411 | 0.5 | 0.5 | 0.0 | 0.0 |
| 411 | 0.5 | 0.5 | 0.0 | 0.0 |
| 411 | 0.5 | 0.5 | 0.0 | 0.0 |
| 411 | 0.5 | 0.5 | 0.0 | 0.0 |
| 411 | 0.5 | 0.5 | 0.0 | 0.0 |
| 411 | 0.5 | 0.5 | 0.0 | 0.0 |
| 411 | 0.5 | 0.5 | 0.0 | 0.0 |
| 411 | 0.5 | 0.5 | 0.0 | 0.0 |
| 411 | 0.5 | 0.5 | 0.0 | 0.0 |
| 411 | 0.5 | 0.5 | 0.0 | 0.0 |
| 411 | 0.5 | 0.5 | 0.0 | 0.0 |
| 411 | 0.5 | 0.5 | 0.0 | 0.0 |
| 411 | 0.5 | 0.5 | 0.0 | 0.0 |
| 411 | 0.5 | 0.5 | 0.0 | 0.0 |
| 411 | 0.5 | 0.5 | 0.0 | 0.0 |
| 411 | 0.5 | 0.5 | 0.0 | 0.0 |
| 411 | 0.5 | 0.0 | 0.5 | 0.0 |
| 411 | 0.0 | 1.0 | 0.0 | 0.0 |
| 411 | 0.0 | 1.0 | 0.0 | 0.0 |
| 411 | 0.0 | 1.0 | 0.0 | 0.0 |
| 411 | 0.0 | 1.0 | 0.0 | 0.0 |
| 411 | 0.0 | 1.0 | 0.0 | 0.0 |
| 411 | 0.0 | 1.0 | 0.0 | 0.0 |
| 411 | 0.0 | 1.0 | 0.0 | 0.0 |
| 411 | 0.0 | 1.0 | 0.0 | 0.0 |
| 411 | 0.0 | 1.0 | 0.0 | 0.0 |
| 411 | 0.0 | 1.0 | 0.0 | 0.0 |
| 411 | 0.0 | 1.0 | 0.0 | 0.0 |
| 411 | 0.0 | 1.0 | 0.0 | 0.0 |
| 411 | 0.0 | 1.0 | 0.0 | 0.0 |
| 411 | 0.0 | 1.0 | 0.0 | 0.0 |
| 411 | 0.0 | 1.0 | 0.0 | 0.0 |
| 411 | 0.0 | 1.0 | 0.0 | 0.0 |
| 411 | 0.0 | 1.0 | 0.0 | 0.0 |
| 411 | 0.0 | 1.0 | 0.0 | 0.0 |
| 411 | 0.0 | 1.0 | 0.0 | 0.0 |
| 411 | 0.0 | 1.0 | 0.0 | 0.0 |
| 411 | 0.0 | 1.0 | 0.0 | 0.0 |
| 411 | 0.0 | 1.0 | 0.0 | 0.0 |
| 411 | 0.0 | 1.0 | 0.0 | 0.0 |
| 411 | 0.0 | 1.0 | 0.0 | 0.0 |
| 411 | 0.0 | 1.0 | 0.0 | 0.0 |
| 411 | 0.0 | 1.0 | 0.0 | 0.0 |
| 411 | 0.0 | 1.0 | 0.0 | 0.0 |
| 411 | 0.0 | 1.0 | 0.0 | 0.0 |
| 411 | 0.0 | 1.0 | 0.0 | 0.0 |
| 411 | 0.0 | 1.0 | 0.0 | 0.0 |
| 411 | 0.0 | 1.0 | 0.0 | 0.0 |
| 411 | 0.0 | 1.0 | 0.0 | 0.0 |
| 411 | 0.0 | 1.0 | 0.0 | 0.0 |
| 411 | 0.0 | 1.0 | 0.0 | 0.0 |
| 411 | 0.0 | 1.0 | 0.0 | 0.0 |
| 411 | 0.0 | 1.0 | 0.0 | 0.0 |
| 411 | 0.0 | 1.0 | 0.0 | 0.0 |
| 411 | 0.0 | 1.0 | 0.0 | 0.0 |
| 411 | 0.0 | 1.0 | 0.0 | 0.0 |
| 411 | 0.0 | 1.0 | 0.0 | 0.0 |
| 411 | 0.0 | 1.0 | 0.0 | 0.0 |
| 411 | 0.0 | 1.0 | 0.0 | 0.0 |
| 411 | 0.0 | 1.0 | 0.0 | 0.0 |
| 411 | 0.0 | 1.0 | 0.0 | 0.0 |
| 411 | 0.0 | 1.0 | 0.0 | 0.0 |
| 411 | 0.0 | 1.0 | 0.0 | 0.0 |
| 411 | 0.0 | 1.0 | 0.0 | 0.0 |
| 411 | 0.0 | 1.0 | 0.0 | 0.0 |
| 411 | 0.0 | 1.0 | 0.0 | 0.0 |
| 411 | 0.0 | 1.0 | 0.0 | 0.0 |
| 411 | 0.0 | 1.0 | 0.0 | 0.0 |
| 411 | 0.0 | 1.0 | 0.0 | 0.0 |
| 411 | 0.0 | 1.0 | 0.0 | 0.0 |
| 411 | 0.0 | 1.0 | 0.0 | 0.0 |
| 411 | 0.0 | 1.0 | 0.0 | 0.0 |
| 411 | 0.0 | 1.0 | 0.0 | 0.0 |
| 411 | 0.0 | 1.0 | 0.0 | 0.0 |
| 411 | 0.0 | 1.0 | 0.0 | 0.0 |
| 411 | 0.0 | 1.0 | 0.0 | 0.0 |
| 411 | 0.0 | 1.0 | 0.0 | 0.0 |
| 411 | 0.0 | 1.0 | 0.0 | 0.0 |
| 411 | 0.0 | 1.0 | 0.0 | 0.0 |
| 411 | 0.0 | 1.0 | 0.0 | 0.0 |
| 411 | 0.0 | 1.0 | 0.0 | 0.0 |
| 411 | 0.0 | 1.0 | 0.0 | 0.0 |
| 411 | 0.0 | 1.0 | 0.0 | 0.0 |
| 411 | 0.0 | 1.0 | 0.0 | 0.0 |
| 411 | 0.0 | 1.0 | 0.0 | 0.0 |
| 411 | 0.0 | 1.0 | 0.0 | 0.0 |
| 411 | 0.0 | 1.0 | 0.0 | 0.0 |
| 411 | 0.0 | 1.0 | 0.0 | 0.0 |
| 411 | 0.0 | 1.0 | 0.0 | 0.0 |
| 411 | 0.0 | 1.0 | 0.0 | 0.0 |
| 411 | 0.0 | 1.0 | 0.0 | 0.0 |
| 411 | 0.0 | 1.0 | 0.0 | 0.0 |
| 411 | 0.0 | 1.0 | 0.0 | 0.0 |
| 411 | 0.0 | 1.0 | 0.0 | 0.0 |
| 411 | 0.0 | 1.0 | 0.0 | 0.0 |
| 411 | 0.0 | 0.5 | 0.5 | 0.0 |
| 411 | 0.0 | 0.5 | 0.5 | 0.0 |
| 411 | 0.0 | 0.0 | 1.0 | 0.0 |
| 411 | 0.0 | 0.0 | 1.0 | 0.0 |
| 411 | 0.0 | 0.0 | 1.0 | 0.0 |
| 411 | 0.0 | 0.0 | 1.0 | 0.0 |
| 411 | 0.0 | 0.0 | 1.0 | 0.0 |
| 411 | 0.0 | 0.0 | 0.5 | 0.5 |GRN No.2
#### Chart
| Category | GRN N3 | GRN N3 | GRN N3 | GRN N3 |
|---|---|---|---|---|
| 544 | 1.0 | 0.0 | 0.0 | 0.0 |
| 549 | 1.0 | 0.0 | 0.0 | 0.0 |
| 550 | 1.0 | 0.0 | 0.0 | 0.0 |
| 551 | 1.0 | 0.0 | 0.0 | 0.0 |
| 552 | 1.0 | 0.0 | 0.0 | 0.0 |
| 554 | 1.0 | 0.0 | 0.0 | 0.0 |
| 556 | 1.0 | 0.0 | 0.0 | 0.0 |
| 559 | 1.0 | 0.0 | 0.0 | 0.0 |
| 560 | 1.0 | 0.0 | 0.0 | 0.0 |
| 562 | 1.0 | 0.0 | 0.0 | 0.0 |
| 563 | 1.0 | 0.0 | 0.0 | 0.0 |
| 569 | 1.0 | 0.0 | 0.0 | 0.0 |
| 574 | 1.0 | 0.0 | 0.0 | 0.0 |
| 575 | 1.0 | 0.0 | 0.0 | 0.0 |
| 581 | 1.0 | 0.0 | 0.0 | 0.0 |
| 582 | 1.0 | 0.0 | 0.0 | 0.0 |
| 587 | 1.0 | 0.0 | 0.0 | 0.0 |
| 595 | 1.0 | 0.0 | 0.0 | 0.0 |
| 596 | 1.0 | 0.0 | 0.0 | 0.0 |
| 597 | 1.0 | 0.0 | 0.0 | 0.0 |
| 602 | 1.0 | 0.0 | 0.0 | 0.0 |
| 607 | 1.0 | 0.0 | 0.0 | 0.0 |
| 609 | 1.0 | 0.0 | 0.0 | 0.0 |
| 611 | 1.0 | 0.0 | 0.0 | 0.0 |
| 614 | 1.0 | 0.0 | 0.0 | 0.0 |
| 617 | 1.0 | 0.0 | 0.0 | 0.0 |
| 618 | 1.0 | 0.0 | 0.0 | 0.0 |
| 619 | 1.0 | 0.0 | 0.0 | 0.0 |
| 625 | 1.0 | 0.0 | 0.0 | 0.0 |
| 627 | 1.0 | 0.0 | 0.0 | 0.0 |
| 634 | 1.0 | 0.0 | 0.0 | 0.0 |
| 636 | 1.0 | 0.0 | 0.0 | 0.0 |
| 640 | 1.0 | 0.0 | 0.0 | 0.0 |
| 641 | 1.0 | 0.0 | 0.0 | 0.0 |
| 643 | 1.0 | 0.0 | 0.0 | 0.0 |
| 647 | 1.0 | 0.0 | 0.0 | 0.0 |
| 648 | 1.0 | 0.0 | 0.0 | 0.0 |
| 655 | 1.0 | 0.0 | 0.0 | 0.0 |
| 659 | 1.0 | 0.0 | 0.0 | 0.0 |
| 660 | 1.0 | 0.0 | 0.0 | 0.0 |
| 674 | 1.0 | 0.0 | 0.0 | 0.0 |
| 688 | 1.0 | 0.0 | 0.0 | 0.0 |
| 691 | 1.0 | 0.0 | 0.0 | 0.0 |
| 692 | 1.0 | 0.0 | 0.0 | 0.0 |
| 696 | 1.0 | 0.0 | 0.0 | 0.0 |
| 579 | 0.5 | 0.5 | 0.0 | 0.0 |
| 580 | 0.5 | 0.5 | 0.0 | 0.0 |
| 603 | 0.5 | 0.5 | 0.0 | 0.0 |
| 644 | 0.5 | 0.5 | 0.0 | 0.0 |
| 545 | 0.5 | 0.5 | 0.0 | 0.0 |
| 561 | 0.5 | 0.5 | 0.0 | 0.0 |
| 604 | 0.5 | 0.5 | 0.0 | 0.0 |
| 646 | 0.5 | 0.5 | 0.0 | 0.0 |
| 664 | 0.5 | 0.5 | 0.0 | 0.0 |
| 694 | 0.5 | 0.5 | 0.0 | 0.0 |
| 573 | 0.5 | 0.5 | 0.0 | 0.0 |
| 576 | 0.5 | 0.5 | 0.0 | 0.0 |
| 623 | 0.5 | 0.5 | 0.0 | 0.0 |
| 632 | 0.5 | 0.5 | 0.0 | 0.0 |
| 652 | 0.5 | 0.5 | 0.0 | 0.0 |
| 678 | 0.5 | 0.5 | 0.0 | 0.0 |
| 693 | 0.5 | 0.5 | 0.0 | 0.0 |
| 540 | 0.0 | 1.0 | 0.0 | 0.0 |
| 541 | 0.0 | 1.0 | 0.0 | 0.0 |
| 542 | 0.0 | 1.0 | 0.0 | 0.0 |
| 543 | 0.0 | 1.0 | 0.0 | 0.0 |
| 546 | 0.0 | 1.0 | 0.0 | 0.0 |
| 553 | 0.0 | 1.0 | 0.0 | 0.0 |
| 555 | 0.0 | 1.0 | 0.0 | 0.0 |
| 557 | 0.0 | 1.0 | 0.0 | 0.0 |
| 558 | 0.0 | 1.0 | 0.0 | 0.0 |
| 564 | 0.0 | 1.0 | 0.0 | 0.0 |
| 565 | 0.0 | 1.0 | 0.0 | 0.0 |
| 566 | 0.0 | 1.0 | 0.0 | 0.0 |
| 567 | 0.0 | 1.0 | 0.0 | 0.0 |
| 568 | 0.0 | 1.0 | 0.0 | 0.0 |
| 570 | 0.0 | 1.0 | 0.0 | 0.0 |
| 571 | 0.0 | 1.0 | 0.0 | 0.0 |
| 572 | 0.0 | 1.0 | 0.0 | 0.0 |
| 577 | 0.0 | 1.0 | 0.0 | 0.0 |
| 578 | 0.0 | 1.0 | 0.0 | 0.0 |
| 586 | 0.0 | 1.0 | 0.0 | 0.0 |
| 588 | 0.0 | 1.0 | 0.0 | 0.0 |
| 593 | 0.0 | 1.0 | 0.0 | 0.0 |
| 594 | 0.0 | 1.0 | 0.0 | 0.0 |
| 598 | 0.0 | 1.0 | 0.0 | 0.0 |
| 601 | 0.0 | 1.0 | 0.0 | 0.0 |
| 608 | 0.0 | 1.0 | 0.0 | 0.0 |
| 616 | 0.0 | 1.0 | 0.0 | 0.0 |
| 621 | 0.0 | 1.0 | 0.0 | 0.0 |
| 622 | 0.0 | 1.0 | 0.0 | 0.0 |
| 624 | 0.0 | 1.0 | 0.0 | 0.0 |
| 628 | 0.0 | 1.0 | 0.0 | 0.0 |
| 629 | 0.0 | 1.0 | 0.0 | 0.0 |
| 633 | 0.0 | 1.0 | 0.0 | 0.0 |
| 635 | 0.0 | 1.0 | 0.0 | 0.0 |
| 637 | 0.0 | 1.0 | 0.0 | 0.0 |
| 638 | 0.0 | 1.0 | 0.0 | 0.0 |
| 649 | 0.0 | 1.0 | 0.0 | 0.0 |
| 650 | 0.0 | 1.0 | 0.0 | 0.0 |
| 651 | 0.0 | 1.0 | 0.0 | 0.0 |
| 653 | 0.0 | 1.0 | 0.0 | 0.0 |
| 654 | 0.0 | 1.0 | 0.0 | 0.0 |
| 656 | 0.0 | 1.0 | 0.0 | 0.0 |
| 657 | 0.0 | 1.0 | 0.0 | 0.0 |
| 658 | 0.0 | 1.0 | 0.0 | 0.0 |
| 661 | 0.0 | 1.0 | 0.0 | 0.0 |
| 662 | 0.0 | 1.0 | 0.0 | 0.0 |
| 666 | 0.0 | 1.0 | 0.0 | 0.0 |
| 669 | 0.0 | 1.0 | 0.0 | 0.0 |
| 670 | 0.0 | 1.0 | 0.0 | 0.0 |
| 672 | 0.0 | 1.0 | 0.0 | 0.0 |
| 673 | 0.0 | 1.0 | 0.0 | 0.0 |
| 675 | 0.0 | 1.0 | 0.0 | 0.0 |
| 676 | 0.0 | 1.0 | 0.0 | 0.0 |
| 677 | 0.0 | 1.0 | 0.0 | 0.0 |
| 681 | 0.0 | 1.0 | 0.0 | 0.0 |
| 683 | 0.0 | 1.0 | 0.0 | 0.0 |
| 684 | 0.0 | 1.0 | 0.0 | 0.0 |
| 685 | 0.0 | 1.0 | 0.0 | 0.0 |
| 689 | 0.0 | 1.0 | 0.0 | 0.0 |
| 690 | 0.0 | 1.0 | 0.0 | 0.0 |
| 698 | 0.0 | 1.0 | 0.0 | 0.0 |
| 548 | 0.0 | 0.5 | 0.5 | 0.0 |
| 585 | 0.0 | 0.5 | 0.5 | 0.0 |
| 589 | 0.0 | 0.0 | 1.0 | 0.0 |
| 613 | 0.0 | 0.0 | 1.0 | 0.0 |
| 620 | 0.0 | 0.0 | 1.0 | 0.0 |
| 626 | 0.0 | 0.0 | 1.0 | 0.0 |
| 642 | 0.0 | 0.0 | 1.0 | 0.0 |
| 665 | 0.0 | 0.0 | 1.0 | 0.0 |
| 667 | 0.0 | 0.0 | 1.0 | 0.0 |
| 605 | 0.0 | 0.0 | 0.0 | 1.0 |GRN No.3
#### Chart
| Category | GRN N4 | GRN N4 | GRN N4 | GRN N4 |
|---|---|---|---|---|
| 719 | 1.0 | 0.0 | 0.0 | 0.0 |
| 727 | 1.0 | 0.0 | 0.0 | 0.0 |
| 793 | 1.0 | 0.0 | 0.0 | 0.0 |
| 781 | 1.0 | 0.0 | 0.0 | 0.0 |
| 770 | 1.0 | 0.0 | 0.0 | 0.0 |
| 762 | 1.0 | 0.0 | 0.0 | 0.0 |
| 728 | 1.0 | 0.0 | 0.0 | 0.0 |
| 810 | 1.0 | 0.0 | 0.0 | 0.0 |
| 733 | 1.0 | 0.0 | 0.0 | 0.0 |
| 732 | 1.0 | 0.0 | 0.0 | 0.0 |
| 730 | 1.0 | 0.0 | 0.0 | 0.0 |
| 760 | 1.0 | 0.0 | 0.0 | 0.0 |
| 711 | 1.0 | 0.0 | 0.0 | 0.0 |
| 756 | 1.0 | 0.0 | 0.0 | 0.0 |
| 812 | 1.0 | 0.0 | 0.0 | 0.0 |
| 699 | 1.0 | 0.0 | 0.0 | 0.0 |
| 734 | 1.0 | 0.0 | 0.0 | 0.0 |
| 759 | 1.0 | 0.0 | 0.0 | 0.0 |
| 772 | 1.0 | 0.0 | 0.0 | 0.0 |
| 761 | 1.0 | 0.0 | 0.0 | 0.0 |
| 835 | 1.0 | 0.0 | 0.0 | 0.0 |
| 764 | 1.0 | 0.0 | 0.0 | 0.0 |
| 804 | 1.0 | 0.0 | 0.0 | 0.0 |
| 831 | 1.0 | 0.0 | 0.0 | 0.0 |
| 830 | 1.0 | 0.0 | 0.0 | 0.0 |
| 805 | 1.0 | 0.0 | 0.0 | 0.0 |
| 738 | 1.0 | 0.0 | 0.0 | 0.0 |
| 771 | 1.0 | 0.0 | 0.0 | 0.0 |
| 819 | 1.0 | 0.0 | 0.0 | 0.0 |
| 818 | 1.0 | 0.0 | 0.0 | 0.0 |
| 751 | 1.0 | 0.0 | 0.0 | 0.0 |
| 790 | 0.5 | 0.5 | 0.0 | 0.0 |
| 786 | 0.5 | 0.5 | 0.0 | 0.0 |
| 740 | 0.5 | 0.5 | 0.0 | 0.0 |
| 787 | 0.5 | 0.5 | 0.0 | 0.0 |
| 834 | 0.5 | 0.5 | 0.0 | 0.0 |
| 706 | 0.5 | 0.5 | 0.0 | 0.0 |
| 746 | 0.5 | 0.5 | 0.0 | 0.0 |
| 789 | 0.5 | 0.5 | 0.0 | 0.0 |
| 726 | 0.5 | 0.5 | 0.0 | 0.0 |
| 717 | 0.5 | 0.5 | 0.0 | 0.0 |
| 748 | 0.5 | 0.5 | 0.0 | 0.0 |
| 768 | 0.5 | 0.5 | 0.0 | 0.0 |
| 758 | 0.5 | 0.5 | 0.0 | 0.0 |
| 709 | 0.5 | 0.5 | 0.0 | 0.0 |
| 723 | 0.5 | 0.5 | 0.0 | 0.0 |
| 752 | 0.5 | 0.5 | 0.0 | 0.0 |
| 742 | 0.5 | 0.5 | 0.0 | 0.0 |
| 718 | 0.5 | 0.5 | 0.0 | 0.0 |
| 773 | 0.5 | 0.5 | 0.0 | 0.0 |
| 807 | 0.5 | 0.5 | 0.0 | 0.0 |
| 827 | 0.5 | 0.5 | 0.0 | 0.0 |
| 823 | 0.5 | 0.5 | 0.0 | 0.0 |
| 794 | 0.0 | 1.0 | 0.0 | 0.0 |
| 792 | 0.0 | 1.0 | 0.0 | 0.0 |
| 791 | 0.0 | 1.0 | 0.0 | 0.0 |
| 753 | 0.0 | 1.0 | 0.0 | 0.0 |
| 715 | 0.0 | 1.0 | 0.0 | 0.0 |
| 788 | 0.0 | 1.0 | 0.0 | 0.0 |
| 750 | 0.0 | 1.0 | 0.0 | 0.0 |
| 724 | 0.0 | 1.0 | 0.0 | 0.0 |
| 741 | 0.0 | 1.0 | 0.0 | 0.0 |
| 815 | 0.0 | 1.0 | 0.0 | 0.0 |
| 774 | 0.0 | 1.0 | 0.0 | 0.0 |
| 754 | 0.0 | 1.0 | 0.0 | 0.0 |
| 735 | 0.0 | 1.0 | 0.0 | 0.0 |
| 767 | 0.0 | 1.0 | 0.0 | 0.0 |
| 796 | 0.0 | 1.0 | 0.0 | 0.0 |
| 749 | 0.0 | 1.0 | 0.0 | 0.0 |
| 744 | 0.0 | 1.0 | 0.0 | 0.0 |
| 778 | 0.0 | 1.0 | 0.0 | 0.0 |
| 747 | 0.0 | 1.0 | 0.0 | 0.0 |
| 784 | 0.0 | 1.0 | 0.0 | 0.0 |
| 708 | 0.0 | 1.0 | 0.0 | 0.0 |
| 755 | 0.0 | 1.0 | 0.0 | 0.0 |
| 725 | 0.0 | 1.0 | 0.0 | 0.0 |
| 714 | 0.0 | 1.0 | 0.0 | 0.0 |
| 775 | 0.0 | 1.0 | 0.0 | 0.0 |
| 785 | 0.0 | 1.0 | 0.0 | 0.0 |
| 776 | 0.0 | 1.0 | 0.0 | 0.0 |
| 710 | 0.0 | 1.0 | 0.0 | 0.0 |
| 701 | 0.0 | 1.0 | 0.0 | 0.0 |
| 700 | 0.0 | 1.0 | 0.0 | 0.0 |
| 765 | 0.0 | 1.0 | 0.0 | 0.0 |
| 779 | 0.0 | 1.0 | 0.0 | 0.0 |
| 766 | 0.0 | 1.0 | 0.0 | 0.0 |
| 803 | 0.0 | 1.0 | 0.0 | 0.0 |
| 806 | 0.0 | 1.0 | 0.0 | 0.0 |
| 798 | 0.0 | 1.0 | 0.0 | 0.0 |
| 712 | 0.0 | 1.0 | 0.0 | 0.0 |
| 783 | 0.0 | 1.0 | 0.0 | 0.0 |
| 814 | 0.0 | 1.0 | 0.0 | 0.0 |
| 813 | 0.0 | 1.0 | 0.0 | 0.0 |
| 795 | 0.0 | 1.0 | 0.0 | 0.0 |
| 797 | 0.0 | 1.0 | 0.0 | 0.0 |
| 763 | 0.0 | 1.0 | 0.0 | 0.0 |
| 703 | 0.0 | 1.0 | 0.0 | 0.0 |
| 716 | 0.0 | 1.0 | 0.0 | 0.0 |
| 757 | 0.0 | 1.0 | 0.0 | 0.0 |
| 720 | 0.0 | 1.0 | 0.0 | 0.0 |
| 811 | 0.0 | 1.0 | 0.0 | 0.0 |
| 822 | 0.0 | 1.0 | 0.0 | 0.0 |
| 729 | 0.0 | 1.0 | 0.0 | 0.0 |
| 745 | 0.0 | 1.0 | 0.0 | 0.0 |
| 828 | 0.0 | 1.0 | 0.0 | 0.0 |
| 829 | 0.0 | 1.0 | 0.0 | 0.0 |
| 769 | 0.0 | 1.0 | 0.0 | 0.0 |
| 780 | 0.0 | 1.0 | 0.0 | 0.0 |
| 707 | 0.0 | 1.0 | 0.0 | 0.0 |
| 702 | 0.0 | 1.0 | 0.0 | 0.0 |
| 817 | 0.0 | 1.0 | 0.0 | 0.0 |
| 826 | 0.0 | 1.0 | 0.0 | 0.0 |
| 782 | 0.0 | 0.5 | 0.5 | 0.0 |
| 743 | 0.0 | 0.5 | 0.5 | 0.0 |
| 731 | 0.0 | 0.5 | 0.5 | 0.0 |
| 777 | 0.0 | 0.5 | 0.0 | 0.5 |B
ATP7B No.1
#### Chart
| Category | ATP7B N1 | ATP7B N1 | ATP7B N1 | ATP7B N1 |
|---|---|---|---|---|
| 9 | 1.0 | 0.0 | 0.0 | 0.0 |
| 12 | 1.0 | 0.0 | 0.0 | 0.0 |
| 57 | 1.0 | 0.0 | 0.0 | 0.0 |
| 60 | 1.0 | 0.0 | 0.0 | 0.0 |
| 81 | 1.0 | 0.0 | 0.0 | 0.0 |
| 140 | 1.0 | 0.0 | 0.0 | 0.0 |
| 153 | 1.0 | 0.0 | 0.0 | 0.0 |
| 169 | 1.0 | 0.0 | 0.0 | 0.0 |
| 172 | 1.0 | 0.0 | 0.0 | 0.0 |
| 175 | 1.0 | 0.0 | 0.0 | 0.0 |
| 179 | 1.0 | 0.0 | 0.0 | 0.0 |
| 10 | 0.5 | 0.5 | 0.0 | 0.0 |
| 17 | 0.5 | 0.5 | 0.0 | 0.0 |
| 53 | 0.5 | 0.5 | 0.0 | 0.0 |
| 118 | 0.5 | 0.5 | 0.0 | 0.0 |
| 19 | 0.5 | 0.5 | 0.0 | 0.0 |
| 25 | 0.5 | 0.5 | 0.0 | 0.0 |
| 45 | 0.5 | 0.5 | 0.0 | 0.0 |
| 52 | 0.5 | 0.5 | 0.0 | 0.0 |
| 109 | 0.5 | 0.5 | 0.0 | 0.0 |
| 142 | 0.5 | 0.5 | 0.0 | 0.0 |
| 166 | 0.5 | 0.5 | 0.0 | 0.0 |
| 178 | 0.5 | 0.5 | 0.0 | 0.0 |
| 1 | 0.5 | 0.5 | 0.0 | 0.0 |
| 11 | 0.5 | 0.5 | 0.0 | 0.0 |
| 75 | 0.5 | 0.5 | 0.0 | 0.0 |
| 84 | 0.5 | 0.5 | 0.0 | 0.0 |
| 97 | 0.5 | 0.5 | 0.0 | 0.0 |
| 105 | 0.5 | 0.5 | 0.0 | 0.0 |
| 161 | 0.5 | 0.5 | 0.0 | 0.0 |
| 173 | 0.5 | 0.5 | 0.0 | 0.0 |
| 2 | 0.0 | 1.0 | 0.0 | 0.0 |
| 4 | 0.0 | 1.0 | 0.0 | 0.0 |
| 13 | 0.0 | 1.0 | 0.0 | 0.0 |
| 14 | 0.0 | 1.0 | 0.0 | 0.0 |
| 15 | 0.0 | 1.0 | 0.0 | 0.0 |
| 20 | 0.0 | 1.0 | 0.0 | 0.0 |
| 21 | 0.0 | 1.0 | 0.0 | 0.0 |
| 22 | 0.0 | 1.0 | 0.0 | 0.0 |
| 27 | 0.0 | 1.0 | 0.0 | 0.0 |
| 29 | 0.0 | 1.0 | 0.0 | 0.0 |
| 32 | 0.0 | 1.0 | 0.0 | 0.0 |
| 33 | 0.0 | 1.0 | 0.0 | 0.0 |
| 34 | 0.0 | 1.0 | 0.0 | 0.0 |
| 35 | 0.0 | 1.0 | 0.0 | 0.0 |
| 36 | 0.0 | 1.0 | 0.0 | 0.0 |
| 37 | 0.0 | 1.0 | 0.0 | 0.0 |
| 42 | 0.0 | 1.0 | 0.0 | 0.0 |
| 43 | 0.0 | 1.0 | 0.0 | 0.0 |
| 46 | 0.0 | 1.0 | 0.0 | 0.0 |
| 47 | 0.0 | 1.0 | 0.0 | 0.0 |
| 48 | 0.0 | 1.0 | 0.0 | 0.0 |
| 49 | 0.0 | 1.0 | 0.0 | 0.0 |
| 50 | 0.0 | 1.0 | 0.0 | 0.0 |
| 51 | 0.0 | 1.0 | 0.0 | 0.0 |
| 55 | 0.0 | 1.0 | 0.0 | 0.0 |
| 56 | 0.0 | 1.0 | 0.0 | 0.0 |
| 58 | 0.0 | 1.0 | 0.0 | 0.0 |
| 59 | 0.0 | 1.0 | 0.0 | 0.0 |
| 61 | 0.0 | 1.0 | 0.0 | 0.0 |
| 62 | 0.0 | 1.0 | 0.0 | 0.0 |
| 63 | 0.0 | 1.0 | 0.0 | 0.0 |
| 64 | 0.0 | 1.0 | 0.0 | 0.0 |
| 65 | 0.0 | 1.0 | 0.0 | 0.0 |
| 66 | 0.0 | 1.0 | 0.0 | 0.0 |
| 68 | 0.0 | 1.0 | 0.0 | 0.0 |
| 69 | 0.0 | 1.0 | 0.0 | 0.0 |
| 71 | 0.0 | 1.0 | 0.0 | 0.0 |
| 72 | 0.0 | 1.0 | 0.0 | 0.0 |
| 73 | 0.0 | 1.0 | 0.0 | 0.0 |
| 77 | 0.0 | 1.0 | 0.0 | 0.0 |
| 79 | 0.0 | 1.0 | 0.0 | 0.0 |
| 83 | 0.0 | 1.0 | 0.0 | 0.0 |
| 86 | 0.0 | 1.0 | 0.0 | 0.0 |
| 87 | 0.0 | 1.0 | 0.0 | 0.0 |
| 88 | 0.0 | 1.0 | 0.0 | 0.0 |
| 89 | 0.0 | 1.0 | 0.0 | 0.0 |
| 96 | 0.0 | 1.0 | 0.0 | 0.0 |
| 101 | 0.0 | 1.0 | 0.0 | 0.0 |
| 102 | 0.0 | 1.0 | 0.0 | 0.0 |
| 106 | 0.0 | 1.0 | 0.0 | 0.0 |
| 107 | 0.0 | 1.0 | 0.0 | 0.0 |
| 108 | 0.0 | 1.0 | 0.0 | 0.0 |
| 110 | 0.0 | 1.0 | 0.0 | 0.0 |
| 111 | 0.0 | 1.0 | 0.0 | 0.0 |
| 113 | 0.0 | 1.0 | 0.0 | 0.0 |
| 114 | 0.0 | 1.0 | 0.0 | 0.0 |
| 117 | 0.0 | 1.0 | 0.0 | 0.0 |
| 121 | 0.0 | 1.0 | 0.0 | 0.0 |
| 122 | 0.0 | 1.0 | 0.0 | 0.0 |
| 123 | 0.0 | 1.0 | 0.0 | 0.0 |
| 125 | 0.0 | 1.0 | 0.0 | 0.0 |
| 126 | 0.0 | 1.0 | 0.0 | 0.0 |
| 128 | 0.0 | 1.0 | 0.0 | 0.0 |
| 129 | 0.0 | 1.0 | 0.0 | 0.0 |
| 130 | 0.0 | 1.0 | 0.0 | 0.0 |
| 131 | 0.0 | 1.0 | 0.0 | 0.0 |
| 132 | 0.0 | 1.0 | 0.0 | 0.0 |
| 133 | 0.0 | 1.0 | 0.0 | 0.0 |
| 134 | 0.0 | 1.0 | 0.0 | 0.0 |
| 135 | 0.0 | 1.0 | 0.0 | 0.0 |
| 136 | 0.0 | 1.0 | 0.0 | 0.0 |
| 137 | 0.0 | 1.0 | 0.0 | 0.0 |
| 138 | 0.0 | 1.0 | 0.0 | 0.0 |
| 139 | 0.0 | 1.0 | 0.0 | 0.0 |
| 141 | 0.0 | 1.0 | 0.0 | 0.0 |
| 143 | 0.0 | 1.0 | 0.0 | 0.0 |
| 144 | 0.0 | 1.0 | 0.0 | 0.0 |
| 145 | 0.0 | 1.0 | 0.0 | 0.0 |
| 147 | 0.0 | 1.0 | 0.0 | 0.0 |
| 152 | 0.0 | 1.0 | 0.0 | 0.0 |
| 154 | 0.0 | 1.0 | 0.0 | 0.0 |
| 155 | 0.0 | 1.0 | 0.0 | 0.0 |
| 157 | 0.0 | 1.0 | 0.0 | 0.0 |
| 160 | 0.0 | 1.0 | 0.0 | 0.0 |
| 162 | 0.0 | 1.0 | 0.0 | 0.0 |
| 163 | 0.0 | 1.0 | 0.0 | 0.0 |
| 165 | 0.0 | 1.0 | 0.0 | 0.0 |
| 167 | 0.0 | 1.0 | 0.0 | 0.0 |
| 168 | 0.0 | 1.0 | 0.0 | 0.0 |
| 171 | 0.0 | 1.0 | 0.0 | 0.0 |
| 177 | 0.0 | 1.0 | 0.0 | 0.0 |
| 38 | 0.0 | 0.5 | 0.5 | 0.0 |ATP7B No.2
#### Chart
| Category | ATP7B N2 | ATP7B N2 | ATP7B N2 | ATP7B N2 |
|---|---|---|---|---|
| 186 | 1.0 | 0.0 | 0.0 | 0.0 |
| 192 | 1.0 | 0.0 | 0.0 | 0.0 |
| 193 | 1.0 | 0.0 | 0.0 | 0.0 |
| 202 | 1.0 | 0.0 | 0.0 | 0.0 |
| 221 | 1.0 | 0.0 | 0.0 | 0.0 |
| 229 | 1.0 | 0.0 | 0.0 | 0.0 |
| 233 | 1.0 | 0.0 | 0.0 | 0.0 |
| 235 | 1.0 | 0.0 | 0.0 | 0.0 |
| 239 | 1.0 | 0.0 | 0.0 | 0.0 |
| 253 | 1.0 | 0.0 | 0.0 | 0.0 |
| 258 | 1.0 | 0.0 | 0.0 | 0.0 |
| 259 | 1.0 | 0.0 | 0.0 | 0.0 |
| 264 | 1.0 | 0.0 | 0.0 | 0.0 |
| 274 | 0.5 | 0.5 | 0.0 | 0.0 |
| 240 | 0.5 | 0.5 | 0.0 | 0.0 |
| 260 | 0.5 | 0.5 | 0.0 | 0.0 |
| 282 | 0.5 | 0.5 | 0.0 | 0.0 |
| 223 | 0.5 | 0.5 | 0.0 | 0.0 |
| 232 | 0.5 | 0.5 | 0.0 | 0.0 |
| 182 | 0.0 | 1.0 | 0.0 | 0.0 |
| 184 | 0.0 | 1.0 | 0.0 | 0.0 |
| 185 | 0.0 | 1.0 | 0.0 | 0.0 |
| 187 | 0.0 | 1.0 | 0.0 | 0.0 |
| 189 | 0.0 | 1.0 | 0.0 | 0.0 |
| 197 | 0.0 | 1.0 | 0.0 | 0.0 |
| 198 | 0.0 | 1.0 | 0.0 | 0.0 |
| 201 | 0.0 | 1.0 | 0.0 | 0.0 |
| 208 | 0.0 | 1.0 | 0.0 | 0.0 |
| 209 | 0.0 | 1.0 | 0.0 | 0.0 |
| 211 | 0.0 | 1.0 | 0.0 | 0.0 |
| 212 | 0.0 | 1.0 | 0.0 | 0.0 |
| 217 | 0.0 | 1.0 | 0.0 | 0.0 |
| 218 | 0.0 | 1.0 | 0.0 | 0.0 |
| 220 | 0.0 | 1.0 | 0.0 | 0.0 |
| 222 | 0.0 | 1.0 | 0.0 | 0.0 |
| 224 | 0.0 | 1.0 | 0.0 | 0.0 |
| 225 | 0.0 | 1.0 | 0.0 | 0.0 |
| 227 | 0.0 | 1.0 | 0.0 | 0.0 |
| 228 | 0.0 | 1.0 | 0.0 | 0.0 |
| 230 | 0.0 | 1.0 | 0.0 | 0.0 |
| 231 | 0.0 | 1.0 | 0.0 | 0.0 |
| 234 | 0.0 | 1.0 | 0.0 | 0.0 |
| 237 | 0.0 | 1.0 | 0.0 | 0.0 |
| 238 | 0.0 | 1.0 | 0.0 | 0.0 |
| 241 | 0.0 | 1.0 | 0.0 | 0.0 |
| 242 | 0.0 | 1.0 | 0.0 | 0.0 |
| 243 | 0.0 | 1.0 | 0.0 | 0.0 |
| 245 | 0.0 | 1.0 | 0.0 | 0.0 |
| 250 | 0.0 | 1.0 | 0.0 | 0.0 |
| 254 | 0.0 | 1.0 | 0.0 | 0.0 |
| 257 | 0.0 | 1.0 | 0.0 | 0.0 |
| 261 | 0.0 | 1.0 | 0.0 | 0.0 |
| 262 | 0.0 | 1.0 | 0.0 | 0.0 |
| 263 | 0.0 | 1.0 | 0.0 | 0.0 |
| 265 | 0.0 | 1.0 | 0.0 | 0.0 |
| 266 | 0.0 | 1.0 | 0.0 | 0.0 |
| 275 | 0.0 | 1.0 | 0.0 | 0.0 |
| 276 | 0.0 | 1.0 | 0.0 | 0.0 |
| 279 | 0.0 | 1.0 | 0.0 | 0.0 |
| 284 | 0.0 | 1.0 | 0.0 | 0.0 |
| 190 | 0.0 | 0.5 | 0.5 | 0.0 |
| 280 | 0.0 | 0.5 | 0.5 | 0.0 |ATP7B No.3
#### Chart
| Category | ATP7B N3 | ATP7B N3 | ATP7B N3 | ATP7B N3 |
|---|---|---|---|---|
| 288 | 1.0 | 0.0 | 0.0 | 0.0 |
| 292 | 1.0 | 0.0 | 0.0 | 0.0 |
| 294 | 1.0 | 0.0 | 0.0 | 0.0 |
| 295 | 1.0 | 0.0 | 0.0 | 0.0 |
| 300 | 1.0 | 0.0 | 0.0 | 0.0 |
| 301 | 1.0 | 0.0 | 0.0 | 0.0 |
| 307 | 1.0 | 0.0 | 0.0 | 0.0 |
| 316 | 1.0 | 0.0 | 0.0 | 0.0 |
| 341 | 1.0 | 0.0 | 0.0 | 0.0 |
| 342 | 1.0 | 0.0 | 0.0 | 0.0 |
| 348 | 1.0 | 0.0 | 0.0 | 0.0 |
| 350 | 1.0 | 0.0 | 0.0 | 0.0 |
| 359 | 1.0 | 0.0 | 0.0 | 0.0 |
| 338 | 0.5 | 0.5 | 0.0 | 0.0 |
| 309 | 0.5 | 0.5 | 0.0 | 0.0 |
| 315 | 0.5 | 0.5 | 0.0 | 0.0 |
| 320 | 0.5 | 0.5 | 0.0 | 0.0 |
| 346 | 0.5 | 0.5 | 0.0 | 0.0 |
| 354 | 0.5 | 0.5 | 0.0 | 0.0 |
| 287 | 0.0 | 1.0 | 0.0 | 0.0 |
| 289 | 0.0 | 1.0 | 0.0 | 0.0 |
| 291 | 0.0 | 1.0 | 0.0 | 0.0 |
| 296 | 0.0 | 1.0 | 0.0 | 0.0 |
| 297 | 0.0 | 1.0 | 0.0 | 0.0 |
| 298 | 0.0 | 1.0 | 0.0 | 0.0 |
| 299 | 0.0 | 1.0 | 0.0 | 0.0 |
| 302 | 0.0 | 1.0 | 0.0 | 0.0 |
| 303 | 0.0 | 1.0 | 0.0 | 0.0 |
| 317 | 0.0 | 1.0 | 0.0 | 0.0 |
| 318 | 0.0 | 1.0 | 0.0 | 0.0 |
| 321 | 0.0 | 1.0 | 0.0 | 0.0 |
| 326 | 0.0 | 1.0 | 0.0 | 0.0 |
| 329 | 0.0 | 1.0 | 0.0 | 0.0 |
| 332 | 0.0 | 1.0 | 0.0 | 0.0 |
| 335 | 0.0 | 1.0 | 0.0 | 0.0 |
| 355 | 0.0 | 1.0 | 0.0 | 0.0 |
| 357 | 0.0 | 1.0 | 0.0 | 0.0 |
| 358 | 0.0 | 1.0 | 0.0 | 0.0 |
| 361 | 0.0 | 1.0 | 0.0 | 0.0 |
| 362 | 0.0 | 1.0 | 0.0 | 0.0 |
| 293 | 0.0 | 0.0 | 1.0 | 0.0 |C
D
Total Allelic Frequencies in GRN
#### Chart
| Category | |
|---|---|(%)
No.1
No.2
No.3
WT NHEJ NHEJ+HDR HDR
Total Allelic Frequencies
in ATP7B
(%)
#### Chart
| Category | |
|---|---|No.1
No.2
No.3
WT NHEJ NHEJ+HDR HDR

### Slide 3
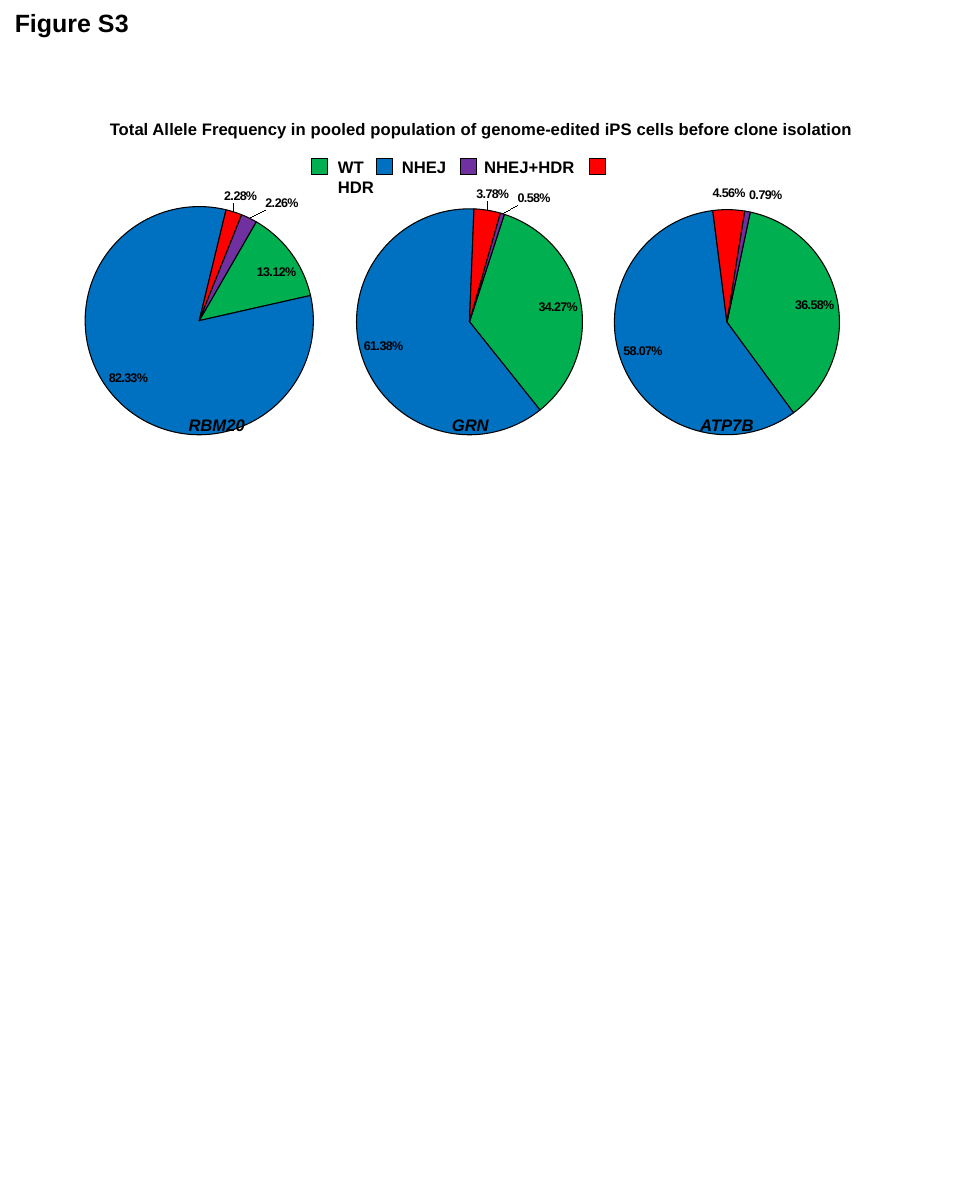

Figure S3
Total Allele Frequency in pooled population of genome-edited iPS cells before clone isolation
WT NHEJ NHEJ+HDR HDR
#### Chart
| Category | |
|---|---|
| WT | 0.13123141210583503 |
| NHEJ | 0.8232995074526338 |
| HDR | 0.02282602142700906 |
| NHEJ+HDR | 0.022643059014522054 |
#### Chart
| Category | |
|---|---|
| WT | 0.3426692081481606 |
| NHEJ | 0.6137509881970814 |
| HDR | 0.03779507783610266 |
| NHEJ+HDR | 0.005784725818655208 |
#### Chart
| Category | |
|---|---|
| WT | 0.3658152040576379 |
| NHEJ | 0.5807241663873334 |
| HDR | 0.045561780985220936 |
| NHEJ+HDR | 0.007898848569807748 |RBM20
GRN
ATP7B

### Slide 4
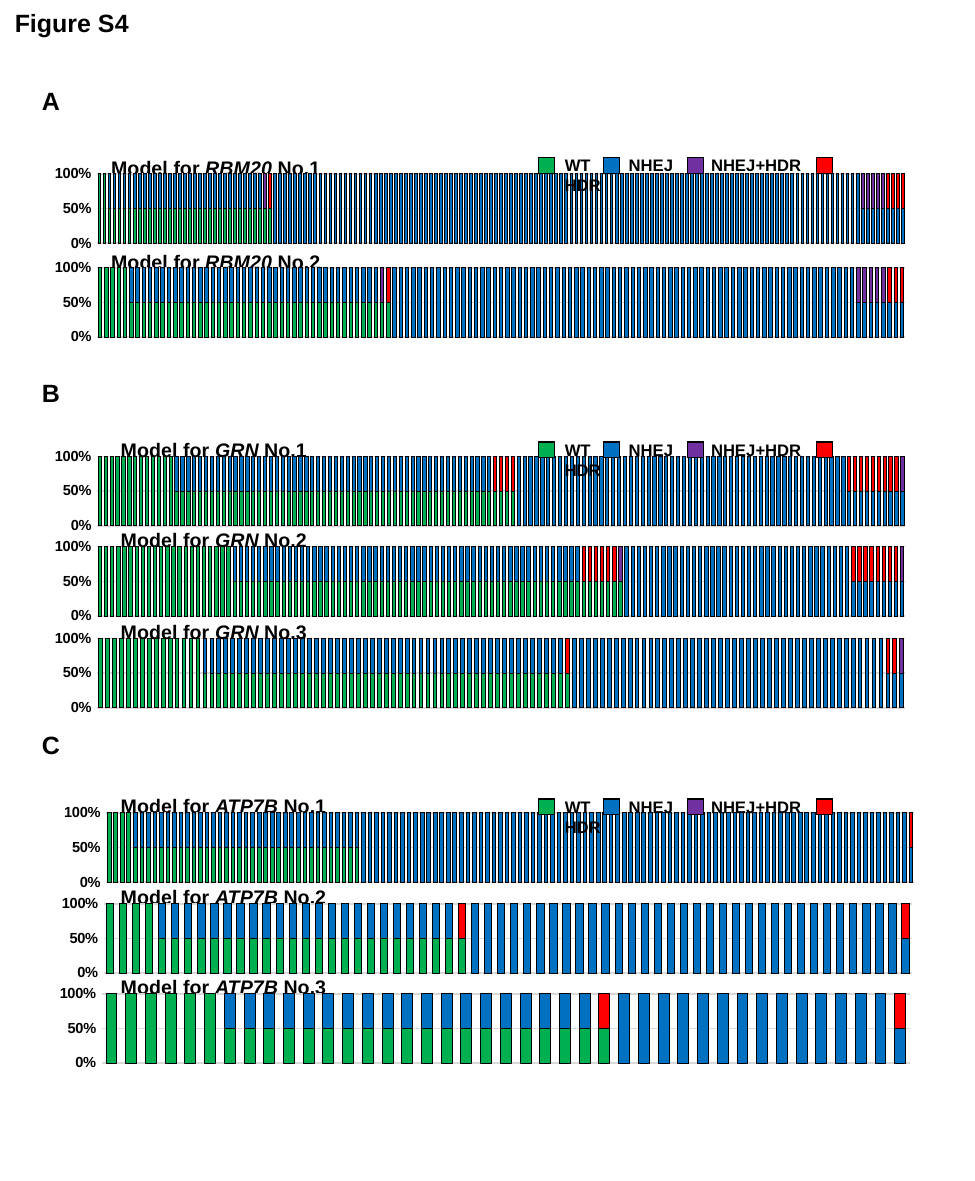

Figure S4
A
WT NHEJ NHEJ+HDR HDR
Model for RBM20 No.1
#### Chart
| Category | WT | NHEJ | HDR | HDR+NHEJ |
|---|---|---|---|---|Model for RBM20 No.2
#### Chart
| Category | WT | NHEJ | HDR | HDR+NHEJ |
|---|---|---|---|---|B
Model for GRN No.1
WT NHEJ NHEJ+HDR HDR
#### Chart
| Category | WT | NHEJ | HDR | HDR+NHEJ |
|---|---|---|---|---|Model for GRN No.2
#### Chart
| Category | WT | NHEJ | HDR | HDR+NHEJ |
|---|---|---|---|---|Model for GRN No.3
#### Chart
| Category | WT | NHEJ | HDR | HDR+NHEJ |
|---|---|---|---|---|C
Model for ATP7B No.1
WT NHEJ NHEJ+HDR HDR
#### Chart
| Category | WT | NHEJ | HDR | HDR+NHEJ |
|---|---|---|---|---|Model for ATP7B No.2
#### Chart
| Category | WT | NHEJ | HDR | HDR+NHEJ |
|---|---|---|---|---|Model for ATP7B No.3
#### Chart
| Category | WT | NHEJ | HDR | HDR+NHEJ |
|---|---|---|---|---|

### Slide 5
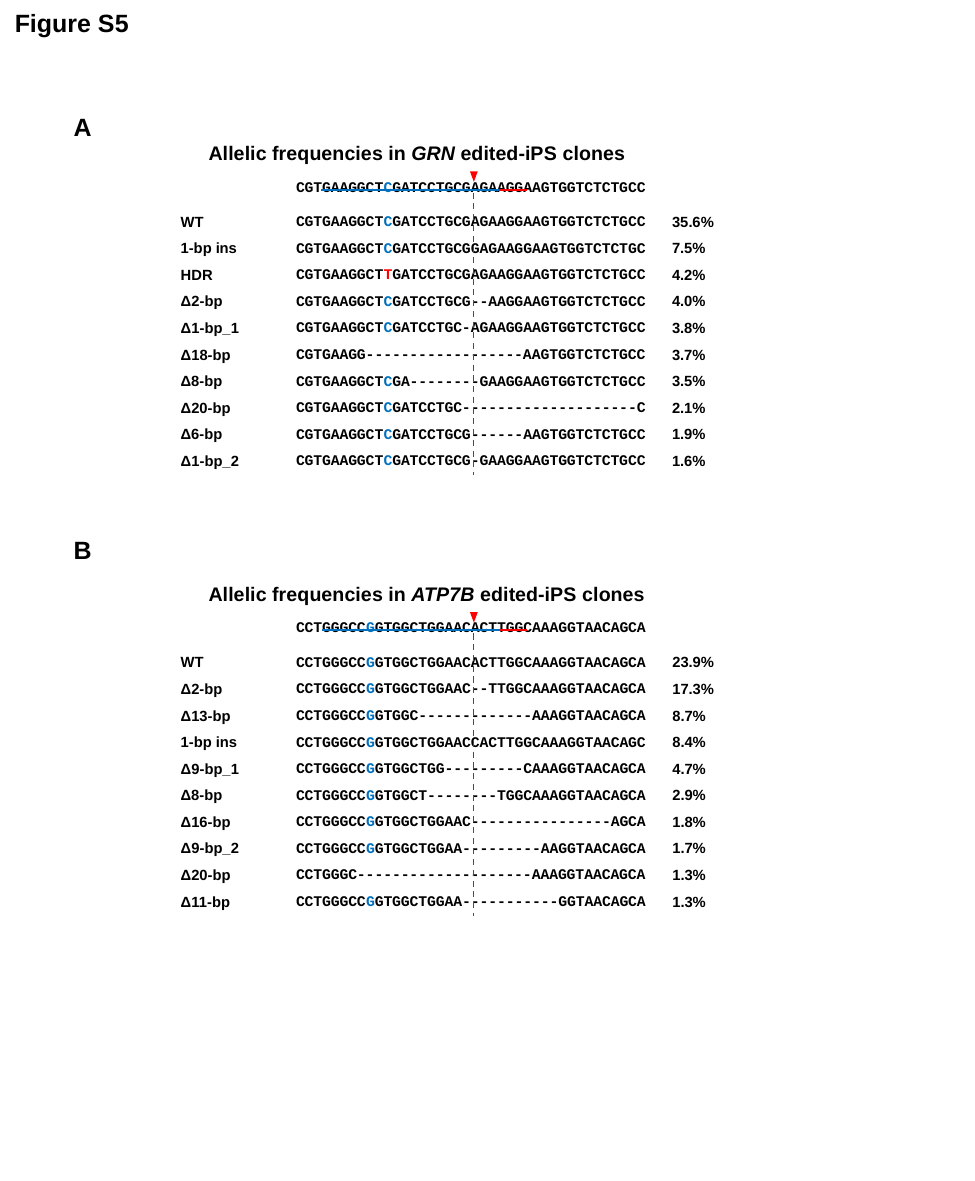

Figure S5
A
Allelic frequencies in GRN edited-iPS clones
CGTGAAGGCTCGATCCTGCGAGAAGGAAGTGGTCTCTGCC
CGTGAAGGCTCGATCCTGCGAGAAGGAAGTGGTCTCTGCC
CGTGAAGGCTCGATCCTGCGGAGAAGGAAGTGGTCTCTGC
CGTGAAGGCTTGATCCTGCGAGAAGGAAGTGGTCTCTGCC
CGTGAAGGCTCGATCCTGCG--AAGGAAGTGGTCTCTGCC
CGTGAAGGCTCGATCCTGC-AGAAGGAAGTGGTCTCTGCC
CGTGAAGG------------------AAGTGGTCTCTGCC
CGTGAAGGCTCGA--------GAAGGAAGTGGTCTCTGCC
CGTGAAGGCTCGATCCTGC--------------------C
CGTGAAGGCTCGATCCTGCG------AAGTGGTCTCTGCC
CGTGAAGGCTCGATCCTGCG-GAAGGAAGTGGTCTCTGCC
WT
1-bp ins
HDR
Δ2-bp
Δ1-bp_1
Δ18-bp
Δ8-bp
Δ20-bp
Δ6-bp
Δ1-bp_2
35.6%
7.5%
4.2%
4.0%
3.8%
3.7%
3.5%
2.1%
1.9%
1.6%
B
Allelic frequencies in ATP7B edited-iPS clones
CCTGGGCCGGTGGCTGGAACACTTGGCAAAGGTAACAGCA
CCTGGGCCGGTGGCTGGAACACTTGGCAAAGGTAACAGCA
CCTGGGCCGGTGGCTGGAAC--TTGGCAAAGGTAACAGCA
CCTGGGCCGGTGGC-------------AAAGGTAACAGCA
CCTGGGCCGGTGGCTGGAACCACTTGGCAAAGGTAACAGC
CCTGGGCCGGTGGCTGG---------CAAAGGTAACAGCA
CCTGGGCCGGTGGCT--------TGGCAAAGGTAACAGCA
CCTGGGCCGGTGGCTGGAAC----------------AGCA
CCTGGGCCGGTGGCTGGAA---------AAGGTAACAGCA
CCTGGGC--------------------AAAGGTAACAGCA
CCTGGGCCGGTGGCTGGAA-----------GGTAACAGCA
WT
Δ2-bp
Δ13-bp
1-bp ins
Δ9-bp_1
Δ8-bp
Δ16-bp
Δ9-bp_2
Δ20-bp
Δ11-bp
23.9%
17.3%
8.7%
8.4%
4.7%
2.9%
1.8%
1.7%
1.3%
1.3%

### Slide 6
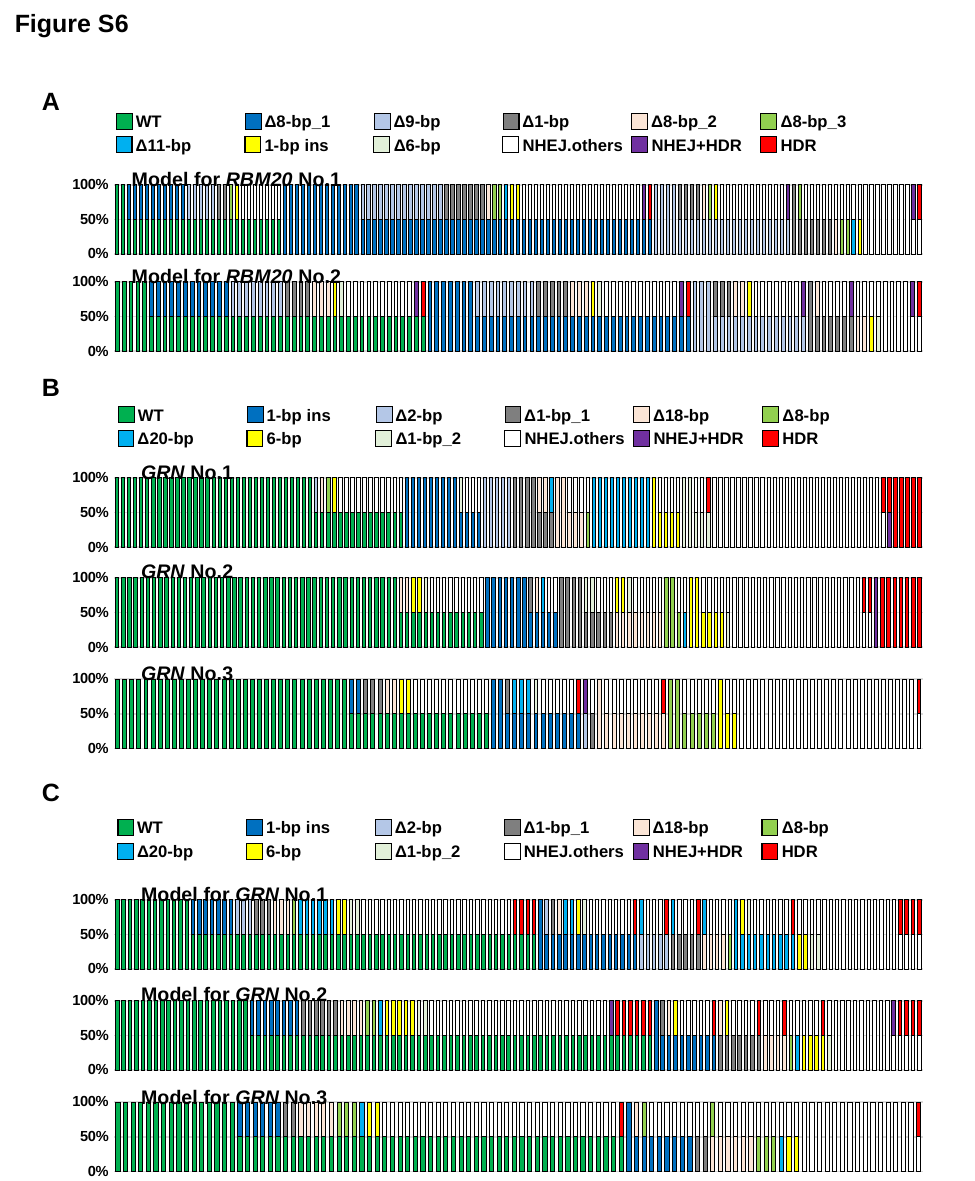

Figure S6
A
WT
Δ8-bp_1
Δ9-bp
Δ1-bp
Δ8-bp_2
Δ8-bp_3
Δ11-bp
1-bp ins
Δ6-bp
NHEJ.others
NHEJ+HDR
HDR
Model for RBM20 No.1
#### Chart
| Category | WT | Δ8-bp_1 | Δ9-bp | Δ1-bp | Δ8-bp_2 | Δ8-bp_3 | Δ11-bp | 1-bp ins | Δ6-bp | NHEJ.others | NHEJ+HDR | HDR |
|---|---|---|---|---|---|---|---|---|---|---|---|---|Model for RBM20 No.2
#### Chart
| Category | WT | Δ8-bp_1 | Δ9-bp | Δ1-bp | Δ8-bp_2 | Δ8-bp_3 | Δ11-bp | 1-bp ins | Δ6-bp | NHEJ.others | NHEJ+HDR | HDR |
|---|---|---|---|---|---|---|---|---|---|---|---|---|B
WT
1-bp ins
Δ2-bp
Δ1-bp_1
Δ18-bp
Δ8-bp
Δ20-bp
6-bp
Δ1-bp_2
NHEJ.others
NHEJ+HDR
HDR
GRN No.1
#### Chart
| Category | WT | 1塩基挿入 | 2塩基欠失 | 1塩基欠失-1 | 18塩基欠失 | 8塩基欠失 | 20塩基欠失 | 6塩基欠失 | 1塩基欠失-2 | NHEJ.others | HDR.NHEJ | HDR |
|---|---|---|---|---|---|---|---|---|---|---|---|---|
| 412 | 100.0 | 0.0 | 0.0 | 0.0 | 0.0 | 0.0 | 0.0 | 0.0 | 0.0 | 0.0 | 0.0 | 0.0 |
| 386 | 100.0 | 0.0 | 0.0 | 0.0 | 0.0 | 0.0 | 0.0 | 0.0 | 0.0 | 0.0 | 0.0 | 0.0 |
| 393 | 100.0 | 0.0 | 0.0 | 0.0 | 0.0 | 0.0 | 0.0 | 0.0 | 0.0 | 0.0 | 0.0 | 0.0 |
| 401 | 100.0 | 0.0 | 0.0 | 0.0 | 0.0 | 0.0 | 0.0 | 0.0 | 0.0 | 0.0 | 0.0 | 0.0 |
| 402 | 100.0 | 0.0 | 0.0 | 0.0 | 0.0 | 0.0 | 0.0 | 0.0 | 0.0 | 0.0 | 0.0 | 0.0 |
| 404 | 100.0 | 0.0 | 0.0 | 0.0 | 0.0 | 0.0 | 0.0 | 0.0 | 0.0 | 0.0 | 0.0 | 0.0 |
| 408 | 100.0 | 0.0 | 0.0 | 0.0 | 0.0 | 0.0 | 0.0 | 0.0 | 0.0 | 0.0 | 0.0 | 0.0 |
| 411 | 100.0 | 0.0 | 0.0 | 0.0 | 0.0 | 0.0 | 0.0 | 0.0 | 0.0 | 0.0 | 0.0 | 0.0 |
| 413 | 100.0 | 0.0 | 0.0 | 0.0 | 0.0 | 0.0 | 0.0 | 0.0 | 0.0 | 0.0 | 0.0 | 0.0 |
| 416 | 100.0 | 0.0 | 0.0 | 0.0 | 0.0 | 0.0 | 0.0 | 0.0 | 0.0 | 0.0 | 0.0 | 0.0 |
| 425 | 100.0 | 0.0 | 0.0 | 0.0 | 0.0 | 0.0 | 0.0 | 0.0 | 0.0 | 0.0 | 0.0 | 0.0 |
| 427 | 100.0 | 0.0 | 0.0 | 0.0 | 0.0 | 0.0 | 0.0 | 0.0 | 0.0 | 0.0 | 0.0 | 0.0 |
| 435 | 100.0 | 0.0 | 0.0 | 0.0 | 0.0 | 0.0 | 0.0 | 0.0 | 0.0 | 0.0 | 0.0 | 0.0 |
| 437 | 100.0 | 0.0 | 0.0 | 0.0 | 0.0 | 0.0 | 0.0 | 0.0 | 0.0 | 0.0 | 0.0 | 0.0 |
| 439 | 100.0 | 0.0 | 0.0 | 0.0 | 0.0 | 0.0 | 0.0 | 0.0 | 0.0 | 0.0 | 0.0 | 0.0 |
| 442 | 100.0 | 0.0 | 0.0 | 0.0 | 0.0 | 0.0 | 0.0 | 0.0 | 0.0 | 0.0 | 0.0 | 0.0 |
| 444 | 100.0 | 0.0 | 0.0 | 0.0 | 0.0 | 0.0 | 0.0 | 0.0 | 0.0 | 0.0 | 0.0 | 0.0 |
| 445 | 100.0 | 0.0 | 0.0 | 0.0 | 0.0 | 0.0 | 0.0 | 0.0 | 0.0 | 0.0 | 0.0 | 0.0 |
| 448 | 100.0 | 0.0 | 0.0 | 0.0 | 0.0 | 0.0 | 0.0 | 0.0 | 0.0 | 0.0 | 0.0 | 0.0 |
| 450 | 100.0 | 0.0 | 0.0 | 0.0 | 0.0 | 0.0 | 0.0 | 0.0 | 0.0 | 0.0 | 0.0 | 0.0 |
| 451 | 100.0 | 0.0 | 0.0 | 0.0 | 0.0 | 0.0 | 0.0 | 0.0 | 0.0 | 0.0 | 0.0 | 0.0 |
| 471 | 100.0 | 0.0 | 0.0 | 0.0 | 0.0 | 0.0 | 0.0 | 0.0 | 0.0 | 0.0 | 0.0 | 0.0 |
| 493 | 100.0 | 0.0 | 0.0 | 0.0 | 0.0 | 0.0 | 0.0 | 0.0 | 0.0 | 0.0 | 0.0 | 0.0 |
| 499 | 100.0 | 0.0 | 0.0 | 0.0 | 0.0 | 0.0 | 0.0 | 0.0 | 0.0 | 0.0 | 0.0 | 0.0 |
| 504 | 100.0 | 0.0 | 0.0 | 0.0 | 0.0 | 0.0 | 0.0 | 0.0 | 0.0 | 0.0 | 0.0 | 0.0 |
| 505 | 100.0 | 0.0 | 0.0 | 0.0 | 0.0 | 0.0 | 0.0 | 0.0 | 0.0 | 0.0 | 0.0 | 0.0 |
| 508 | 100.0 | 0.0 | 0.0 | 0.0 | 0.0 | 0.0 | 0.0 | 0.0 | 0.0 | 0.0 | 0.0 | 0.0 |
| 513 | 100.0 | 0.0 | 0.0 | 0.0 | 0.0 | 0.0 | 0.0 | 0.0 | 0.0 | 0.0 | 0.0 | 0.0 |
| 515 | 100.0 | 0.0 | 0.0 | 0.0 | 0.0 | 0.0 | 0.0 | 0.0 | 0.0 | 0.0 | 0.0 | 0.0 |
| 516 | 100.0 | 0.0 | 0.0 | 0.0 | 0.0 | 0.0 | 0.0 | 0.0 | 0.0 | 0.0 | 0.0 | 0.0 |
| 526 | 100.0 | 0.0 | 0.0 | 0.0 | 0.0 | 0.0 | 0.0 | 0.0 | 0.0 | 0.0 | 0.0 | 0.0 |
| 538 | 100.0 | 0.0 | 0.0 | 0.0 | 0.0 | 0.0 | 0.0 | 0.0 | 0.0 | 0.0 | 0.0 | 0.0 |
| 539 | 100.0 | 0.0 | 0.0 | 0.0 | 0.0 | 0.0 | 0.0 | 0.0 | 0.0 | 0.0 | 0.0 | 0.0 |
| 517 | 50.0 | 0.0 | 50.0 | 0.0 | 0.0 | 0.0 | 0.0 | 0.0 | 0.0 | 0.0 | 0.0 | 0.0 |
| 424 | 50.0 | 0.0 | 0.0 | 0.0 | 50.0 | 0.0 | 0.0 | 0.0 | 0.0 | 0.0 | 0.0 | 0.0 |
| 474 | 50.0 | 0.0 | 0.0 | 0.0 | 0.0 | 50.0 | 0.0 | 0.0 | 0.0 | 0.0 | 0.0 | 0.0 |
| 529 | 50.0 | 0.0 | 0.0 | 0.0 | 0.0 | 0.0 | 0.0 | 50.0 | 0.0 | 0.0 | 0.0 | 0.0 |
| 376 | 50.0 | 0.0 | 0.0 | 0.0 | 0.0 | 0.0 | 0.0 | 0.0 | 0.0 | 50.0 | 0.0 | 0.0 |
| 372 | 50.0 | 0.0 | 0.0 | 0.0 | 0.0 | 0.0 | 0.0 | 0.0 | 0.0 | 50.0 | 0.0 | 0.0 |
| 366 | 50.0 | 0.0 | 0.0 | 0.0 | 0.0 | 0.0 | 0.0 | 0.0 | 0.0 | 50.0 | 0.0 | 0.0 |
| 370 | 50.0 | 0.0 | 0.0 | 0.0 | 0.0 | 0.0 | 0.0 | 0.0 | 0.0 | 50.0 | 0.0 | 0.0 |
| 371 | 50.0 | 0.0 | 0.0 | 0.0 | 0.0 | 0.0 | 0.0 | 0.0 | 0.0 | 50.0 | 0.0 | 0.0 |
| 373 | 50.0 | 0.0 | 0.0 | 0.0 | 0.0 | 0.0 | 0.0 | 0.0 | 0.0 | 50.0 | 0.0 | 0.0 |
| 374 | 50.0 | 0.0 | 0.0 | 0.0 | 0.0 | 0.0 | 0.0 | 0.0 | 0.0 | 50.0 | 0.0 | 0.0 |
| 375 | 50.0 | 0.0 | 0.0 | 0.0 | 0.0 | 0.0 | 0.0 | 0.0 | 0.0 | 50.0 | 0.0 | 0.0 |
| 418 | 50.0 | 0.0 | 0.0 | 0.0 | 0.0 | 0.0 | 0.0 | 0.0 | 0.0 | 50.0 | 0.0 | 0.0 |
| 421 | 50.0 | 0.0 | 0.0 | 0.0 | 0.0 | 0.0 | 0.0 | 0.0 | 0.0 | 50.0 | 0.0 | 0.0 |
| 518 | 50.0 | 0.0 | 0.0 | 0.0 | 0.0 | 0.0 | 0.0 | 0.0 | 0.0 | 50.0 | 0.0 | 0.0 |
| 480 | 0.0 | 100.0 | 0.0 | 0.0 | 0.0 | 0.0 | 0.0 | 0.0 | 0.0 | 0.0 | 0.0 | 0.0 |
| 388 | 0.0 | 100.0 | 0.0 | 0.0 | 0.0 | 0.0 | 0.0 | 0.0 | 0.0 | 0.0 | 0.0 | 0.0 |
| 429 | 0.0 | 100.0 | 0.0 | 0.0 | 0.0 | 0.0 | 0.0 | 0.0 | 0.0 | 0.0 | 0.0 | 0.0 |
| 433 | 0.0 | 100.0 | 0.0 | 0.0 | 0.0 | 0.0 | 0.0 | 0.0 | 0.0 | 0.0 | 0.0 | 0.0 |
| 441 | 0.0 | 100.0 | 0.0 | 0.0 | 0.0 | 0.0 | 0.0 | 0.0 | 0.0 | 0.0 | 0.0 | 0.0 |
| 446 | 0.0 | 100.0 | 0.0 | 0.0 | 0.0 | 0.0 | 0.0 | 0.0 | 0.0 | 0.0 | 0.0 | 0.0 |
| 449 | 0.0 | 100.0 | 0.0 | 0.0 | 0.0 | 0.0 | 0.0 | 0.0 | 0.0 | 0.0 | 0.0 | 0.0 |
| 463 | 0.0 | 100.0 | 0.0 | 0.0 | 0.0 | 0.0 | 0.0 | 0.0 | 0.0 | 0.0 | 0.0 | 0.0 |
| 488 | 0.0 | 100.0 | 0.0 | 0.0 | 0.0 | 0.0 | 0.0 | 0.0 | 0.0 | 0.0 | 0.0 | 0.0 |
| 475 | 0.0 | 50.0 | 0.0 | 0.0 | 0.0 | 0.0 | 0.0 | 0.0 | 50.0 | 0.0 | 0.0 | 0.0 |
| 532 | 0.0 | 50.0 | 0.0 | 0.0 | 0.0 | 0.0 | 0.0 | 0.0 | 0.0 | 50.0 | 0.0 | 0.0 |
| 489 | 0.0 | 50.0 | 0.0 | 0.0 | 0.0 | 0.0 | 0.0 | 0.0 | 0.0 | 50.0 | 0.0 | 0.0 |
| 522 | 0.0 | 50.0 | 0.0 | 0.0 | 0.0 | 0.0 | 0.0 | 0.0 | 0.0 | 50.0 | 0.0 | 0.0 |
| 420 | 0.0 | 0.0 | 100.0 | 0.0 | 0.0 | 0.0 | 0.0 | 0.0 | 0.0 | 0.0 | 0.0 | 0.0 |
| 468 | 0.0 | 0.0 | 100.0 | 0.0 | 0.0 | 0.0 | 0.0 | 0.0 | 0.0 | 0.0 | 0.0 | 0.0 |
| 476 | 0.0 | 0.0 | 100.0 | 0.0 | 0.0 | 0.0 | 0.0 | 0.0 | 0.0 | 0.0 | 0.0 | 0.0 |
| 511 | 0.0 | 0.0 | 100.0 | 0.0 | 0.0 | 0.0 | 0.0 | 0.0 | 0.0 | 0.0 | 0.0 | 0.0 |
| 537 | 0.0 | 0.0 | 100.0 | 0.0 | 0.0 | 0.0 | 0.0 | 0.0 | 0.0 | 0.0 | 0.0 | 0.0 |
| 395 | 0.0 | 0.0 | 0.0 | 100.0 | 0.0 | 0.0 | 0.0 | 0.0 | 0.0 | 0.0 | 0.0 | 0.0 |
| 438 | 0.0 | 0.0 | 0.0 | 100.0 | 0.0 | 0.0 | 0.0 | 0.0 | 0.0 | 0.0 | 0.0 | 0.0 |
| 497 | 0.0 | 0.0 | 0.0 | 100.0 | 0.0 | 0.0 | 0.0 | 0.0 | 0.0 | 0.0 | 0.0 | 0.0 |
| 512 | 0.0 | 0.0 | 0.0 | 100.0 | 0.0 | 0.0 | 0.0 | 0.0 | 0.0 | 0.0 | 0.0 | 0.0 |
| 523 | 0.0 | 0.0 | 0.0 | 50.0 | 50.0 | 0.0 | 0.0 | 0.0 | 0.0 | 0.0 | 0.0 | 0.0 |
| 534 | 0.0 | 0.0 | 0.0 | 50.0 | 50.0 | 0.0 | 0.0 | 0.0 | 0.0 | 0.0 | 0.0 | 0.0 |
| 498 | 0.0 | 0.0 | 0.0 | 50.0 | 0.0 | 0.0 | 50.0 | 0.0 | 0.0 | 0.0 | 0.0 | 0.0 |
| 479 | 0.0 | 0.0 | 0.0 | 0.0 | 100.0 | 0.0 | 0.0 | 0.0 | 0.0 | 0.0 | 0.0 | 0.0 |
| 491 | 0.0 | 0.0 | 0.0 | 0.0 | 100.0 | 0.0 | 0.0 | 0.0 | 0.0 | 0.0 | 0.0 | 0.0 |
| 390 | 0.0 | 0.0 | 0.0 | 0.0 | 50.0 | 0.0 | 0.0 | 0.0 | 0.0 | 50.0 | 0.0 | 0.0 |
| 410 | 0.0 | 0.0 | 0.0 | 0.0 | 50.0 | 0.0 | 0.0 | 0.0 | 0.0 | 50.0 | 0.0 | 0.0 |
| 483 | 0.0 | 0.0 | 0.0 | 0.0 | 50.0 | 0.0 | 0.0 | 0.0 | 0.0 | 50.0 | 0.0 | 0.0 |
| 520 | 0.0 | 0.0 | 0.0 | 0.0 | 0.0 | 50.0 | 0.0 | 0.0 | 0.0 | 50.0 | 0.0 | 0.0 |
| 507 | 0.0 | 0.0 | 0.0 | 0.0 | 0.0 | 0.0 | 100.0 | 0.0 | 0.0 | 0.0 | 0.0 | 0.0 |
| 417 | 0.0 | 0.0 | 0.0 | 0.0 | 0.0 | 0.0 | 100.0 | 0.0 | 0.0 | 0.0 | 0.0 | 0.0 |
| 428 | 0.0 | 0.0 | 0.0 | 0.0 | 0.0 | 0.0 | 100.0 | 0.0 | 0.0 | 0.0 | 0.0 | 0.0 |
| 453 | 0.0 | 0.0 | 0.0 | 0.0 | 0.0 | 0.0 | 100.0 | 0.0 | 0.0 | 0.0 | 0.0 | 0.0 |
| 459 | 0.0 | 0.0 | 0.0 | 0.0 | 0.0 | 0.0 | 100.0 | 0.0 | 0.0 | 0.0 | 0.0 | 0.0 |
| 469 | 0.0 | 0.0 | 0.0 | 0.0 | 0.0 | 0.0 | 100.0 | 0.0 | 0.0 | 0.0 | 0.0 | 0.0 |
| 472 | 0.0 | 0.0 | 0.0 | 0.0 | 0.0 | 0.0 | 100.0 | 0.0 | 0.0 | 0.0 | 0.0 | 0.0 |
| 509 | 0.0 | 0.0 | 0.0 | 0.0 | 0.0 | 0.0 | 100.0 | 0.0 | 0.0 | 0.0 | 0.0 | 0.0 |
| 530 | 0.0 | 0.0 | 0.0 | 0.0 | 0.0 | 0.0 | 100.0 | 0.0 | 0.0 | 0.0 | 0.0 | 0.0 |
| 531 | 0.0 | 0.0 | 0.0 | 0.0 | 0.0 | 0.0 | 100.0 | 0.0 | 0.0 | 0.0 | 0.0 | 0.0 |
| 430 | 0.0 | 0.0 | 0.0 | 0.0 | 0.0 | 0.0 | 0.0 | 100.0 | 0.0 | 0.0 | 0.0 | 0.0 |
| 403 | 0.0 | 0.0 | 0.0 | 0.0 | 0.0 | 0.0 | 0.0 | 50.0 | 0.0 | 50.0 | 0.0 | 0.0 |
| 426 | 0.0 | 0.0 | 0.0 | 0.0 | 0.0 | 0.0 | 0.0 | 50.0 | 0.0 | 50.0 | 0.0 | 0.0 |
| 462 | 0.0 | 0.0 | 0.0 | 0.0 | 0.0 | 0.0 | 0.0 | 50.0 | 0.0 | 50.0 | 0.0 | 0.0 |
| 466 | 0.0 | 0.0 | 0.0 | 0.0 | 0.0 | 0.0 | 0.0 | 50.0 | 0.0 | 50.0 | 0.0 | 0.0 |
| 461 | 0.0 | 0.0 | 0.0 | 0.0 | 0.0 | 0.0 | 0.0 | 0.0 | 100.0 | 0.0 | 0.0 | 0.0 |
| 506 | 0.0 | 0.0 | 0.0 | 0.0 | 0.0 | 0.0 | 0.0 | 0.0 | 100.0 | 0.0 | 0.0 | 0.0 |
| 432 | 0.0 | 0.0 | 0.0 | 0.0 | 0.0 | 0.0 | 0.0 | 0.0 | 50.0 | 50.0 | 0.0 | 0.0 |
| 440 | 0.0 | 0.0 | 0.0 | 0.0 | 0.0 | 0.0 | 0.0 | 0.0 | 50.0 | 50.0 | 0.0 | 0.0 |
| 385 | 0.0 | 0.0 | 0.0 | 0.0 | 0.0 | 0.0 | 0.0 | 0.0 | 50.0 | 0.0 | 0.0 | 50.0 |
| 419 | 0.0 | 0.0 | 0.0 | 0.0 | 0.0 | 0.0 | 0.0 | 0.0 | 0.0 | 100.0 | 0.0 | 0.0 |
| 454 | 0.0 | 0.0 | 0.0 | 0.0 | 0.0 | 0.0 | 0.0 | 0.0 | 0.0 | 100.0 | 0.0 | 0.0 |
| 387 | 0.0 | 0.0 | 0.0 | 0.0 | 0.0 | 0.0 | 0.0 | 0.0 | 0.0 | 100.0 | 0.0 | 0.0 |
| 389 | 0.0 | 0.0 | 0.0 | 0.0 | 0.0 | 0.0 | 0.0 | 0.0 | 0.0 | 100.0 | 0.0 | 0.0 |
| 392 | 0.0 | 0.0 | 0.0 | 0.0 | 0.0 | 0.0 | 0.0 | 0.0 | 0.0 | 100.0 | 0.0 | 0.0 |
| 394 | 0.0 | 0.0 | 0.0 | 0.0 | 0.0 | 0.0 | 0.0 | 0.0 | 0.0 | 100.0 | 0.0 | 0.0 |
| 397 | 0.0 | 0.0 | 0.0 | 0.0 | 0.0 | 0.0 | 0.0 | 0.0 | 0.0 | 100.0 | 0.0 | 0.0 |
| 405 | 0.0 | 0.0 | 0.0 | 0.0 | 0.0 | 0.0 | 0.0 | 0.0 | 0.0 | 100.0 | 0.0 | 0.0 |
| 406 | 0.0 | 0.0 | 0.0 | 0.0 | 0.0 | 0.0 | 0.0 | 0.0 | 0.0 | 100.0 | 0.0 | 0.0 |
| 434 | 0.0 | 0.0 | 0.0 | 0.0 | 0.0 | 0.0 | 0.0 | 0.0 | 0.0 | 100.0 | 0.0 | 0.0 |
| 456 | 0.0 | 0.0 | 0.0 | 0.0 | 0.0 | 0.0 | 0.0 | 0.0 | 0.0 | 100.0 | 0.0 | 0.0 |
| 457 | 0.0 | 0.0 | 0.0 | 0.0 | 0.0 | 0.0 | 0.0 | 0.0 | 0.0 | 100.0 | 0.0 | 0.0 |
| 458 | 0.0 | 0.0 | 0.0 | 0.0 | 0.0 | 0.0 | 0.0 | 0.0 | 0.0 | 100.0 | 0.0 | 0.0 |
| 470 | 0.0 | 0.0 | 0.0 | 0.0 | 0.0 | 0.0 | 0.0 | 0.0 | 0.0 | 100.0 | 0.0 | 0.0 |
| 473 | 0.0 | 0.0 | 0.0 | 0.0 | 0.0 | 0.0 | 0.0 | 0.0 | 0.0 | 100.0 | 0.0 | 0.0 |
| 477 | 0.0 | 0.0 | 0.0 | 0.0 | 0.0 | 0.0 | 0.0 | 0.0 | 0.0 | 100.0 | 0.0 | 0.0 |
| 481 | 0.0 | 0.0 | 0.0 | 0.0 | 0.0 | 0.0 | 0.0 | 0.0 | 0.0 | 100.0 | 0.0 | 0.0 |
| 482 | 0.0 | 0.0 | 0.0 | 0.0 | 0.0 | 0.0 | 0.0 | 0.0 | 0.0 | 100.0 | 0.0 | 0.0 |
| 485 | 0.0 | 0.0 | 0.0 | 0.0 | 0.0 | 0.0 | 0.0 | 0.0 | 0.0 | 100.0 | 0.0 | 0.0 |
| 486 | 0.0 | 0.0 | 0.0 | 0.0 | 0.0 | 0.0 | 0.0 | 0.0 | 0.0 | 100.0 | 0.0 | 0.0 |
| 490 | 0.0 | 0.0 | 0.0 | 0.0 | 0.0 | 0.0 | 0.0 | 0.0 | 0.0 | 100.0 | 0.0 | 0.0 |
| 500 | 0.0 | 0.0 | 0.0 | 0.0 | 0.0 | 0.0 | 0.0 | 0.0 | 0.0 | 100.0 | 0.0 | 0.0 |
| 501 | 0.0 | 0.0 | 0.0 | 0.0 | 0.0 | 0.0 | 0.0 | 0.0 | 0.0 | 100.0 | 0.0 | 0.0 |
| 502 | 0.0 | 0.0 | 0.0 | 0.0 | 0.0 | 0.0 | 0.0 | 0.0 | 0.0 | 100.0 | 0.0 | 0.0 |
| 524 | 0.0 | 0.0 | 0.0 | 0.0 | 0.0 | 0.0 | 0.0 | 0.0 | 0.0 | 100.0 | 0.0 | 0.0 |
| 527 | 0.0 | 0.0 | 0.0 | 0.0 | 0.0 | 0.0 | 0.0 | 0.0 | 0.0 | 100.0 | 0.0 | 0.0 |
| 533 | 0.0 | 0.0 | 0.0 | 0.0 | 0.0 | 0.0 | 0.0 | 0.0 | 0.0 | 100.0 | 0.0 | 0.0 |
| 536 | 0.0 | 0.0 | 0.0 | 0.0 | 0.0 | 0.0 | 0.0 | 0.0 | 0.0 | 100.0 | 0.0 | 0.0 |
| 528 | 0.0 | 0.0 | 0.0 | 0.0 | 0.0 | 0.0 | 0.0 | 0.0 | 0.0 | 50.0 | 0.0 | 50.0 |
| 447 | 0.0 | 0.0 | 0.0 | 0.0 | 0.0 | 0.0 | 0.0 | 0.0 | 0.0 | 0.0 | 50.0 | 50.0 |
| 409 | 0.0 | 0.0 | 0.0 | 0.0 | 0.0 | 0.0 | 0.0 | 0.0 | 0.0 | 0.0 | 0.0 | 100.0 |
| 452 | 0.0 | 0.0 | 0.0 | 0.0 | 0.0 | 0.0 | 0.0 | 0.0 | 0.0 | 0.0 | 0.0 | 100.0 |
| 455 | 0.0 | 0.0 | 0.0 | 0.0 | 0.0 | 0.0 | 0.0 | 0.0 | 0.0 | 0.0 | 0.0 | 100.0 |
| 467 | 0.0 | 0.0 | 0.0 | 0.0 | 0.0 | 0.0 | 0.0 | 0.0 | 0.0 | 0.0 | 0.0 | 100.0 |
| 510 | 0.0 | 0.0 | 0.0 | 0.0 | 0.0 | 0.0 | 0.0 | 0.0 | 0.0 | 0.0 | 0.0 | 100.0 |GRN No.2
#### Chart
| Category | WT | 1塩基挿入 | 2塩基欠失 | 1塩基欠失-1 | 18塩基欠失 | 8塩基欠失 | 20塩基欠失 | 6塩基欠失 | 1塩基欠失-2 | NHEJ.others | HDR.NHEJ | HDR |
|---|---|---|---|---|---|---|---|---|---|---|---|---|
| 544 | 100.0 | 0.0 | 0.0 | 0.0 | 0.0 | 0.0 | 0.0 | 0.0 | 0.0 | 0.0 | 0.0 | 0.0 |
| 549 | 100.0 | 0.0 | 0.0 | 0.0 | 0.0 | 0.0 | 0.0 | 0.0 | 0.0 | 0.0 | 0.0 | 0.0 |
| 550 | 100.0 | 0.0 | 0.0 | 0.0 | 0.0 | 0.0 | 0.0 | 0.0 | 0.0 | 0.0 | 0.0 | 0.0 |
| 551 | 100.0 | 0.0 | 0.0 | 0.0 | 0.0 | 0.0 | 0.0 | 0.0 | 0.0 | 0.0 | 0.0 | 0.0 |
| 552 | 100.0 | 0.0 | 0.0 | 0.0 | 0.0 | 0.0 | 0.0 | 0.0 | 0.0 | 0.0 | 0.0 | 0.0 |
| 554 | 100.0 | 0.0 | 0.0 | 0.0 | 0.0 | 0.0 | 0.0 | 0.0 | 0.0 | 0.0 | 0.0 | 0.0 |
| 556 | 100.0 | 0.0 | 0.0 | 0.0 | 0.0 | 0.0 | 0.0 | 0.0 | 0.0 | 0.0 | 0.0 | 0.0 |
| 559 | 100.0 | 0.0 | 0.0 | 0.0 | 0.0 | 0.0 | 0.0 | 0.0 | 0.0 | 0.0 | 0.0 | 0.0 |
| 560 | 100.0 | 0.0 | 0.0 | 0.0 | 0.0 | 0.0 | 0.0 | 0.0 | 0.0 | 0.0 | 0.0 | 0.0 |
| 562 | 100.0 | 0.0 | 0.0 | 0.0 | 0.0 | 0.0 | 0.0 | 0.0 | 0.0 | 0.0 | 0.0 | 0.0 |
| 563 | 100.0 | 0.0 | 0.0 | 0.0 | 0.0 | 0.0 | 0.0 | 0.0 | 0.0 | 0.0 | 0.0 | 0.0 |
| 569 | 100.0 | 0.0 | 0.0 | 0.0 | 0.0 | 0.0 | 0.0 | 0.0 | 0.0 | 0.0 | 0.0 | 0.0 |
| 574 | 100.0 | 0.0 | 0.0 | 0.0 | 0.0 | 0.0 | 0.0 | 0.0 | 0.0 | 0.0 | 0.0 | 0.0 |
| 575 | 100.0 | 0.0 | 0.0 | 0.0 | 0.0 | 0.0 | 0.0 | 0.0 | 0.0 | 0.0 | 0.0 | 0.0 |
| 581 | 100.0 | 0.0 | 0.0 | 0.0 | 0.0 | 0.0 | 0.0 | 0.0 | 0.0 | 0.0 | 0.0 | 0.0 |
| 582 | 100.0 | 0.0 | 0.0 | 0.0 | 0.0 | 0.0 | 0.0 | 0.0 | 0.0 | 0.0 | 0.0 | 0.0 |
| 587 | 100.0 | 0.0 | 0.0 | 0.0 | 0.0 | 0.0 | 0.0 | 0.0 | 0.0 | 0.0 | 0.0 | 0.0 |
| 595 | 100.0 | 0.0 | 0.0 | 0.0 | 0.0 | 0.0 | 0.0 | 0.0 | 0.0 | 0.0 | 0.0 | 0.0 |
| 596 | 100.0 | 0.0 | 0.0 | 0.0 | 0.0 | 0.0 | 0.0 | 0.0 | 0.0 | 0.0 | 0.0 | 0.0 |
| 597 | 100.0 | 0.0 | 0.0 | 0.0 | 0.0 | 0.0 | 0.0 | 0.0 | 0.0 | 0.0 | 0.0 | 0.0 |
| 602 | 100.0 | 0.0 | 0.0 | 0.0 | 0.0 | 0.0 | 0.0 | 0.0 | 0.0 | 0.0 | 0.0 | 0.0 |
| 603 | 100.0 | 0.0 | 0.0 | 0.0 | 0.0 | 0.0 | 0.0 | 0.0 | 0.0 | 0.0 | 0.0 | 0.0 |
| 607 | 100.0 | 0.0 | 0.0 | 0.0 | 0.0 | 0.0 | 0.0 | 0.0 | 0.0 | 0.0 | 0.0 | 0.0 |
| 609 | 100.0 | 0.0 | 0.0 | 0.0 | 0.0 | 0.0 | 0.0 | 0.0 | 0.0 | 0.0 | 0.0 | 0.0 |
| 611 | 100.0 | 0.0 | 0.0 | 0.0 | 0.0 | 0.0 | 0.0 | 0.0 | 0.0 | 0.0 | 0.0 | 0.0 |
| 614 | 100.0 | 0.0 | 0.0 | 0.0 | 0.0 | 0.0 | 0.0 | 0.0 | 0.0 | 0.0 | 0.0 | 0.0 |
| 617 | 100.0 | 0.0 | 0.0 | 0.0 | 0.0 | 0.0 | 0.0 | 0.0 | 0.0 | 0.0 | 0.0 | 0.0 |
| 618 | 100.0 | 0.0 | 0.0 | 0.0 | 0.0 | 0.0 | 0.0 | 0.0 | 0.0 | 0.0 | 0.0 | 0.0 |
| 619 | 100.0 | 0.0 | 0.0 | 0.0 | 0.0 | 0.0 | 0.0 | 0.0 | 0.0 | 0.0 | 0.0 | 0.0 |
| 625 | 100.0 | 0.0 | 0.0 | 0.0 | 0.0 | 0.0 | 0.0 | 0.0 | 0.0 | 0.0 | 0.0 | 0.0 |
| 627 | 100.0 | 0.0 | 0.0 | 0.0 | 0.0 | 0.0 | 0.0 | 0.0 | 0.0 | 0.0 | 0.0 | 0.0 |
| 634 | 100.0 | 0.0 | 0.0 | 0.0 | 0.0 | 0.0 | 0.0 | 0.0 | 0.0 | 0.0 | 0.0 | 0.0 |
| 636 | 100.0 | 0.0 | 0.0 | 0.0 | 0.0 | 0.0 | 0.0 | 0.0 | 0.0 | 0.0 | 0.0 | 0.0 |
| 640 | 100.0 | 0.0 | 0.0 | 0.0 | 0.0 | 0.0 | 0.0 | 0.0 | 0.0 | 0.0 | 0.0 | 0.0 |
| 641 | 100.0 | 0.0 | 0.0 | 0.0 | 0.0 | 0.0 | 0.0 | 0.0 | 0.0 | 0.0 | 0.0 | 0.0 |
| 643 | 100.0 | 0.0 | 0.0 | 0.0 | 0.0 | 0.0 | 0.0 | 0.0 | 0.0 | 0.0 | 0.0 | 0.0 |
| 647 | 100.0 | 0.0 | 0.0 | 0.0 | 0.0 | 0.0 | 0.0 | 0.0 | 0.0 | 0.0 | 0.0 | 0.0 |
| 648 | 100.0 | 0.0 | 0.0 | 0.0 | 0.0 | 0.0 | 0.0 | 0.0 | 0.0 | 0.0 | 0.0 | 0.0 |
| 655 | 100.0 | 0.0 | 0.0 | 0.0 | 0.0 | 0.0 | 0.0 | 0.0 | 0.0 | 0.0 | 0.0 | 0.0 |
| 659 | 100.0 | 0.0 | 0.0 | 0.0 | 0.0 | 0.0 | 0.0 | 0.0 | 0.0 | 0.0 | 0.0 | 0.0 |
| 660 | 100.0 | 0.0 | 0.0 | 0.0 | 0.0 | 0.0 | 0.0 | 0.0 | 0.0 | 0.0 | 0.0 | 0.0 |
| 674 | 100.0 | 0.0 | 0.0 | 0.0 | 0.0 | 0.0 | 0.0 | 0.0 | 0.0 | 0.0 | 0.0 | 0.0 |
| 688 | 100.0 | 0.0 | 0.0 | 0.0 | 0.0 | 0.0 | 0.0 | 0.0 | 0.0 | 0.0 | 0.0 | 0.0 |
| 691 | 100.0 | 0.0 | 0.0 | 0.0 | 0.0 | 0.0 | 0.0 | 0.0 | 0.0 | 0.0 | 0.0 | 0.0 |
| 692 | 100.0 | 0.0 | 0.0 | 0.0 | 0.0 | 0.0 | 0.0 | 0.0 | 0.0 | 0.0 | 0.0 | 0.0 |
| 696 | 100.0 | 0.0 | 0.0 | 0.0 | 0.0 | 0.0 | 0.0 | 0.0 | 0.0 | 0.0 | 0.0 | 0.0 |
| 644 | 50.0 | 0.0 | 0.0 | 0.0 | 50.0 | 0.0 | 0.0 | 0.0 | 0.0 | 0.0 | 0.0 | 0.0 |
| 664 | 50.0 | 0.0 | 0.0 | 0.0 | 50.0 | 0.0 | 0.0 | 0.0 | 0.0 | 0.0 | 0.0 | 0.0 |
| 580 | 50.0 | 0.0 | 0.0 | 0.0 | 0.0 | 0.0 | 0.0 | 50.0 | 0.0 | 0.0 | 0.0 | 0.0 |
| 646 | 50.0 | 0.0 | 0.0 | 0.0 | 0.0 | 0.0 | 0.0 | 50.0 | 0.0 | 0.0 | 0.0 | 0.0 |
| 632 | 50.0 | 0.0 | 0.0 | 0.0 | 0.0 | 0.0 | 0.0 | 0.0 | 50.0 | 0.0 | 0.0 | 0.0 |
| 545 | 50.0 | 0.0 | 0.0 | 0.0 | 0.0 | 0.0 | 0.0 | 0.0 | 0.0 | 50.0 | 0.0 | 0.0 |
| 561 | 50.0 | 0.0 | 0.0 | 0.0 | 0.0 | 0.0 | 0.0 | 0.0 | 0.0 | 50.0 | 0.0 | 0.0 |
| 576 | 50.0 | 0.0 | 0.0 | 0.0 | 0.0 | 0.0 | 0.0 | 0.0 | 0.0 | 50.0 | 0.0 | 0.0 |
| 579 | 50.0 | 0.0 | 0.0 | 0.0 | 0.0 | 0.0 | 0.0 | 0.0 | 0.0 | 50.0 | 0.0 | 0.0 |
| 604 | 50.0 | 0.0 | 0.0 | 0.0 | 0.0 | 0.0 | 0.0 | 0.0 | 0.0 | 50.0 | 0.0 | 0.0 |
| 652 | 50.0 | 0.0 | 0.0 | 0.0 | 0.0 | 0.0 | 0.0 | 0.0 | 0.0 | 50.0 | 0.0 | 0.0 |
| 678 | 50.0 | 0.0 | 0.0 | 0.0 | 0.0 | 0.0 | 0.0 | 0.0 | 0.0 | 50.0 | 0.0 | 0.0 |
| 693 | 50.0 | 0.0 | 0.0 | 0.0 | 0.0 | 0.0 | 0.0 | 0.0 | 0.0 | 50.0 | 0.0 | 0.0 |
| 694 | 50.0 | 0.0 | 0.0 | 0.0 | 0.0 | 0.0 | 0.0 | 0.0 | 0.0 | 50.0 | 0.0 | 0.0 |
| 568 | 0.0 | 100.0 | 0.0 | 0.0 | 0.0 | 0.0 | 0.0 | 0.0 | 0.0 | 0.0 | 0.0 | 0.0 |
| 594 | 0.0 | 100.0 | 0.0 | 0.0 | 0.0 | 0.0 | 0.0 | 0.0 | 0.0 | 0.0 | 0.0 | 0.0 |
| 621 | 0.0 | 100.0 | 0.0 | 0.0 | 0.0 | 0.0 | 0.0 | 0.0 | 0.0 | 0.0 | 0.0 | 0.0 |
| 633 | 0.0 | 100.0 | 0.0 | 0.0 | 0.0 | 0.0 | 0.0 | 0.0 | 0.0 | 0.0 | 0.0 | 0.0 |
| 673 | 0.0 | 100.0 | 0.0 | 0.0 | 0.0 | 0.0 | 0.0 | 0.0 | 0.0 | 0.0 | 0.0 | 0.0 |
| 676 | 0.0 | 100.0 | 0.0 | 0.0 | 0.0 | 0.0 | 0.0 | 0.0 | 0.0 | 0.0 | 0.0 | 0.0 |
| 698 | 0.0 | 100.0 | 0.0 | 0.0 | 0.0 | 0.0 | 0.0 | 0.0 | 0.0 | 0.0 | 0.0 | 0.0 |
| 689 | 0.0 | 50.0 | 0.0 | 50.0 | 0.0 | 0.0 | 0.0 | 0.0 | 0.0 | 0.0 | 0.0 | 0.0 |
| 684 | 0.0 | 50.0 | 0.0 | 0.0 | 50.0 | 0.0 | 0.0 | 0.0 | 0.0 | 0.0 | 0.0 | 0.0 |
| 666 | 0.0 | 50.0 | 0.0 | 0.0 | 0.0 | 0.0 | 50.0 | 0.0 | 0.0 | 0.0 | 0.0 | 0.0 |
| 577 | 0.0 | 50.0 | 0.0 | 0.0 | 0.0 | 0.0 | 0.0 | 0.0 | 0.0 | 50.0 | 0.0 | 0.0 |
| 623 | 0.0 | 50.0 | 0.0 | 0.0 | 0.0 | 0.0 | 0.0 | 0.0 | 0.0 | 50.0 | 0.0 | 0.0 |
| 553 | 0.0 | 0.0 | 0.0 | 100.0 | 0.0 | 0.0 | 0.0 | 0.0 | 0.0 | 0.0 | 0.0 | 0.0 |
| 608 | 0.0 | 0.0 | 0.0 | 100.0 | 0.0 | 0.0 | 0.0 | 0.0 | 0.0 | 0.0 | 0.0 | 0.0 |
| 649 | 0.0 | 0.0 | 0.0 | 100.0 | 0.0 | 0.0 | 0.0 | 0.0 | 0.0 | 0.0 | 0.0 | 0.0 |
| 685 | 0.0 | 0.0 | 0.0 | 100.0 | 0.0 | 0.0 | 0.0 | 0.0 | 0.0 | 0.0 | 0.0 | 0.0 |
| 542 | 0.0 | 0.0 | 0.0 | 50.0 | 0.0 | 0.0 | 0.0 | 0.0 | 50.0 | 0.0 | 0.0 | 0.0 |
| 558 | 0.0 | 0.0 | 0.0 | 50.0 | 0.0 | 0.0 | 0.0 | 0.0 | 50.0 | 0.0 | 0.0 | 0.0 |
| 565 | 0.0 | 0.0 | 0.0 | 50.0 | 0.0 | 0.0 | 0.0 | 0.0 | 0.0 | 50.0 | 0.0 | 0.0 |
| 598 | 0.0 | 0.0 | 0.0 | 50.0 | 0.0 | 0.0 | 0.0 | 0.0 | 0.0 | 50.0 | 0.0 | 0.0 |
| 651 | 0.0 | 0.0 | 0.0 | 50.0 | 0.0 | 0.0 | 0.0 | 0.0 | 0.0 | 50.0 | 0.0 | 0.0 |
| 557 | 0.0 | 0.0 | 0.0 | 0.0 | 50.0 | 0.0 | 0.0 | 50.0 | 0.0 | 0.0 | 0.0 | 0.0 |
| 690 | 0.0 | 0.0 | 0.0 | 0.0 | 50.0 | 0.0 | 0.0 | 50.0 | 0.0 | 0.0 | 0.0 | 0.0 |
| 658 | 0.0 | 0.0 | 0.0 | 0.0 | 50.0 | 0.0 | 0.0 | 0.0 | 50.0 | 0.0 | 0.0 | 0.0 |
| 624 | 0.0 | 0.0 | 0.0 | 0.0 | 50.0 | 0.0 | 0.0 | 0.0 | 0.0 | 50.0 | 0.0 | 0.0 |
| 654 | 0.0 | 0.0 | 0.0 | 0.0 | 50.0 | 0.0 | 0.0 | 0.0 | 0.0 | 50.0 | 0.0 | 0.0 |
| 669 | 0.0 | 0.0 | 0.0 | 0.0 | 50.0 | 0.0 | 0.0 | 0.0 | 0.0 | 50.0 | 0.0 | 0.0 |
| 672 | 0.0 | 0.0 | 0.0 | 0.0 | 50.0 | 0.0 | 0.0 | 0.0 | 0.0 | 50.0 | 0.0 | 0.0 |
| 677 | 0.0 | 0.0 | 0.0 | 0.0 | 50.0 | 0.0 | 0.0 | 0.0 | 0.0 | 50.0 | 0.0 | 0.0 |
| 555 | 0.0 | 0.0 | 0.0 | 0.0 | 0.0 | 100.0 | 0.0 | 0.0 | 0.0 | 0.0 | 0.0 | 0.0 |
| 653 | 0.0 | 0.0 | 0.0 | 0.0 | 0.0 | 100.0 | 0.0 | 0.0 | 0.0 | 0.0 | 0.0 | 0.0 |
| 543 | 0.0 | 0.0 | 0.0 | 0.0 | 0.0 | 50.0 | 0.0 | 0.0 | 0.0 | 50.0 | 0.0 | 0.0 |
| 683 | 0.0 | 0.0 | 0.0 | 0.0 | 0.0 | 0.0 | 50.0 | 0.0 | 0.0 | 50.0 | 0.0 | 0.0 |
| 616 | 0.0 | 0.0 | 0.0 | 0.0 | 0.0 | 0.0 | 0.0 | 100.0 | 0.0 | 0.0 | 0.0 | 0.0 |
| 670 | 0.0 | 0.0 | 0.0 | 0.0 | 0.0 | 0.0 | 0.0 | 100.0 | 0.0 | 0.0 | 0.0 | 0.0 |
| 571 | 0.0 | 0.0 | 0.0 | 0.0 | 0.0 | 0.0 | 0.0 | 50.0 | 0.0 | 50.0 | 0.0 | 0.0 |
| 593 | 0.0 | 0.0 | 0.0 | 0.0 | 0.0 | 0.0 | 0.0 | 50.0 | 0.0 | 50.0 | 0.0 | 0.0 |
| 622 | 0.0 | 0.0 | 0.0 | 0.0 | 0.0 | 0.0 | 0.0 | 50.0 | 0.0 | 50.0 | 0.0 | 0.0 |
| 635 | 0.0 | 0.0 | 0.0 | 0.0 | 0.0 | 0.0 | 0.0 | 50.0 | 0.0 | 50.0 | 0.0 | 0.0 |
| 629 | 0.0 | 0.0 | 0.0 | 0.0 | 0.0 | 0.0 | 0.0 | 0.0 | 50.0 | 50.0 | 0.0 | 0.0 |
| 540 | 0.0 | 0.0 | 0.0 | 0.0 | 0.0 | 0.0 | 0.0 | 0.0 | 0.0 | 100.0 | 0.0 | 0.0 |
| 541 | 0.0 | 0.0 | 0.0 | 0.0 | 0.0 | 0.0 | 0.0 | 0.0 | 0.0 | 100.0 | 0.0 | 0.0 |
| 546 | 0.0 | 0.0 | 0.0 | 0.0 | 0.0 | 0.0 | 0.0 | 0.0 | 0.0 | 100.0 | 0.0 | 0.0 |
| 566 | 0.0 | 0.0 | 0.0 | 0.0 | 0.0 | 0.0 | 0.0 | 0.0 | 0.0 | 100.0 | 0.0 | 0.0 |
| 567 | 0.0 | 0.0 | 0.0 | 0.0 | 0.0 | 0.0 | 0.0 | 0.0 | 0.0 | 100.0 | 0.0 | 0.0 |
| 570 | 0.0 | 0.0 | 0.0 | 0.0 | 0.0 | 0.0 | 0.0 | 0.0 | 0.0 | 100.0 | 0.0 | 0.0 |
| 572 | 0.0 | 0.0 | 0.0 | 0.0 | 0.0 | 0.0 | 0.0 | 0.0 | 0.0 | 100.0 | 0.0 | 0.0 |
| 578 | 0.0 | 0.0 | 0.0 | 0.0 | 0.0 | 0.0 | 0.0 | 0.0 | 0.0 | 100.0 | 0.0 | 0.0 |
| 586 | 0.0 | 0.0 | 0.0 | 0.0 | 0.0 | 0.0 | 0.0 | 0.0 | 0.0 | 100.0 | 0.0 | 0.0 |
| 588 | 0.0 | 0.0 | 0.0 | 0.0 | 0.0 | 0.0 | 0.0 | 0.0 | 0.0 | 100.0 | 0.0 | 0.0 |
| 601 | 0.0 | 0.0 | 0.0 | 0.0 | 0.0 | 0.0 | 0.0 | 0.0 | 0.0 | 100.0 | 0.0 | 0.0 |
| 628 | 0.0 | 0.0 | 0.0 | 0.0 | 0.0 | 0.0 | 0.0 | 0.0 | 0.0 | 100.0 | 0.0 | 0.0 |
| 637 | 0.0 | 0.0 | 0.0 | 0.0 | 0.0 | 0.0 | 0.0 | 0.0 | 0.0 | 100.0 | 0.0 | 0.0 |
| 638 | 0.0 | 0.0 | 0.0 | 0.0 | 0.0 | 0.0 | 0.0 | 0.0 | 0.0 | 100.0 | 0.0 | 0.0 |
| 650 | 0.0 | 0.0 | 0.0 | 0.0 | 0.0 | 0.0 | 0.0 | 0.0 | 0.0 | 100.0 | 0.0 | 0.0 |
| 656 | 0.0 | 0.0 | 0.0 | 0.0 | 0.0 | 0.0 | 0.0 | 0.0 | 0.0 | 100.0 | 0.0 | 0.0 |
| 657 | 0.0 | 0.0 | 0.0 | 0.0 | 0.0 | 0.0 | 0.0 | 0.0 | 0.0 | 100.0 | 0.0 | 0.0 |
| 661 | 0.0 | 0.0 | 0.0 | 0.0 | 0.0 | 0.0 | 0.0 | 0.0 | 0.0 | 100.0 | 0.0 | 0.0 |
| 662 | 0.0 | 0.0 | 0.0 | 0.0 | 0.0 | 0.0 | 0.0 | 0.0 | 0.0 | 100.0 | 0.0 | 0.0 |
| 675 | 0.0 | 0.0 | 0.0 | 0.0 | 0.0 | 0.0 | 0.0 | 0.0 | 0.0 | 100.0 | 0.0 | 0.0 |
| 681 | 0.0 | 0.0 | 0.0 | 0.0 | 0.0 | 0.0 | 0.0 | 0.0 | 0.0 | 100.0 | 0.0 | 0.0 |
| 548 | 0.0 | 0.0 | 0.0 | 0.0 | 0.0 | 0.0 | 0.0 | 0.0 | 0.0 | 50.0 | 0.0 | 50.0 |
| 585 | 0.0 | 0.0 | 0.0 | 0.0 | 0.0 | 0.0 | 0.0 | 0.0 | 0.0 | 50.0 | 0.0 | 50.0 |
| 605 | 0.0 | 0.0 | 0.0 | 0.0 | 0.0 | 0.0 | 0.0 | 0.0 | 0.0 | 0.0 | 100.0 | 0.0 |
| 589 | 0.0 | 0.0 | 0.0 | 0.0 | 0.0 | 0.0 | 0.0 | 0.0 | 0.0 | 0.0 | 0.0 | 100.0 |
| 613 | 0.0 | 0.0 | 0.0 | 0.0 | 0.0 | 0.0 | 0.0 | 0.0 | 0.0 | 0.0 | 0.0 | 100.0 |
| 620 | 0.0 | 0.0 | 0.0 | 0.0 | 0.0 | 0.0 | 0.0 | 0.0 | 0.0 | 0.0 | 0.0 | 100.0 |
| 626 | 0.0 | 0.0 | 0.0 | 0.0 | 0.0 | 0.0 | 0.0 | 0.0 | 0.0 | 0.0 | 0.0 | 100.0 |
| 642 | 0.0 | 0.0 | 0.0 | 0.0 | 0.0 | 0.0 | 0.0 | 0.0 | 0.0 | 0.0 | 0.0 | 100.0 |
| 665 | 0.0 | 0.0 | 0.0 | 0.0 | 0.0 | 0.0 | 0.0 | 0.0 | 0.0 | 0.0 | 0.0 | 100.0 |
| 667 | 0.0 | 0.0 | 0.0 | 0.0 | 0.0 | 0.0 | 0.0 | 0.0 | 0.0 | 0.0 | 0.0 | 100.0 |GRN No.3
#### Chart
| Category | WT | 1塩基挿入 | 2塩基欠失 | 1塩基欠失-1 | 18塩基欠失 | 8塩基欠失 | 20塩基欠失 | 6塩基欠失 | 1塩基欠失-2 | NHEJ.others | HDR.NHEJ | HDR |
|---|---|---|---|---|---|---|---|---|---|---|---|---|
| 699 | 100.0 | 0.0 | 0.0 | 0.0 | 0.0 | 0.0 | 0.0 | 0.0 | 0.0 | 0.0 | 0.0 | 0.0 |
| 711 | 100.0 | 0.0 | 0.0 | 0.0 | 0.0 | 0.0 | 0.0 | 0.0 | 0.0 | 0.0 | 0.0 | 0.0 |
| 719 | 100.0 | 0.0 | 0.0 | 0.0 | 0.0 | 0.0 | 0.0 | 0.0 | 0.0 | 0.0 | 0.0 | 0.0 |
| 727 | 100.0 | 0.0 | 0.0 | 0.0 | 0.0 | 0.0 | 0.0 | 0.0 | 0.0 | 0.0 | 0.0 | 0.0 |
| 728 | 100.0 | 0.0 | 0.0 | 0.0 | 0.0 | 0.0 | 0.0 | 0.0 | 0.0 | 0.0 | 0.0 | 0.0 |
| 730 | 100.0 | 0.0 | 0.0 | 0.0 | 0.0 | 0.0 | 0.0 | 0.0 | 0.0 | 0.0 | 0.0 | 0.0 |
| 732 | 100.0 | 0.0 | 0.0 | 0.0 | 0.0 | 0.0 | 0.0 | 0.0 | 0.0 | 0.0 | 0.0 | 0.0 |
| 733 | 100.0 | 0.0 | 0.0 | 0.0 | 0.0 | 0.0 | 0.0 | 0.0 | 0.0 | 0.0 | 0.0 | 0.0 |
| 734 | 100.0 | 0.0 | 0.0 | 0.0 | 0.0 | 0.0 | 0.0 | 0.0 | 0.0 | 0.0 | 0.0 | 0.0 |
| 738 | 100.0 | 0.0 | 0.0 | 0.0 | 0.0 | 0.0 | 0.0 | 0.0 | 0.0 | 0.0 | 0.0 | 0.0 |
| 751 | 100.0 | 0.0 | 0.0 | 0.0 | 0.0 | 0.0 | 0.0 | 0.0 | 0.0 | 0.0 | 0.0 | 0.0 |
| 756 | 100.0 | 0.0 | 0.0 | 0.0 | 0.0 | 0.0 | 0.0 | 0.0 | 0.0 | 0.0 | 0.0 | 0.0 |
| 759 | 100.0 | 0.0 | 0.0 | 0.0 | 0.0 | 0.0 | 0.0 | 0.0 | 0.0 | 0.0 | 0.0 | 0.0 |
| 760 | 100.0 | 0.0 | 0.0 | 0.0 | 0.0 | 0.0 | 0.0 | 0.0 | 0.0 | 0.0 | 0.0 | 0.0 |
| 761 | 100.0 | 0.0 | 0.0 | 0.0 | 0.0 | 0.0 | 0.0 | 0.0 | 0.0 | 0.0 | 0.0 | 0.0 |
| 762 | 100.0 | 0.0 | 0.0 | 0.0 | 0.0 | 0.0 | 0.0 | 0.0 | 0.0 | 0.0 | 0.0 | 0.0 |
| 764 | 100.0 | 0.0 | 0.0 | 0.0 | 0.0 | 0.0 | 0.0 | 0.0 | 0.0 | 0.0 | 0.0 | 0.0 |
| 770 | 100.0 | 0.0 | 0.0 | 0.0 | 0.0 | 0.0 | 0.0 | 0.0 | 0.0 | 0.0 | 0.0 | 0.0 |
| 771 | 100.0 | 0.0 | 0.0 | 0.0 | 0.0 | 0.0 | 0.0 | 0.0 | 0.0 | 0.0 | 0.0 | 0.0 |
| 772 | 100.0 | 0.0 | 0.0 | 0.0 | 0.0 | 0.0 | 0.0 | 0.0 | 0.0 | 0.0 | 0.0 | 0.0 |
| 781 | 100.0 | 0.0 | 0.0 | 0.0 | 0.0 | 0.0 | 0.0 | 0.0 | 0.0 | 0.0 | 0.0 | 0.0 |
| 793 | 100.0 | 0.0 | 0.0 | 0.0 | 0.0 | 0.0 | 0.0 | 0.0 | 0.0 | 0.0 | 0.0 | 0.0 |
| 804 | 100.0 | 0.0 | 0.0 | 0.0 | 0.0 | 0.0 | 0.0 | 0.0 | 0.0 | 0.0 | 0.0 | 0.0 |
| 805 | 100.0 | 0.0 | 0.0 | 0.0 | 0.0 | 0.0 | 0.0 | 0.0 | 0.0 | 0.0 | 0.0 | 0.0 |
| 810 | 100.0 | 0.0 | 0.0 | 0.0 | 0.0 | 0.0 | 0.0 | 0.0 | 0.0 | 0.0 | 0.0 | 0.0 |
| 812 | 100.0 | 0.0 | 0.0 | 0.0 | 0.0 | 0.0 | 0.0 | 0.0 | 0.0 | 0.0 | 0.0 | 0.0 |
| 818 | 100.0 | 0.0 | 0.0 | 0.0 | 0.0 | 0.0 | 0.0 | 0.0 | 0.0 | 0.0 | 0.0 | 0.0 |
| 819 | 100.0 | 0.0 | 0.0 | 0.0 | 0.0 | 0.0 | 0.0 | 0.0 | 0.0 | 0.0 | 0.0 | 0.0 |
| 830 | 100.0 | 0.0 | 0.0 | 0.0 | 0.0 | 0.0 | 0.0 | 0.0 | 0.0 | 0.0 | 0.0 | 0.0 |
| 831 | 100.0 | 0.0 | 0.0 | 0.0 | 0.0 | 0.0 | 0.0 | 0.0 | 0.0 | 0.0 | 0.0 | 0.0 |
| 834 | 100.0 | 0.0 | 0.0 | 0.0 | 0.0 | 0.0 | 0.0 | 0.0 | 0.0 | 0.0 | 0.0 | 0.0 |
| 835 | 100.0 | 0.0 | 0.0 | 0.0 | 0.0 | 0.0 | 0.0 | 0.0 | 0.0 | 0.0 | 0.0 | 0.0 |
| 790 | 100.0 | 0.0 | 0.0 | 0.0 | 0.0 | 0.0 | 0.0 | 0.0 | 0.0 | 0.0 | 0.0 | 0.0 |
| 709 | 50.0 | 50.0 | 0.0 | 0.0 | 0.0 | 0.0 | 0.0 | 0.0 | 0.0 | 0.0 | 0.0 | 0.0 |
| 746 | 50.0 | 50.0 | 0.0 | 0.0 | 0.0 | 0.0 | 0.0 | 0.0 | 0.0 | 0.0 | 0.0 | 0.0 |
| 723 | 50.0 | 0.0 | 0.0 | 50.0 | 0.0 | 0.0 | 0.0 | 0.0 | 0.0 | 0.0 | 0.0 | 0.0 |
| 758 | 50.0 | 0.0 | 0.0 | 50.0 | 0.0 | 0.0 | 0.0 | 0.0 | 0.0 | 0.0 | 0.0 | 0.0 |
| 787 | 50.0 | 0.0 | 0.0 | 50.0 | 0.0 | 0.0 | 0.0 | 0.0 | 0.0 | 0.0 | 0.0 | 0.0 |
| 717 | 50.0 | 0.0 | 0.0 | 0.0 | 50.0 | 0.0 | 0.0 | 0.0 | 0.0 | 0.0 | 0.0 | 0.0 |
| 768 | 50.0 | 0.0 | 0.0 | 0.0 | 50.0 | 0.0 | 0.0 | 0.0 | 0.0 | 0.0 | 0.0 | 0.0 |
| 807 | 50.0 | 0.0 | 0.0 | 0.0 | 0.0 | 0.0 | 0.0 | 50.0 | 0.0 | 0.0 | 0.0 | 0.0 |
| 823 | 50.0 | 0.0 | 0.0 | 0.0 | 0.0 | 0.0 | 0.0 | 50.0 | 0.0 | 0.0 | 0.0 | 0.0 |
| 706 | 50.0 | 0.0 | 0.0 | 0.0 | 0.0 | 0.0 | 0.0 | 0.0 | 0.0 | 50.0 | 0.0 | 0.0 |
| 718 | 50.0 | 0.0 | 0.0 | 0.0 | 0.0 | 0.0 | 0.0 | 0.0 | 0.0 | 50.0 | 0.0 | 0.0 |
| 726 | 50.0 | 0.0 | 0.0 | 0.0 | 0.0 | 0.0 | 0.0 | 0.0 | 0.0 | 50.0 | 0.0 | 0.0 |
| 740 | 50.0 | 0.0 | 0.0 | 0.0 | 0.0 | 0.0 | 0.0 | 0.0 | 0.0 | 50.0 | 0.0 | 0.0 |
| 742 | 50.0 | 0.0 | 0.0 | 0.0 | 0.0 | 0.0 | 0.0 | 0.0 | 0.0 | 50.0 | 0.0 | 0.0 |
| 748 | 50.0 | 0.0 | 0.0 | 0.0 | 0.0 | 0.0 | 0.0 | 0.0 | 0.0 | 50.0 | 0.0 | 0.0 |
| 752 | 50.0 | 0.0 | 0.0 | 0.0 | 0.0 | 0.0 | 0.0 | 0.0 | 0.0 | 50.0 | 0.0 | 0.0 |
| 773 | 50.0 | 0.0 | 0.0 | 0.0 | 0.0 | 0.0 | 0.0 | 0.0 | 0.0 | 50.0 | 0.0 | 0.0 |
| 786 | 50.0 | 0.0 | 0.0 | 0.0 | 0.0 | 0.0 | 0.0 | 0.0 | 0.0 | 50.0 | 0.0 | 0.0 |
| 789 | 50.0 | 0.0 | 0.0 | 0.0 | 0.0 | 0.0 | 0.0 | 0.0 | 0.0 | 50.0 | 0.0 | 0.0 |
| 827 | 50.0 | 0.0 | 0.0 | 0.0 | 0.0 | 0.0 | 0.0 | 0.0 | 0.0 | 50.0 | 0.0 | 0.0 |
| 701 | 0.0 | 100.0 | 0.0 | 0.0 | 0.0 | 0.0 | 0.0 | 0.0 | 0.0 | 0.0 | 0.0 | 0.0 |
| 767 | 0.0 | 100.0 | 0.0 | 0.0 | 0.0 | 0.0 | 0.0 | 0.0 | 0.0 | 0.0 | 0.0 | 0.0 |
| 780 | 0.0 | 50.0 | 0.0 | 50.0 | 0.0 | 0.0 | 0.0 | 0.0 | 0.0 | 0.0 | 0.0 | 0.0 |
| 776 | 0.0 | 50.0 | 0.0 | 0.0 | 0.0 | 0.0 | 50.0 | 0.0 | 0.0 | 0.0 | 0.0 | 0.0 |
| 792 | 0.0 | 50.0 | 0.0 | 0.0 | 0.0 | 0.0 | 50.0 | 0.0 | 0.0 | 0.0 | 0.0 | 0.0 |
| 798 | 0.0 | 50.0 | 0.0 | 0.0 | 0.0 | 0.0 | 50.0 | 0.0 | 0.0 | 0.0 | 0.0 | 0.0 |
| 779 | 0.0 | 50.0 | 0.0 | 0.0 | 0.0 | 0.0 | 0.0 | 0.0 | 50.0 | 0.0 | 0.0 | 0.0 |
| 700 | 0.0 | 50.0 | 0.0 | 0.0 | 0.0 | 0.0 | 0.0 | 0.0 | 0.0 | 50.0 | 0.0 | 0.0 |
| 715 | 0.0 | 50.0 | 0.0 | 0.0 | 0.0 | 0.0 | 0.0 | 0.0 | 0.0 | 50.0 | 0.0 | 0.0 |
| 785 | 0.0 | 50.0 | 0.0 | 0.0 | 0.0 | 0.0 | 0.0 | 0.0 | 0.0 | 50.0 | 0.0 | 0.0 |
| 822 | 0.0 | 50.0 | 0.0 | 0.0 | 0.0 | 0.0 | 0.0 | 0.0 | 0.0 | 50.0 | 0.0 | 0.0 |
| 829 | 0.0 | 50.0 | 0.0 | 0.0 | 0.0 | 0.0 | 0.0 | 0.0 | 0.0 | 50.0 | 0.0 | 0.0 |
| 743 | 0.0 | 50.0 | 0.0 | 0.0 | 0.0 | 0.0 | 0.0 | 0.0 | 0.0 | 0.0 | 0.0 | 50.0 |
| 777 | 0.0 | 0.0 | 50.0 | 0.0 | 0.0 | 0.0 | 0.0 | 0.0 | 0.0 | 0.0 | 50.0 | 0.0 |
| 745 | 0.0 | 0.0 | 0.0 | 50.0 | 0.0 | 0.0 | 0.0 | 0.0 | 0.0 | 50.0 | 0.0 | 0.0 |
| 828 | 0.0 | 0.0 | 0.0 | 0.0 | 100.0 | 0.0 | 0.0 | 0.0 | 0.0 | 0.0 | 0.0 | 0.0 |
| 755 | 0.0 | 0.0 | 0.0 | 0.0 | 50.0 | 0.0 | 0.0 | 0.0 | 0.0 | 50.0 | 0.0 | 0.0 |
| 766 | 0.0 | 0.0 | 0.0 | 0.0 | 50.0 | 0.0 | 0.0 | 0.0 | 0.0 | 50.0 | 0.0 | 0.0 |
| 769 | 0.0 | 0.0 | 0.0 | 0.0 | 50.0 | 0.0 | 0.0 | 0.0 | 0.0 | 50.0 | 0.0 | 0.0 |
| 783 | 0.0 | 0.0 | 0.0 | 0.0 | 50.0 | 0.0 | 0.0 | 0.0 | 0.0 | 50.0 | 0.0 | 0.0 |
| 791 | 0.0 | 0.0 | 0.0 | 0.0 | 50.0 | 0.0 | 0.0 | 0.0 | 0.0 | 50.0 | 0.0 | 0.0 |
| 796 | 0.0 | 0.0 | 0.0 | 0.0 | 50.0 | 0.0 | 0.0 | 0.0 | 0.0 | 50.0 | 0.0 | 0.0 |
| 813 | 0.0 | 0.0 | 0.0 | 0.0 | 50.0 | 0.0 | 0.0 | 0.0 | 0.0 | 50.0 | 0.0 | 0.0 |
| 817 | 0.0 | 0.0 | 0.0 | 0.0 | 50.0 | 0.0 | 0.0 | 0.0 | 0.0 | 50.0 | 0.0 | 0.0 |
| 731 | 0.0 | 0.0 | 0.0 | 0.0 | 50.0 | 0.0 | 0.0 | 0.0 | 0.0 | 0.0 | 0.0 | 50.0 |
| 710 | 0.0 | 0.0 | 0.0 | 0.0 | 0.0 | 100.0 | 0.0 | 0.0 | 0.0 | 0.0 | 0.0 | 0.0 |
| 815 | 0.0 | 0.0 | 0.0 | 0.0 | 0.0 | 100.0 | 0.0 | 0.0 | 0.0 | 0.0 | 0.0 | 0.0 |
| 707 | 0.0 | 0.0 | 0.0 | 0.0 | 0.0 | 50.0 | 0.0 | 0.0 | 0.0 | 50.0 | 0.0 | 0.0 |
| 716 | 0.0 | 0.0 | 0.0 | 0.0 | 0.0 | 50.0 | 0.0 | 0.0 | 0.0 | 50.0 | 0.0 | 0.0 |
| 784 | 0.0 | 0.0 | 0.0 | 0.0 | 0.0 | 50.0 | 0.0 | 0.0 | 0.0 | 50.0 | 0.0 | 0.0 |
| 788 | 0.0 | 0.0 | 0.0 | 0.0 | 0.0 | 50.0 | 0.0 | 0.0 | 0.0 | 50.0 | 0.0 | 0.0 |
| 797 | 0.0 | 0.0 | 0.0 | 0.0 | 0.0 | 50.0 | 0.0 | 0.0 | 0.0 | 50.0 | 0.0 | 0.0 |
| 720 | 0.0 | 0.0 | 0.0 | 0.0 | 0.0 | 0.0 | 0.0 | 100.0 | 0.0 | 0.0 | 0.0 | 0.0 |
| 749 | 0.0 | 0.0 | 0.0 | 0.0 | 0.0 | 0.0 | 0.0 | 50.0 | 0.0 | 50.0 | 0.0 | 0.0 |
| 765 | 0.0 | 0.0 | 0.0 | 0.0 | 0.0 | 0.0 | 0.0 | 50.0 | 0.0 | 50.0 | 0.0 | 0.0 |
| 702 | 0.0 | 0.0 | 0.0 | 0.0 | 0.0 | 0.0 | 0.0 | 0.0 | 0.0 | 100.0 | 0.0 | 0.0 |
| 703 | 0.0 | 0.0 | 0.0 | 0.0 | 0.0 | 0.0 | 0.0 | 0.0 | 0.0 | 100.0 | 0.0 | 0.0 |
| 708 | 0.0 | 0.0 | 0.0 | 0.0 | 0.0 | 0.0 | 0.0 | 0.0 | 0.0 | 100.0 | 0.0 | 0.0 |
| 712 | 0.0 | 0.0 | 0.0 | 0.0 | 0.0 | 0.0 | 0.0 | 0.0 | 0.0 | 100.0 | 0.0 | 0.0 |
| 714 | 0.0 | 0.0 | 0.0 | 0.0 | 0.0 | 0.0 | 0.0 | 0.0 | 0.0 | 100.0 | 0.0 | 0.0 |
| 724 | 0.0 | 0.0 | 0.0 | 0.0 | 0.0 | 0.0 | 0.0 | 0.0 | 0.0 | 100.0 | 0.0 | 0.0 |
| 725 | 0.0 | 0.0 | 0.0 | 0.0 | 0.0 | 0.0 | 0.0 | 0.0 | 0.0 | 100.0 | 0.0 | 0.0 |
| 729 | 0.0 | 0.0 | 0.0 | 0.0 | 0.0 | 0.0 | 0.0 | 0.0 | 0.0 | 100.0 | 0.0 | 0.0 |
| 735 | 0.0 | 0.0 | 0.0 | 0.0 | 0.0 | 0.0 | 0.0 | 0.0 | 0.0 | 100.0 | 0.0 | 0.0 |
| 741 | 0.0 | 0.0 | 0.0 | 0.0 | 0.0 | 0.0 | 0.0 | 0.0 | 0.0 | 100.0 | 0.0 | 0.0 |
| 744 | 0.0 | 0.0 | 0.0 | 0.0 | 0.0 | 0.0 | 0.0 | 0.0 | 0.0 | 100.0 | 0.0 | 0.0 |
| 747 | 0.0 | 0.0 | 0.0 | 0.0 | 0.0 | 0.0 | 0.0 | 0.0 | 0.0 | 100.0 | 0.0 | 0.0 |
| 750 | 0.0 | 0.0 | 0.0 | 0.0 | 0.0 | 0.0 | 0.0 | 0.0 | 0.0 | 100.0 | 0.0 | 0.0 |
| 753 | 0.0 | 0.0 | 0.0 | 0.0 | 0.0 | 0.0 | 0.0 | 0.0 | 0.0 | 100.0 | 0.0 | 0.0 |
| 754 | 0.0 | 0.0 | 0.0 | 0.0 | 0.0 | 0.0 | 0.0 | 0.0 | 0.0 | 100.0 | 0.0 | 0.0 |
| 757 | 0.0 | 0.0 | 0.0 | 0.0 | 0.0 | 0.0 | 0.0 | 0.0 | 0.0 | 100.0 | 0.0 | 0.0 |
| 763 | 0.0 | 0.0 | 0.0 | 0.0 | 0.0 | 0.0 | 0.0 | 0.0 | 0.0 | 100.0 | 0.0 | 0.0 |
| 774 | 0.0 | 0.0 | 0.0 | 0.0 | 0.0 | 0.0 | 0.0 | 0.0 | 0.0 | 100.0 | 0.0 | 0.0 |
| 775 | 0.0 | 0.0 | 0.0 | 0.0 | 0.0 | 0.0 | 0.0 | 0.0 | 0.0 | 100.0 | 0.0 | 0.0 |
| 778 | 0.0 | 0.0 | 0.0 | 0.0 | 0.0 | 0.0 | 0.0 | 0.0 | 0.0 | 100.0 | 0.0 | 0.0 |
| 794 | 0.0 | 0.0 | 0.0 | 0.0 | 0.0 | 0.0 | 0.0 | 0.0 | 0.0 | 100.0 | 0.0 | 0.0 |
| 795 | 0.0 | 0.0 | 0.0 | 0.0 | 0.0 | 0.0 | 0.0 | 0.0 | 0.0 | 100.0 | 0.0 | 0.0 |
| 806 | 0.0 | 0.0 | 0.0 | 0.0 | 0.0 | 0.0 | 0.0 | 0.0 | 0.0 | 100.0 | 0.0 | 0.0 |
| 811 | 0.0 | 0.0 | 0.0 | 0.0 | 0.0 | 0.0 | 0.0 | 0.0 | 0.0 | 100.0 | 0.0 | 0.0 |
| 814 | 0.0 | 0.0 | 0.0 | 0.0 | 0.0 | 0.0 | 0.0 | 0.0 | 0.0 | 100.0 | 0.0 | 0.0 |
| 782 | 0.0 | 0.0 | 0.0 | 0.0 | 0.0 | 0.0 | 0.0 | 0.0 | 0.0 | 50.0 | 0.0 | 50.0 |C
WT
1-bp ins
Δ2-bp
Δ1-bp_1
Δ18-bp
Δ8-bp
Δ20-bp
6-bp
Δ1-bp_2
NHEJ.others
NHEJ+HDR
HDR
Model for GRN No.1
#### Chart
| Category | WT | 1-bp ins | Δ2-bp | Δ1-bp_1 | Δ18-bp | Δ8-bp | Δ20-bp | Δ6-bp | Δ1-bp_2 | NHEJ.others | NHEJ+HDR | HDR |
|---|---|---|---|---|---|---|---|---|---|---|---|---|Model for GRN No.2
#### Chart
| Category | WT | 1-bp ins | Δ2-bp | Δ1-bp_1 | Δ18-bp | Δ8-bp | Δ20-bp | Δ6-bp | Δ1-bp_2 | NHEJ.others | NHEJ+HDR | HDR |
|---|---|---|---|---|---|---|---|---|---|---|---|---|Model for GRN No.3
#### Chart
| Category | WT | 1-bp ins | Δ2-bp | Δ1-bp_1 | Δ18-bp | Δ8-bp | Δ20-bp | Δ6-bp | Δ1-bp_2 | NHEJ.others | NHEJ+HDR | HDR |
|---|---|---|---|---|---|---|---|---|---|---|---|---|

### Slide 7
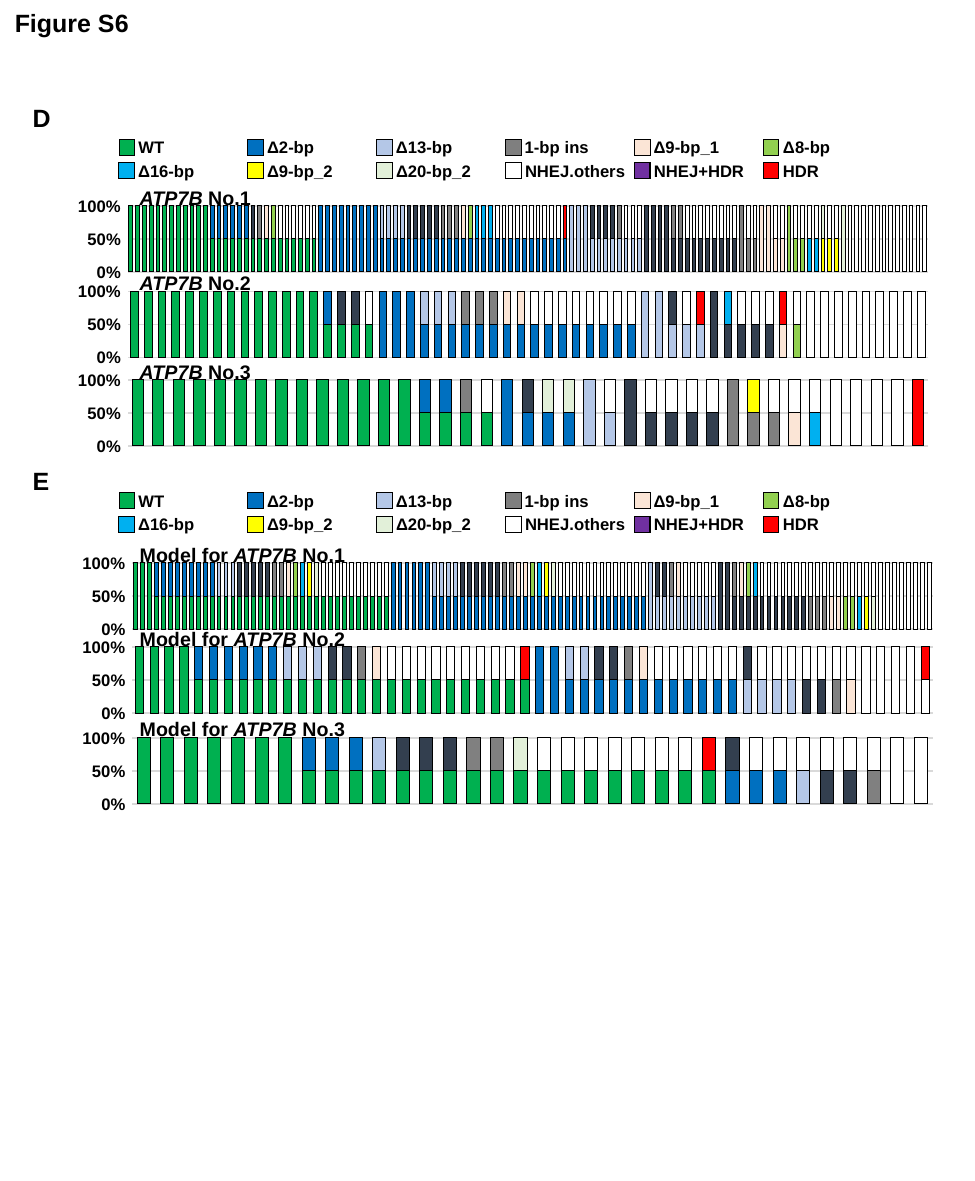

Figure S6
D
WT
Δ2-bp
Δ13-bp
1-bp ins
Δ9-bp_1
Δ8-bp
Δ16-bp
Δ9-bp_2
Δ20-bp_2
NHEJ.others
NHEJ+HDR
HDR
ATP7B No.1
#### Chart
| Category | WT | 2塩基欠失 | 13塩基欠失 | 1塩基挿入 | 9塩基欠失-1 | 8塩基欠失 | 16塩基欠失 | 9塩基欠失-2 | 20塩基欠失 | 11塩基欠失 | NHEJ.others | HDR |
|---|---|---|---|---|---|---|---|---|---|---|---|---|
| 9 | 100.0 | 0.0 | 0.0 | 0.0 | 0.0 | 0.0 | 0.0 | 0.0 | 0.0 | 0.0 | 0.0 | 0.0 |
| 12 | 100.0 | 0.0 | 0.0 | 0.0 | 0.0 | 0.0 | 0.0 | 0.0 | 0.0 | 0.0 | 0.0 | 0.0 |
| 57 | 100.0 | 0.0 | 0.0 | 0.0 | 0.0 | 0.0 | 0.0 | 0.0 | 0.0 | 0.0 | 0.0 | 0.0 |
| 60 | 100.0 | 0.0 | 0.0 | 0.0 | 0.0 | 0.0 | 0.0 | 0.0 | 0.0 | 0.0 | 0.0 | 0.0 |
| 81 | 100.0 | 0.0 | 0.0 | 0.0 | 0.0 | 0.0 | 0.0 | 0.0 | 0.0 | 0.0 | 0.0 | 0.0 |
| 140 | 100.0 | 0.0 | 0.0 | 0.0 | 0.0 | 0.0 | 0.0 | 0.0 | 0.0 | 0.0 | 0.0 | 0.0 |
| 153 | 100.0 | 0.0 | 0.0 | 0.0 | 0.0 | 0.0 | 0.0 | 0.0 | 0.0 | 0.0 | 0.0 | 0.0 |
| 169 | 100.0 | 0.0 | 0.0 | 0.0 | 0.0 | 0.0 | 0.0 | 0.0 | 0.0 | 0.0 | 0.0 | 0.0 |
| 172 | 100.0 | 0.0 | 0.0 | 0.0 | 0.0 | 0.0 | 0.0 | 0.0 | 0.0 | 0.0 | 0.0 | 0.0 |
| 175 | 100.0 | 0.0 | 0.0 | 0.0 | 0.0 | 0.0 | 0.0 | 0.0 | 0.0 | 0.0 | 0.0 | 0.0 |
| 179 | 100.0 | 0.0 | 0.0 | 0.0 | 0.0 | 0.0 | 0.0 | 0.0 | 0.0 | 0.0 | 0.0 | 0.0 |
| 10 | 100.0 | 0.0 | 0.0 | 0.0 | 0.0 | 0.0 | 0.0 | 0.0 | 0.0 | 0.0 | 0.0 | 0.0 |
| 25 | 50.0 | 50.0 | 0.0 | 0.0 | 0.0 | 0.0 | 0.0 | 0.0 | 0.0 | 0.0 | 0.0 | 0.0 |
| 52 | 50.0 | 50.0 | 0.0 | 0.0 | 0.0 | 0.0 | 0.0 | 0.0 | 0.0 | 0.0 | 0.0 | 0.0 |
| 109 | 50.0 | 50.0 | 0.0 | 0.0 | 0.0 | 0.0 | 0.0 | 0.0 | 0.0 | 0.0 | 0.0 | 0.0 |
| 166 | 50.0 | 50.0 | 0.0 | 0.0 | 0.0 | 0.0 | 0.0 | 0.0 | 0.0 | 0.0 | 0.0 | 0.0 |
| 178 | 50.0 | 50.0 | 0.0 | 0.0 | 0.0 | 0.0 | 0.0 | 0.0 | 0.0 | 0.0 | 0.0 | 0.0 |
| 84 | 50.0 | 50.0 | 0.0 | 0.0 | 0.0 | 0.0 | 0.0 | 0.0 | 0.0 | 0.0 | 0.0 | 0.0 |
| 17 | 50.0 | 0.0 | 0.0 | 50.0 | 0.0 | 0.0 | 0.0 | 0.0 | 0.0 | 0.0 | 0.0 | 0.0 |
| 19 | 50.0 | 0.0 | 0.0 | 0.0 | 50.0 | 0.0 | 0.0 | 0.0 | 0.0 | 0.0 | 0.0 | 0.0 |
| 45 | 50.0 | 0.0 | 0.0 | 0.0 | 0.0 | 50.0 | 0.0 | 0.0 | 0.0 | 0.0 | 0.0 | 0.0 |
| 97 | 50.0 | 0.0 | 0.0 | 0.0 | 0.0 | 0.0 | 50.0 | 0.0 | 0.0 | 0.0 | 0.0 | 0.0 |
| 118 | 50.0 | 0.0 | 0.0 | 0.0 | 0.0 | 0.0 | 0.0 | 0.0 | 0.0 | 0.0 | 50.0 | 0.0 |
| 142 | 50.0 | 0.0 | 0.0 | 0.0 | 0.0 | 0.0 | 0.0 | 0.0 | 0.0 | 0.0 | 50.0 | 0.0 |
| 53 | 50.0 | 0.0 | 0.0 | 0.0 | 0.0 | 0.0 | 0.0 | 0.0 | 0.0 | 0.0 | 50.0 | 0.0 |
| 75 | 50.0 | 0.0 | 0.0 | 0.0 | 0.0 | 0.0 | 0.0 | 0.0 | 0.0 | 0.0 | 50.0 | 0.0 |
| 105 | 50.0 | 0.0 | 0.0 | 0.0 | 0.0 | 0.0 | 0.0 | 0.0 | 0.0 | 0.0 | 50.0 | 0.0 |
| 173 | 50.0 | 0.0 | 0.0 | 0.0 | 0.0 | 0.0 | 0.0 | 0.0 | 0.0 | 0.0 | 50.0 | 0.0 |
| 15 | 0.0 | 100.0 | 0.0 | 0.0 | 0.0 | 0.0 | 0.0 | 0.0 | 0.0 | 0.0 | 0.0 | 0.0 |
| 34 | 0.0 | 100.0 | 0.0 | 0.0 | 0.0 | 0.0 | 0.0 | 0.0 | 0.0 | 0.0 | 0.0 | 0.0 |
| 36 | 0.0 | 100.0 | 0.0 | 0.0 | 0.0 | 0.0 | 0.0 | 0.0 | 0.0 | 0.0 | 0.0 | 0.0 |
| 49 | 0.0 | 100.0 | 0.0 | 0.0 | 0.0 | 0.0 | 0.0 | 0.0 | 0.0 | 0.0 | 0.0 | 0.0 |
| 56 | 0.0 | 100.0 | 0.0 | 0.0 | 0.0 | 0.0 | 0.0 | 0.0 | 0.0 | 0.0 | 0.0 | 0.0 |
| 110 | 0.0 | 100.0 | 0.0 | 0.0 | 0.0 | 0.0 | 0.0 | 0.0 | 0.0 | 0.0 | 0.0 | 0.0 |
| 157 | 0.0 | 100.0 | 0.0 | 0.0 | 0.0 | 0.0 | 0.0 | 0.0 | 0.0 | 0.0 | 0.0 | 0.0 |
| 65 | 0.0 | 100.0 | 0.0 | 0.0 | 0.0 | 0.0 | 0.0 | 0.0 | 0.0 | 0.0 | 0.0 | 0.0 |
| 154 | 0.0 | 100.0 | 0.0 | 0.0 | 0.0 | 0.0 | 0.0 | 0.0 | 0.0 | 0.0 | 0.0 | 0.0 |
| 2 | 0.0 | 50.0 | 50.0 | 0.0 | 0.0 | 0.0 | 0.0 | 0.0 | 0.0 | 0.0 | 0.0 | 0.0 |
| 79 | 0.0 | 50.0 | 50.0 | 0.0 | 0.0 | 0.0 | 0.0 | 0.0 | 0.0 | 0.0 | 0.0 | 0.0 |
| 83 | 0.0 | 50.0 | 50.0 | 0.0 | 0.0 | 0.0 | 0.0 | 0.0 | 0.0 | 0.0 | 0.0 | 0.0 |
| 101 | 0.0 | 50.0 | 50.0 | 0.0 | 0.0 | 0.0 | 0.0 | 0.0 | 0.0 | 0.0 | 0.0 | 0.0 |
| 32 | 0.0 | 50.0 | 0.0 | 50.0 | 0.0 | 0.0 | 0.0 | 0.0 | 0.0 | 0.0 | 0.0 | 0.0 |
| 43 | 0.0 | 50.0 | 0.0 | 50.0 | 0.0 | 0.0 | 0.0 | 0.0 | 0.0 | 0.0 | 0.0 | 0.0 |
| 68 | 0.0 | 50.0 | 0.0 | 50.0 | 0.0 | 0.0 | 0.0 | 0.0 | 0.0 | 0.0 | 0.0 | 0.0 |
| 72 | 0.0 | 50.0 | 0.0 | 50.0 | 0.0 | 0.0 | 0.0 | 0.0 | 0.0 | 0.0 | 0.0 | 0.0 |
| 107 | 0.0 | 50.0 | 0.0 | 50.0 | 0.0 | 0.0 | 0.0 | 0.0 | 0.0 | 0.0 | 0.0 | 0.0 |
| 121 | 0.0 | 50.0 | 0.0 | 0.0 | 50.0 | 0.0 | 0.0 | 0.0 | 0.0 | 0.0 | 0.0 | 0.0 |
| 96 | 0.0 | 50.0 | 0.0 | 0.0 | 50.0 | 0.0 | 0.0 | 0.0 | 0.0 | 0.0 | 0.0 | 0.0 |
| 13 | 0.0 | 50.0 | 0.0 | 0.0 | 50.0 | 0.0 | 0.0 | 0.0 | 0.0 | 0.0 | 0.0 | 0.0 |
| 177 | 0.0 | 50.0 | 0.0 | 0.0 | 0.0 | 50.0 | 0.0 | 0.0 | 0.0 | 0.0 | 0.0 | 0.0 |
| 69 | 0.0 | 50.0 | 0.0 | 0.0 | 0.0 | 0.0 | 50.0 | 0.0 | 0.0 | 0.0 | 0.0 | 0.0 |
| 27 | 0.0 | 50.0 | 0.0 | 0.0 | 0.0 | 0.0 | 0.0 | 50.0 | 0.0 | 0.0 | 0.0 | 0.0 |
| 126 | 0.0 | 50.0 | 0.0 | 0.0 | 0.0 | 0.0 | 0.0 | 50.0 | 0.0 | 0.0 | 0.0 | 0.0 |
| 152 | 0.0 | 50.0 | 0.0 | 0.0 | 0.0 | 0.0 | 0.0 | 50.0 | 0.0 | 0.0 | 0.0 | 0.0 |
| 14 | 0.0 | 50.0 | 0.0 | 0.0 | 0.0 | 0.0 | 0.0 | 0.0 | 0.0 | 0.0 | 50.0 | 0.0 |
| 63 | 0.0 | 50.0 | 0.0 | 0.0 | 0.0 | 0.0 | 0.0 | 0.0 | 0.0 | 0.0 | 50.0 | 0.0 |
| 35 | 0.0 | 50.0 | 0.0 | 0.0 | 0.0 | 0.0 | 0.0 | 0.0 | 0.0 | 0.0 | 50.0 | 0.0 |
| 64 | 0.0 | 50.0 | 0.0 | 0.0 | 0.0 | 0.0 | 0.0 | 0.0 | 0.0 | 0.0 | 50.0 | 0.0 |
| 102 | 0.0 | 50.0 | 0.0 | 0.0 | 0.0 | 0.0 | 0.0 | 0.0 | 0.0 | 0.0 | 50.0 | 0.0 |
| 165 | 0.0 | 50.0 | 0.0 | 0.0 | 0.0 | 0.0 | 0.0 | 0.0 | 0.0 | 0.0 | 50.0 | 0.0 |
| 59 | 0.0 | 50.0 | 0.0 | 0.0 | 0.0 | 0.0 | 0.0 | 0.0 | 0.0 | 0.0 | 50.0 | 0.0 |
| 33 | 0.0 | 50.0 | 0.0 | 0.0 | 0.0 | 0.0 | 0.0 | 0.0 | 0.0 | 0.0 | 50.0 | 0.0 |
| 143 | 0.0 | 50.0 | 0.0 | 0.0 | 0.0 | 0.0 | 0.0 | 0.0 | 0.0 | 0.0 | 50.0 | 0.0 |
| 145 | 0.0 | 50.0 | 0.0 | 0.0 | 0.0 | 0.0 | 0.0 | 0.0 | 0.0 | 0.0 | 50.0 | 0.0 |
| 38 | 0.0 | 50.0 | 0.0 | 0.0 | 0.0 | 0.0 | 0.0 | 0.0 | 0.0 | 0.0 | 0.0 | 50.0 |
| 47 | 0.0 | 0.0 | 100.0 | 0.0 | 0.0 | 0.0 | 0.0 | 0.0 | 0.0 | 0.0 | 0.0 | 0.0 |
| 130 | 0.0 | 0.0 | 100.0 | 0.0 | 0.0 | 0.0 | 0.0 | 0.0 | 0.0 | 0.0 | 0.0 | 0.0 |
| 136 | 0.0 | 0.0 | 100.0 | 0.0 | 0.0 | 0.0 | 0.0 | 0.0 | 0.0 | 0.0 | 0.0 | 0.0 |
| 86 | 0.0 | 0.0 | 50.0 | 50.0 | 0.0 | 0.0 | 0.0 | 0.0 | 0.0 | 0.0 | 0.0 | 0.0 |
| 125 | 0.0 | 0.0 | 50.0 | 50.0 | 0.0 | 0.0 | 0.0 | 0.0 | 0.0 | 0.0 | 0.0 | 0.0 |
| 132 | 0.0 | 0.0 | 50.0 | 50.0 | 0.0 | 0.0 | 0.0 | 0.0 | 0.0 | 0.0 | 0.0 | 0.0 |
| 134 | 0.0 | 0.0 | 50.0 | 50.0 | 0.0 | 0.0 | 0.0 | 0.0 | 0.0 | 0.0 | 0.0 | 0.0 |
| 133 | 0.0 | 0.0 | 50.0 | 0.0 | 50.0 | 0.0 | 0.0 | 0.0 | 0.0 | 0.0 | 0.0 | 0.0 |
| 137 | 0.0 | 0.0 | 50.0 | 0.0 | 0.0 | 0.0 | 0.0 | 0.0 | 0.0 | 0.0 | 50.0 | 0.0 |
| 4 | 0.0 | 0.0 | 50.0 | 0.0 | 0.0 | 0.0 | 0.0 | 0.0 | 0.0 | 0.0 | 50.0 | 0.0 |
| 128 | 0.0 | 0.0 | 50.0 | 0.0 | 0.0 | 0.0 | 0.0 | 0.0 | 0.0 | 0.0 | 50.0 | 0.0 |
| 61 | 0.0 | 0.0 | 0.0 | 100.0 | 0.0 | 0.0 | 0.0 | 0.0 | 0.0 | 0.0 | 0.0 | 0.0 |
| 89 | 0.0 | 0.0 | 0.0 | 100.0 | 0.0 | 0.0 | 0.0 | 0.0 | 0.0 | 0.0 | 0.0 | 0.0 |
| 131 | 0.0 | 0.0 | 0.0 | 100.0 | 0.0 | 0.0 | 0.0 | 0.0 | 0.0 | 0.0 | 0.0 | 0.0 |
| 167 | 0.0 | 0.0 | 0.0 | 100.0 | 0.0 | 0.0 | 0.0 | 0.0 | 0.0 | 0.0 | 0.0 | 0.0 |
| 123 | 0.0 | 0.0 | 0.0 | 50.0 | 50.0 | 0.0 | 0.0 | 0.0 | 0.0 | 0.0 | 0.0 | 0.0 |
| 77 | 0.0 | 0.0 | 0.0 | 50.0 | 50.0 | 0.0 | 0.0 | 0.0 | 0.0 | 0.0 | 0.0 | 0.0 |
| 51 | 0.0 | 0.0 | 0.0 | 50.0 | 0.0 | 0.0 | 0.0 | 0.0 | 0.0 | 0.0 | 50.0 | 0.0 |
| 87 | 0.0 | 0.0 | 0.0 | 50.0 | 0.0 | 0.0 | 0.0 | 0.0 | 0.0 | 0.0 | 50.0 | 0.0 |
| 108 | 0.0 | 0.0 | 0.0 | 50.0 | 0.0 | 0.0 | 0.0 | 0.0 | 0.0 | 0.0 | 50.0 | 0.0 |
| 111 | 0.0 | 0.0 | 0.0 | 50.0 | 0.0 | 0.0 | 0.0 | 0.0 | 0.0 | 0.0 | 50.0 | 0.0 |
| 113 | 0.0 | 0.0 | 0.0 | 50.0 | 0.0 | 0.0 | 0.0 | 0.0 | 0.0 | 0.0 | 50.0 | 0.0 |
| 147 | 0.0 | 0.0 | 0.0 | 50.0 | 0.0 | 0.0 | 0.0 | 0.0 | 0.0 | 0.0 | 50.0 | 0.0 |
| 58 | 0.0 | 0.0 | 0.0 | 50.0 | 0.0 | 0.0 | 0.0 | 0.0 | 0.0 | 0.0 | 50.0 | 0.0 |
| 141 | 0.0 | 0.0 | 0.0 | 50.0 | 0.0 | 0.0 | 0.0 | 0.0 | 0.0 | 0.0 | 50.0 | 0.0 |
| 117 | 0.0 | 0.0 | 0.0 | 0.0 | 100.0 | 0.0 | 0.0 | 0.0 | 0.0 | 0.0 | 0.0 | 0.0 |
| 20 | 0.0 | 0.0 | 0.0 | 0.0 | 50.0 | 0.0 | 0.0 | 0.0 | 0.0 | 0.0 | 50.0 | 0.0 |
| 171 | 0.0 | 0.0 | 0.0 | 0.0 | 50.0 | 0.0 | 0.0 | 0.0 | 0.0 | 0.0 | 50.0 | 0.0 |
| 106 | 0.0 | 0.0 | 0.0 | 0.0 | 0.0 | 100.0 | 0.0 | 0.0 | 0.0 | 0.0 | 0.0 | 0.0 |
| 144 | 0.0 | 0.0 | 0.0 | 0.0 | 0.0 | 100.0 | 0.0 | 0.0 | 0.0 | 0.0 | 0.0 | 0.0 |
| 139 | 0.0 | 0.0 | 0.0 | 0.0 | 0.0 | 50.0 | 0.0 | 0.0 | 0.0 | 0.0 | 50.0 | 0.0 |
| 114 | 0.0 | 0.0 | 0.0 | 0.0 | 0.0 | 50.0 | 0.0 | 0.0 | 0.0 | 0.0 | 50.0 | 0.0 |
| 29 | 0.0 | 0.0 | 0.0 | 0.0 | 0.0 | 0.0 | 100.0 | 0.0 | 0.0 | 0.0 | 0.0 | 0.0 |
| 168 | 0.0 | 0.0 | 0.0 | 0.0 | 0.0 | 0.0 | 50.0 | 0.0 | 0.0 | 0.0 | 50.0 | 0.0 |
| 46 | 0.0 | 0.0 | 0.0 | 0.0 | 0.0 | 0.0 | 50.0 | 0.0 | 0.0 | 0.0 | 50.0 | 0.0 |
| 88 | 0.0 | 0.0 | 0.0 | 0.0 | 0.0 | 0.0 | 0.0 | 50.0 | 0.0 | 0.0 | 50.0 | 0.0 |
| 71 | 0.0 | 0.0 | 0.0 | 0.0 | 0.0 | 0.0 | 0.0 | 50.0 | 0.0 | 0.0 | 50.0 | 0.0 |
| 129 | 0.0 | 0.0 | 0.0 | 0.0 | 0.0 | 0.0 | 0.0 | 0.0 | 50.0 | 50.0 | 0.0 | 0.0 |
| 48 | 0.0 | 0.0 | 0.0 | 0.0 | 0.0 | 0.0 | 0.0 | 0.0 | 50.0 | 0.0 | 50.0 | 0.0 |
| 155 | 0.0 | 0.0 | 0.0 | 0.0 | 0.0 | 0.0 | 0.0 | 0.0 | 50.0 | 0.0 | 50.0 | 0.0 |
| 163 | 0.0 | 0.0 | 0.0 | 0.0 | 0.0 | 0.0 | 0.0 | 0.0 | 0.0 | 100.0 | 0.0 | 0.0 |
| 21 | 0.0 | 0.0 | 0.0 | 0.0 | 0.0 | 0.0 | 0.0 | 0.0 | 0.0 | 0.0 | 100.0 | 0.0 |
| 22 | 0.0 | 0.0 | 0.0 | 0.0 | 0.0 | 0.0 | 0.0 | 0.0 | 0.0 | 0.0 | 100.0 | 0.0 |
| 37 | 0.0 | 0.0 | 0.0 | 0.0 | 0.0 | 0.0 | 0.0 | 0.0 | 0.0 | 0.0 | 100.0 | 0.0 |
| 50 | 0.0 | 0.0 | 0.0 | 0.0 | 0.0 | 0.0 | 0.0 | 0.0 | 0.0 | 0.0 | 100.0 | 0.0 |
| 55 | 0.0 | 0.0 | 0.0 | 0.0 | 0.0 | 0.0 | 0.0 | 0.0 | 0.0 | 0.0 | 100.0 | 0.0 |
| 66 | 0.0 | 0.0 | 0.0 | 0.0 | 0.0 | 0.0 | 0.0 | 0.0 | 0.0 | 0.0 | 100.0 | 0.0 |
| 73 | 0.0 | 0.0 | 0.0 | 0.0 | 0.0 | 0.0 | 0.0 | 0.0 | 0.0 | 0.0 | 100.0 | 0.0 |
| 122 | 0.0 | 0.0 | 0.0 | 0.0 | 0.0 | 0.0 | 0.0 | 0.0 | 0.0 | 0.0 | 100.0 | 0.0 |
| 135 | 0.0 | 0.0 | 0.0 | 0.0 | 0.0 | 0.0 | 0.0 | 0.0 | 0.0 | 0.0 | 100.0 | 0.0 |
| 138 | 0.0 | 0.0 | 0.0 | 0.0 | 0.0 | 0.0 | 0.0 | 0.0 | 0.0 | 0.0 | 100.0 | 0.0 |
| 160 | 0.0 | 0.0 | 0.0 | 0.0 | 0.0 | 0.0 | 0.0 | 0.0 | 0.0 | 0.0 | 100.0 | 0.0 |
| 162 | 0.0 | 0.0 | 0.0 | 0.0 | 0.0 | 0.0 | 0.0 | 0.0 | 0.0 | 0.0 | 100.0 | 0.0 |ATP7B No.2
#### Chart
| Category | WT | 2塩基欠失 | 13塩基欠失 | 1塩基挿入 | 9塩基欠失-1 | 8塩基欠失 | 16塩基欠失 | 9塩基欠失-2 | 20塩基欠失 | 11塩基欠失 | NHEJ.others | HDR |
|---|---|---|---|---|---|---|---|---|---|---|---|---|
| 202 | 100.0 | 0.0 | 0.0 | 0.0 | 0.0 | 0.0 | 0.0 | 0.0 | 0.0 | 0.0 | 0.0 | 0.0 |
| 235 | 100.0 | 0.0 | 0.0 | 0.0 | 0.0 | 0.0 | 0.0 | 0.0 | 0.0 | 0.0 | 0.0 | 0.0 |
| 239 | 100.0 | 0.0 | 0.0 | 0.0 | 0.0 | 0.0 | 0.0 | 0.0 | 0.0 | 0.0 | 0.0 | 0.0 |
| 264 | 100.0 | 0.0 | 0.0 | 0.0 | 0.0 | 0.0 | 0.0 | 0.0 | 0.0 | 0.0 | 0.0 | 0.0 |
| 266 | 100.0 | 0.0 | 0.0 | 0.0 | 0.0 | 0.0 | 0.0 | 0.0 | 0.0 | 0.0 | 0.0 | 0.0 |
| 238 | 100.0 | 0.0 | 0.0 | 0.0 | 0.0 | 0.0 | 0.0 | 0.0 | 0.0 | 0.0 | 0.0 | 0.0 |
| 230 | 100.0 | 0.0 | 0.0 | 0.0 | 0.0 | 0.0 | 0.0 | 0.0 | 0.0 | 0.0 | 0.0 | 0.0 |
| 211 | 100.0 | 0.0 | 0.0 | 0.0 | 0.0 | 0.0 | 0.0 | 0.0 | 0.0 | 0.0 | 0.0 | 0.0 |
| 234 | 100.0 | 0.0 | 0.0 | 0.0 | 0.0 | 0.0 | 0.0 | 0.0 | 0.0 | 0.0 | 0.0 | 0.0 |
| 250 | 100.0 | 0.0 | 0.0 | 0.0 | 0.0 | 0.0 | 0.0 | 0.0 | 0.0 | 0.0 | 0.0 | 0.0 |
| 243 | 100.0 | 0.0 | 0.0 | 0.0 | 0.0 | 0.0 | 0.0 | 0.0 | 0.0 | 0.0 | 0.0 | 0.0 |
| 218 | 100.0 | 0.0 | 0.0 | 0.0 | 0.0 | 0.0 | 0.0 | 0.0 | 0.0 | 0.0 | 0.0 | 0.0 |
| 201 | 100.0 | 0.0 | 0.0 | 0.0 | 0.0 | 0.0 | 0.0 | 0.0 | 0.0 | 0.0 | 0.0 | 0.0 |
| 237 | 100.0 | 0.0 | 0.0 | 0.0 | 0.0 | 0.0 | 0.0 | 0.0 | 0.0 | 0.0 | 0.0 | 0.0 |
| 242 | 50.0 | 50.0 | 0.0 | 0.0 | 0.0 | 0.0 | 0.0 | 0.0 | 0.0 | 0.0 | 0.0 | 0.0 |
| 254 | 50.0 | 0.0 | 0.0 | 50.0 | 0.0 | 0.0 | 0.0 | 0.0 | 0.0 | 0.0 | 0.0 | 0.0 |
| 275 | 50.0 | 0.0 | 0.0 | 50.0 | 0.0 | 0.0 | 0.0 | 0.0 | 0.0 | 0.0 | 0.0 | 0.0 |
| 209 | 50.0 | 0.0 | 0.0 | 0.0 | 0.0 | 0.0 | 0.0 | 0.0 | 0.0 | 0.0 | 50.0 | 0.0 |
| 217 | 0.0 | 100.0 | 0.0 | 0.0 | 0.0 | 0.0 | 0.0 | 0.0 | 0.0 | 0.0 | 0.0 | 0.0 |
| 185 | 0.0 | 100.0 | 0.0 | 0.0 | 0.0 | 0.0 | 0.0 | 0.0 | 0.0 | 0.0 | 0.0 | 0.0 |
| 228 | 0.0 | 100.0 | 0.0 | 0.0 | 0.0 | 0.0 | 0.0 | 0.0 | 0.0 | 0.0 | 0.0 | 0.0 |
| 229 | 0.0 | 50.0 | 50.0 | 0.0 | 0.0 | 0.0 | 0.0 | 0.0 | 0.0 | 0.0 | 0.0 | 0.0 |
| 260 | 0.0 | 50.0 | 50.0 | 0.0 | 0.0 | 0.0 | 0.0 | 0.0 | 0.0 | 0.0 | 0.0 | 0.0 |
| 223 | 0.0 | 50.0 | 50.0 | 0.0 | 0.0 | 0.0 | 0.0 | 0.0 | 0.0 | 0.0 | 0.0 | 0.0 |
| 231 | 0.0 | 50.0 | 0.0 | 0.0 | 50.0 | 0.0 | 0.0 | 0.0 | 0.0 | 0.0 | 0.0 | 0.0 |
| 222 | 0.0 | 50.0 | 0.0 | 0.0 | 50.0 | 0.0 | 0.0 | 0.0 | 0.0 | 0.0 | 0.0 | 0.0 |
| 190 | 0.0 | 50.0 | 0.0 | 0.0 | 50.0 | 0.0 | 0.0 | 0.0 | 0.0 | 0.0 | 0.0 | 0.0 |
| 193 | 0.0 | 50.0 | 0.0 | 0.0 | 0.0 | 50.0 | 0.0 | 0.0 | 0.0 | 0.0 | 0.0 | 0.0 |
| 262 | 0.0 | 50.0 | 0.0 | 0.0 | 0.0 | 50.0 | 0.0 | 0.0 | 0.0 | 0.0 | 0.0 | 0.0 |
| 258 | 0.0 | 50.0 | 0.0 | 0.0 | 0.0 | 0.0 | 0.0 | 0.0 | 0.0 | 0.0 | 50.0 | 0.0 |
| 240 | 0.0 | 50.0 | 0.0 | 0.0 | 0.0 | 0.0 | 0.0 | 0.0 | 0.0 | 0.0 | 50.0 | 0.0 |
| 282 | 0.0 | 50.0 | 0.0 | 0.0 | 0.0 | 0.0 | 0.0 | 0.0 | 0.0 | 0.0 | 50.0 | 0.0 |
| 189 | 0.0 | 50.0 | 0.0 | 0.0 | 0.0 | 0.0 | 0.0 | 0.0 | 0.0 | 0.0 | 50.0 | 0.0 |
| 198 | 0.0 | 50.0 | 0.0 | 0.0 | 0.0 | 0.0 | 0.0 | 0.0 | 0.0 | 0.0 | 50.0 | 0.0 |
| 280 | 0.0 | 50.0 | 0.0 | 0.0 | 0.0 | 0.0 | 0.0 | 0.0 | 0.0 | 0.0 | 50.0 | 0.0 |
| 257 | 0.0 | 50.0 | 0.0 | 0.0 | 0.0 | 0.0 | 0.0 | 0.0 | 0.0 | 0.0 | 50.0 | 0.0 |
| 276 | 0.0 | 50.0 | 0.0 | 0.0 | 0.0 | 0.0 | 0.0 | 0.0 | 0.0 | 0.0 | 50.0 | 0.0 |
| 186 | 0.0 | 0.0 | 100.0 | 0.0 | 0.0 | 0.0 | 0.0 | 0.0 | 0.0 | 0.0 | 0.0 | 0.0 |
| 192 | 0.0 | 0.0 | 100.0 | 0.0 | 0.0 | 0.0 | 0.0 | 0.0 | 0.0 | 0.0 | 0.0 | 0.0 |
| 274 | 0.0 | 0.0 | 50.0 | 50.0 | 0.0 | 0.0 | 0.0 | 0.0 | 0.0 | 0.0 | 0.0 | 0.0 |
| 208 | 0.0 | 0.0 | 50.0 | 0.0 | 0.0 | 0.0 | 0.0 | 0.0 | 0.0 | 0.0 | 50.0 | 0.0 |
| 263 | 0.0 | 0.0 | 50.0 | 0.0 | 0.0 | 0.0 | 0.0 | 0.0 | 0.0 | 0.0 | 0.0 | 50.0 |
| 221 | 0.0 | 0.0 | 0.0 | 100.0 | 0.0 | 0.0 | 0.0 | 0.0 | 0.0 | 0.0 | 0.0 | 0.0 |
| 182 | 0.0 | 0.0 | 0.0 | 50.0 | 0.0 | 0.0 | 0.0 | 50.0 | 0.0 | 0.0 | 0.0 | 0.0 |
| 227 | 0.0 | 0.0 | 0.0 | 50.0 | 0.0 | 0.0 | 0.0 | 0.0 | 0.0 | 0.0 | 50.0 | 0.0 |
| 284 | 0.0 | 0.0 | 0.0 | 50.0 | 0.0 | 0.0 | 0.0 | 0.0 | 0.0 | 0.0 | 50.0 | 0.0 |
| 261 | 0.0 | 0.0 | 0.0 | 50.0 | 0.0 | 0.0 | 0.0 | 0.0 | 0.0 | 0.0 | 50.0 | 0.0 |
| 233 | 0.0 | 0.0 | 0.0 | 0.0 | 0.0 | 50.0 | 0.0 | 0.0 | 0.0 | 0.0 | 0.0 | 50.0 |
| 253 | 0.0 | 0.0 | 0.0 | 0.0 | 0.0 | 0.0 | 50.0 | 0.0 | 0.0 | 0.0 | 50.0 | 0.0 |
| 259 | 0.0 | 0.0 | 0.0 | 0.0 | 0.0 | 0.0 | 0.0 | 0.0 | 0.0 | 0.0 | 100.0 | 0.0 |
| 220 | 0.0 | 0.0 | 0.0 | 0.0 | 0.0 | 0.0 | 0.0 | 0.0 | 0.0 | 0.0 | 100.0 | 0.0 |
| 212 | 0.0 | 0.0 | 0.0 | 0.0 | 0.0 | 0.0 | 0.0 | 0.0 | 0.0 | 0.0 | 100.0 | 0.0 |
| 279 | 0.0 | 0.0 | 0.0 | 0.0 | 0.0 | 0.0 | 0.0 | 0.0 | 0.0 | 0.0 | 100.0 | 0.0 |
| 184 | 0.0 | 0.0 | 0.0 | 0.0 | 0.0 | 0.0 | 0.0 | 0.0 | 0.0 | 0.0 | 100.0 | 0.0 |
| 187 | 0.0 | 0.0 | 0.0 | 0.0 | 0.0 | 0.0 | 0.0 | 0.0 | 0.0 | 0.0 | 100.0 | 0.0 |
| 197 | 0.0 | 0.0 | 0.0 | 0.0 | 0.0 | 0.0 | 0.0 | 0.0 | 0.0 | 0.0 | 100.0 | 0.0 |
| 241 | 0.0 | 0.0 | 0.0 | 0.0 | 0.0 | 0.0 | 0.0 | 0.0 | 0.0 | 0.0 | 100.0 | 0.0 |
| 245 | 0.0 | 0.0 | 0.0 | 0.0 | 0.0 | 0.0 | 0.0 | 0.0 | 0.0 | 0.0 | 100.0 | 0.0 |ATP7B No.3
#### Chart
| Category | WT | 2塩基欠失 | 13塩基欠失 | 1塩基挿入 | 9塩基欠失-1 | 8塩基欠失 | 16塩基欠失 | 9塩基欠失-2 | 20塩基欠失 | 11塩基欠失 | NHEJ.others | HDR |
|---|---|---|---|---|---|---|---|---|---|---|---|---|
| 288 | 100.0 | 0.0 | 0.0 | 0.0 | 0.0 | 0.0 | 0.0 | 0.0 | 0.0 | 0.0 | 0.0 | 0.0 |
| 292 | 100.0 | 0.0 | 0.0 | 0.0 | 0.0 | 0.0 | 0.0 | 0.0 | 0.0 | 0.0 | 0.0 | 0.0 |
| 294 | 100.0 | 0.0 | 0.0 | 0.0 | 0.0 | 0.0 | 0.0 | 0.0 | 0.0 | 0.0 | 0.0 | 0.0 |
| 295 | 100.0 | 0.0 | 0.0 | 0.0 | 0.0 | 0.0 | 0.0 | 0.0 | 0.0 | 0.0 | 0.0 | 0.0 |
| 300 | 100.0 | 0.0 | 0.0 | 0.0 | 0.0 | 0.0 | 0.0 | 0.0 | 0.0 | 0.0 | 0.0 | 0.0 |
| 301 | 100.0 | 0.0 | 0.0 | 0.0 | 0.0 | 0.0 | 0.0 | 0.0 | 0.0 | 0.0 | 0.0 | 0.0 |
| 307 | 100.0 | 0.0 | 0.0 | 0.0 | 0.0 | 0.0 | 0.0 | 0.0 | 0.0 | 0.0 | 0.0 | 0.0 |
| 316 | 100.0 | 0.0 | 0.0 | 0.0 | 0.0 | 0.0 | 0.0 | 0.0 | 0.0 | 0.0 | 0.0 | 0.0 |
| 338 | 100.0 | 0.0 | 0.0 | 0.0 | 0.0 | 0.0 | 0.0 | 0.0 | 0.0 | 0.0 | 0.0 | 0.0 |
| 341 | 100.0 | 0.0 | 0.0 | 0.0 | 0.0 | 0.0 | 0.0 | 0.0 | 0.0 | 0.0 | 0.0 | 0.0 |
| 342 | 100.0 | 0.0 | 0.0 | 0.0 | 0.0 | 0.0 | 0.0 | 0.0 | 0.0 | 0.0 | 0.0 | 0.0 |
| 348 | 100.0 | 0.0 | 0.0 | 0.0 | 0.0 | 0.0 | 0.0 | 0.0 | 0.0 | 0.0 | 0.0 | 0.0 |
| 350 | 100.0 | 0.0 | 0.0 | 0.0 | 0.0 | 0.0 | 0.0 | 0.0 | 0.0 | 0.0 | 0.0 | 0.0 |
| 359 | 100.0 | 0.0 | 0.0 | 0.0 | 0.0 | 0.0 | 0.0 | 0.0 | 0.0 | 0.0 | 0.0 | 0.0 |
| 309 | 50.0 | 50.0 | 0.0 | 0.0 | 0.0 | 0.0 | 0.0 | 0.0 | 0.0 | 0.0 | 0.0 | 0.0 |
| 315 | 50.0 | 50.0 | 0.0 | 0.0 | 0.0 | 0.0 | 0.0 | 0.0 | 0.0 | 0.0 | 0.0 | 0.0 |
| 320 | 50.0 | 0.0 | 0.0 | 0.0 | 50.0 | 0.0 | 0.0 | 0.0 | 0.0 | 0.0 | 0.0 | 0.0 |
| 354 | 50.0 | 0.0 | 0.0 | 0.0 | 0.0 | 0.0 | 0.0 | 0.0 | 0.0 | 0.0 | 50.0 | 0.0 |
| 302 | 0.0 | 100.0 | 0.0 | 0.0 | 0.0 | 0.0 | 0.0 | 0.0 | 0.0 | 0.0 | 0.0 | 0.0 |
| 332 | 0.0 | 50.0 | 0.0 | 50.0 | 0.0 | 0.0 | 0.0 | 0.0 | 0.0 | 0.0 | 0.0 | 0.0 |
| 289 | 0.0 | 50.0 | 0.0 | 0.0 | 0.0 | 0.0 | 0.0 | 0.0 | 0.0 | 50.0 | 0.0 | 0.0 |
| 291 | 0.0 | 50.0 | 0.0 | 0.0 | 0.0 | 0.0 | 0.0 | 0.0 | 0.0 | 50.0 | 0.0 | 0.0 |
| 298 | 0.0 | 0.0 | 100.0 | 0.0 | 0.0 | 0.0 | 0.0 | 0.0 | 0.0 | 0.0 | 0.0 | 0.0 |
| 317 | 0.0 | 0.0 | 50.0 | 0.0 | 0.0 | 0.0 | 0.0 | 0.0 | 0.0 | 0.0 | 50.0 | 0.0 |
| 358 | 0.0 | 0.0 | 0.0 | 100.0 | 0.0 | 0.0 | 0.0 | 0.0 | 0.0 | 0.0 | 0.0 | 0.0 |
| 303 | 0.0 | 0.0 | 0.0 | 50.0 | 0.0 | 0.0 | 0.0 | 0.0 | 0.0 | 0.0 | 50.0 | 0.0 |
| 326 | 0.0 | 0.0 | 0.0 | 50.0 | 0.0 | 0.0 | 0.0 | 0.0 | 0.0 | 0.0 | 50.0 | 0.0 |
| 335 | 0.0 | 0.0 | 0.0 | 50.0 | 0.0 | 0.0 | 0.0 | 0.0 | 0.0 | 0.0 | 50.0 | 0.0 |
| 362 | 0.0 | 0.0 | 0.0 | 50.0 | 0.0 | 0.0 | 0.0 | 0.0 | 0.0 | 0.0 | 50.0 | 0.0 |
| 329 | 0.0 | 0.0 | 0.0 | 0.0 | 100.0 | 0.0 | 0.0 | 0.0 | 0.0 | 0.0 | 0.0 | 0.0 |
| 287 | 0.0 | 0.0 | 0.0 | 0.0 | 50.0 | 0.0 | 0.0 | 0.0 | 50.0 | 0.0 | 0.0 | 0.0 |
| 355 | 0.0 | 0.0 | 0.0 | 0.0 | 50.0 | 0.0 | 0.0 | 0.0 | 0.0 | 0.0 | 50.0 | 0.0 |
| 296 | 0.0 | 0.0 | 0.0 | 0.0 | 0.0 | 50.0 | 0.0 | 0.0 | 0.0 | 0.0 | 50.0 | 0.0 |
| 299 | 0.0 | 0.0 | 0.0 | 0.0 | 0.0 | 0.0 | 0.0 | 50.0 | 0.0 | 0.0 | 50.0 | 0.0 |
| 318 | 0.0 | 0.0 | 0.0 | 0.0 | 0.0 | 0.0 | 0.0 | 0.0 | 0.0 | 0.0 | 100.0 | 0.0 |
| 321 | 0.0 | 0.0 | 0.0 | 0.0 | 0.0 | 0.0 | 0.0 | 0.0 | 0.0 | 0.0 | 100.0 | 0.0 |
| 357 | 0.0 | 0.0 | 0.0 | 0.0 | 0.0 | 0.0 | 0.0 | 0.0 | 0.0 | 0.0 | 100.0 | 0.0 |
| 361 | 0.0 | 0.0 | 0.0 | 0.0 | 0.0 | 0.0 | 0.0 | 0.0 | 0.0 | 0.0 | 100.0 | 0.0 |
| 293 | 0.0 | 0.0 | 0.0 | 0.0 | 0.0 | 0.0 | 0.0 | 0.0 | 0.0 | 0.0 | 0.0 | 100.0 |E
WT
Δ2-bp
Δ13-bp
1-bp ins
Δ9-bp_1
Δ8-bp
Δ16-bp
Δ9-bp_2
Δ20-bp_2
NHEJ.others
NHEJ+HDR
HDR
Model for ATP7B No.1
#### Chart
| Category | WT | Δ2-bp | Δ13-bp | Δ1-bp_1 | Δ9-bp_1 | Δ8-bp | Δ16-bp | Δ9-bp_2 | Δ20-bp | Δ11-bp | NHEJ.others | HDR |
|---|---|---|---|---|---|---|---|---|---|---|---|---|Model for ATP7B No.2
#### Chart
| Category | WT | Δ2-bp | Δ13-bp | Δ1-bp_1 | Δ9-bp_1 | Δ8-bp | Δ16-bp | Δ9-bp_2 | Δ20-bp | Δ11-bp | NHEJ.others | HDR |
|---|---|---|---|---|---|---|---|---|---|---|---|---|Model for ATP7B No.3
#### Chart
| Category | WT | Δ2-bp | Δ13-bp | Δ1-bp_1 | Δ9-bp_1 | Δ8-bp | Δ16-bp | Δ9-bp_2 | Δ20-bp | Δ11-bp | NHEJ.others | HDR |
|---|---|---|---|---|---|---|---|---|---|---|---|---|

### Slide 8
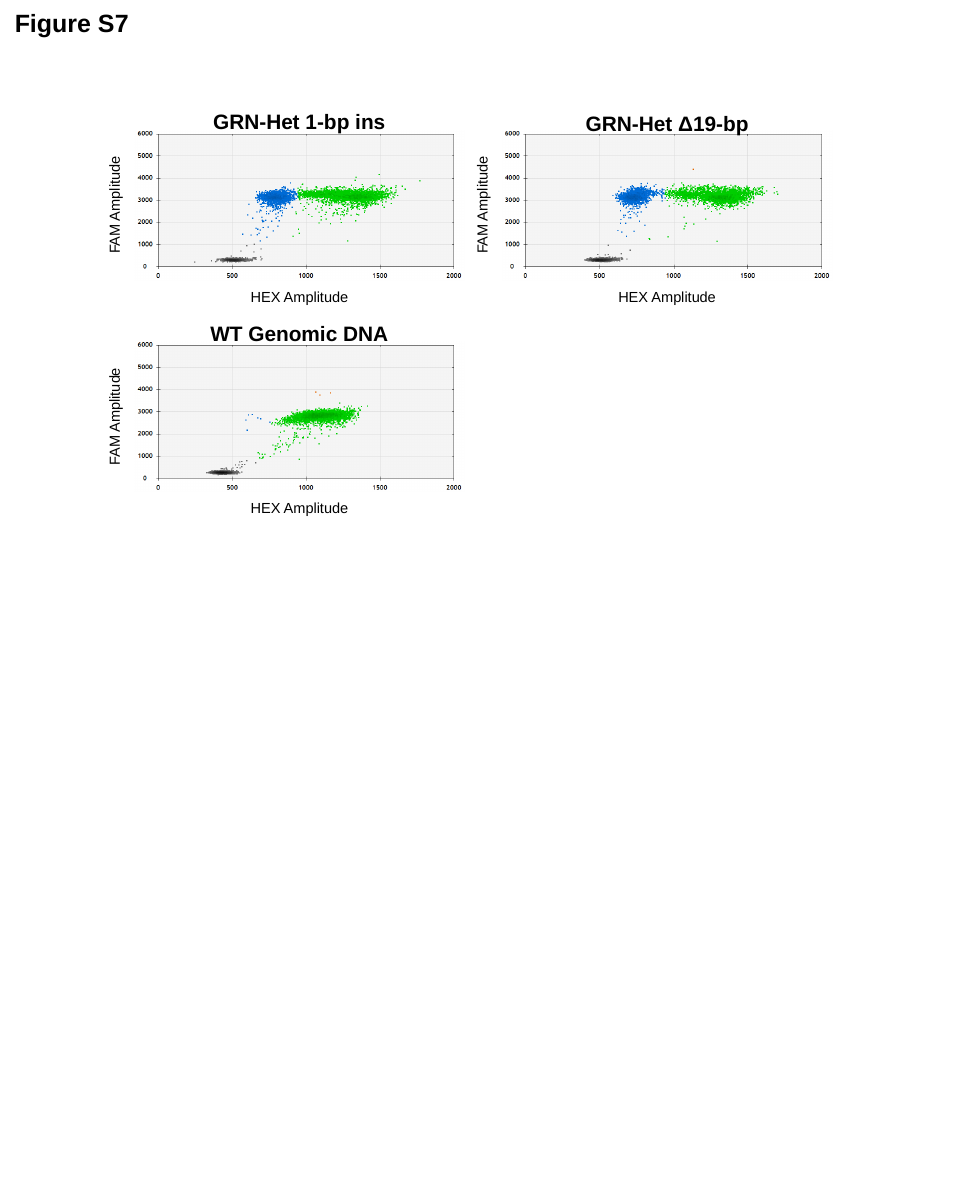

Figure S7
GRN-Het 1-bp ins
FAM Amplitude
HEX Amplitude
GRN-Het Δ19-bp
FAM Amplitude
HEX Amplitude
WT Genomic DNA
FAM Amplitude
HEX Amplitude
