## Supplemental Infomation for "High-throughput robotic isolation of human iPS cell clones reveals frequent homozygous induction of identical genetic manipulations by CRISPR-Cas9"

**Supplementary Figure Legends**

**Figure S1.**

**Characterization of iPS cell clones isolated by robotic picking.**

**(A)** Maintenance of pluripotency marker expression in iPS cell clones isolated by robotic picking. Immunocytochemistry of SOX2 and OCT4 in isolated iPS cell clones. The negative control (NTC) did not contain any primary antibodies. Scale bar: 100 μm. **(B)** Cloning efficiency of robotic picking of iPS cell clones for the 3 target genes. **(C)** iPS cells with mixed fluorescence before transfer into Matrigel domes to validate clonality. The 3 iPS cell lines expressing EGFP (green), mCherry (red), or EBFP (blue) were mixed. Scale bar: 100 μm.

**Figure S2.**

**Genotypes of genome-edited iPS cell clones.**

**(A and B)** Genome editing outcomes in isolated clones derived from single human iPS cells edited by Cas9. GRN **(A)** and ATP7B **(B)** editing outcomes are shown (No.1 to No.3). Each bar represents 1 clone, and the genotypes of WT (green), NHEJ (blue), HDR (red), and HDR + NHEJ (purple) in 1 clone are also shown in each bar. **(C and D)** Total allelic frequencies of WT (green), NHEJ (blue), HDR (red), and HDR + NHEJ (purple) in genome-edited iPS cells in the 3 experiments in GRN **(C)** and ATP7B **(D)** shown in Figure S2A and S2B, respectively.

**Figure S3.**

**Total allelic frequencies in genome-edited iPS cell pools before clone isolation.**

Amplicon sequencing of genome-edited iPS cell pools was used to determine the total allelic frequencies before clonal isolation.

**Figure S4.**

**Model diagrams of the distributions of clones with different genotypes.**

**(A)** Models assuming different alleles are randomly distributed at the observed frequencies for RBM20 No.1 and No.2. **(B)** Models for GRN No.1 to No.3. **(C)** Models for ATP7B No.1 to No.3.

**Figure S5.**

**Sequences and frequencies of alleles generated through GRN R493X and ATP7B R778L editing.**

**(A and B)** The 10 most frequently observed alleles after GRN **(A)** and ATP7B **(B)** editing. Black and red underlines indicate guide RNA and PAM sequences, respectively. Red triangles and red dotted lines indicate the site of cleavage by Cas9. Blue and red characters indicate unedited and substituted nucleotides, respectively.

**Figure S6.**

**Profiles and models of various indels induced by NHEJ in RBM20, GRN, and ATP7B**

**(A)** Model diagrams of the distributions of clones with different genotypes assuming different alleles are randomly distributed at the observed frequencies. Models for RBM20 No.1 and No.2 are shown. **(B)** Genome editing outcomes with distinguished NHEJ sequences in isolated clones derived from single human iPS cells edited by Cas9. GRN editing outcomes are shown (No.1 to No.3). **(C)** Models of the distributions of clones with different genotypes assuming different alleles are randomly distributed at the observed frequencies. Models for GRN No.1 to No.3 are shown. **(D)** Genome editing outcomes with distinguished NHEJ sequences in isolated clones. ATP7B editing outcomes are shown (No.1 to No.3). **(E)** Models of the distributions of clones for ATP7B No.1 to No.3 are shown.

**Figure S7.**

**Measurement of allelic frequencies by ddPCR in 2 GRN heterozygous knockout iPS cell lines.**

WT and NHEJ allelic frequencies of GRN-Het 1-bp and GRN-Het Δ19-bp lines were determined using ddPCR. The WT allele was detected as FAM+ HEX+ droplets (green) and the NHEJ alleles were detected as FAM-HEX + droplets (blue) in the 2D scatter plots.

**Supplementary Tables**

**Table S1. Parameters for picking cell clumps by CELL HANDLER**

| Parameter | Description |
| --- | --- |
| Major Diameter | Length of the major axis diameter of a cell clump.  Set range: 50-200 (μm) |
| Circularity | Value that represents how circular a cell clump is. A perfect circle is 1.0.  Set range: 0.3 - 0.95^a^ |
| Neighbor Distance | Distance between cell clumps, which is set to avoid accidental aspiration of neighboring cell clumps. Cell clumps with the largest distance between them are preferentially picked.  Set range: 60-360 (μm) |

^a^As air bubbles have values between 0.96 and 1.0, these are excluded.

**Table S2. Number of isolated iPS cell colonies with fluorescences**

| Exp. | No.1 | No.2 |
| --- | --- | --- |
| Wells with clones | 28 | 43 |
| EGFP | 17 | 27 |
| mCherry | 3 | 5 |
| EBFP | 1 | 2 |
| No Fluorescence | 7 | 9 |
| Mixed Fluorescence | 0 | 0 |

**Table S3. Sequences of oligonucleotide donor DNAs and gRNAs used in this study**

| Donor/gRNA | Name | Sequence (5’-3’) |
| --- | --- | --- |
| Donor | RBM20 R636S | ACAGATATGGCCCAGAAAGGCCGCGGTCT**A**GTAGTCCGGTGAGCCGGTCACTCTCCCCGA |
| Donor | GRN R493X | CGGCTGGCTACACCTGCAACGTGAAGGCT**T**GATCCTGCGAGAAGGAAGTGGTCTCTGCCC |
| Donor | ATP7B R778L | CATGCTCTTTGTGTTCATTGCCCTGGGCC**T**GTGGCTGGAACACTTGGCAAAGGTAACAGC |
| gRNA | RBM20 | GGTCT**C**GTAGTCCGGTGAGCCGG |
| gRNA | GRN | GAAGGCT**C**GATCCTGCGAGAAGG |
| gRNA | ATP7B | GGGCC**G**GTGGCTGGAACACTTGG |

PAM Sequences are underlined. Bold letters indicate the sites of single nucleotide substitutions.

**Table S4. Primary and secondary antibodies used in this study**

| Primary/Secondary | Target | Vendor and Cat# | Concentration |
| --- | --- | --- | --- |
| Primary | SOX2 | Abcam, ab97959 | 1:500 |
| Primary | OCT4 | Abcam, ab19857 | 1:500 |
| Secondary | Rabbit-IgG-568 | Invitrogen, A11011 | 1:2000 |

**Supplementary Videos**

**Video S1.** **Automated robotic iPS cell clump picking and seeding by CELL HANDLER**

First, imaging of cell clumps and tip loading was performed in parallel (0:00-0:18). Next, the loaded tips were corrected, and tip washing was conducted (0:19-0:39). Cell clumps selected by an image analysis were precisely picked and seeded into dispensing plates (0:40-1:12). Finally, the tips were discarded (1:13-1:22).
